# Emergent self-inhibition governs the landscape of stable states in complex ecosystems

**DOI:** 10.1101/2025.11.09.687513

**Authors:** Nitesh Kumar Patro, Washington Taylor, Akshit Goyal

## Abstract

Species-rich ecosystems often exhibit multiple stable states with distinct species compositions. Yet, the factors determining the likelihood of each state’s occurrence remain poorly understood. Here, we characterize and explain the landscape of stable states in the random Generalized Lotka–Volterra (GLV) model, in which multistability is widespread. We focus on the regime of the model where interspecies interactions are predominantly competitive and nearly symmetric. We find that the same pool of species with random initial abundances can result in different stable states, whose likelihoods typically differ by orders of magnitude. A state’s likelihood increases sharply with its total biomass, or inverse self-inhibition. We develop a simplified model to predict and explain this behavior, by coarse-graining ecological interactions so that each stable state behaves as a unit. In this setting, we can accurately predict the entire landscape of stable states using only two macroscopic properties: the biomass of each state and species diversity. Our analyses also provide insight into the biomass–likelihood relationship: High-biomass states have low self-inhibition and thus grow faster, outcompete others, and become much more likely. These results reveal emergent self-inhibition as a fundamental organizing principle for the attractor landscape of complex ecosystems—and provide a path to predict ecosystem outcomes without knowing microscopic interactions.

---

Species-rich ecosystems often exhibit multiple stable states with distinct species compositions and biomass levels [1]. Examples range from lush versus barren forests [2–4] to healthy versus diseased gut microbiomes [5, 6]. While multistability is well documented [7–13], the factors that govern which states are more *likely* across a variety of initial conditions remain poorly understood—limiting our ability to predict or control ecological transitions [14–17].

Understanding these likelihood patterns requires characterizing the sizes of attractor basins of different stable states. This becomes particularly challenging in diverse ecosystems, which are high-dimensional due to a large number of interacting species. In diverse ecosystems, attractor landscapes become complex and unintuitive [18], and classical low-dimensional analysis offers limited in-sight. Recent work using statistical physics tools has revealed rich phase behavior in disordered ecological models [19–32], including stability-to-chaos transitions [33– 38]. Yet, the multistable regime—where multiple attractors coexist—has received comparatively little attention. While previous studies have characterized the number of stable states [17, 39–42], we lack understanding of which factors predict the sizes of different attractor basins. Recent work on a certain class of “niche-structured” models [43] found that larger attractor basins correlate with increased steady-state biomass, suggesting this relationship may hold more broadly. However, there has been little work that describes the landscape of attractor basins for more general ecological models (though see [44]).

In this Letter, we analyze the multistability landscape of the random Generalized Lotka–Volterra (GLV) model with predominantly competitive, and nearly symmetric interspecies interactions. We show that in this model, a state’s likelihood increases sharply with its biomass in a strikingly consistent fashion, where biomass is computed as the sum of abundances of surviving species. We explain this relationship via a coarse-graining approach, in terms of an effective model in which individual stable states compete with each other. In this setting, each state behaves like a single unit governed by an emergent self-inhibition, which we find is inversely proportional to its steady-state biomass. This self-inhibition simultaneously slows growth and limits final abundance in opposing ways, creating a tradeoff that favors high-biomass states. We analytically derive the biomass–likelihood relationship for related monodominant and block-dominant situations, where only a single species or block of species survives in each distinct stable state, and use this to accurately predict the likelihood of 100s of stable states in a complex disordered ecosystem. Notably, this prediction can be made without precise details of the interaction matrix, which remains extremely hard to infer for highly-diverse ecosystems [45– 48]. Instead, it only requires macroscopic quantities— each state’s biomass (or more generally, net growth rate) and species diversity—which are easy to measure experimentally [49–51]. Thus our results provide a framework for estimating the relative likelihoods of alternative ecological states, without requiring exhaustive observational sampling or high-throughput experiments.

## Setup

Our starting point is the Generalized Lotka–Volterra (GLV) model, which describes the dynamics of *S* interacting species:

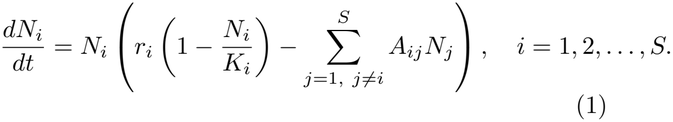

Here, *N*_*i*_ denotes the abundance (in units of biomass) of species *i, r*_*i*_ is its growth rate, *K*_*i*_ is its carrying capacity, and *A*_*ij*_ represents the effect of species *j* on species *i*. The matrix *A* thus encodes all interspecies interactions and determines the stability landscape of the ecosystem. Following the statistical physics tradition of replacing complexity with randomness, we assume the parameters *A*_*ij*_ are drawn as quenched random variables [19, 20, 23, 25, 33, 36, 52]. Specifically, we take *A*_*ij*_ to be distributed with mean *µ* and standard deviation *σ*. We consider the *strong interaction* regime: both *µ* and *σ* are finite constants that do not scale with *S*. This contrasts with some recent statistical physics treatments which focus on *weak interactions* where *µ* ∼ *O*(1*/S*) and *σ*^2^ ∼ *O*(1*/S*) [19, 20, 25, 33, 35], though there are exceptions [23, 36, 53]. Weak interactions are assumed only to ensure a proper thermodynamic limit as *S* → ∞ . Since natural ecosystems contain a finite number of species, and intuitively the interaction between a given pair of species does not change when a broader pool of species is available, we instead stick to strong interactions. We further assume that interactions are reciprocal, so *A* is symmetric, and we fix *r*_*i*_ = *K*_*i*_ = 1 for random matrices. Our results are qualitatively robust to relaxing these simplifying assumptions (Appendix C). Note that when *r*_*i*_’s differ by species, the growth-rate weighted biomass 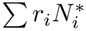 can be a slightly better predictor of likelihoods (see Appendix C for explanation).

By varying the two parameters *µ* and *σ*, we map out the phase diagram of the model (Fig. 1a), in agreement with past work [18, 20]. This diagram is dominated by a multistable phase (red), in which each community settles into one of multiple stable states depending on initial conditions. This region is separated from a unique stable state regime (white) by an analytically derived multistability boundary (black) using the cavity method (by appropriate rescaling of the boundary obtained for weak interactions [19]; Appendix J):

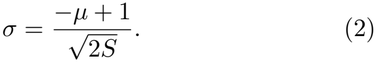

**FIG. 1:**
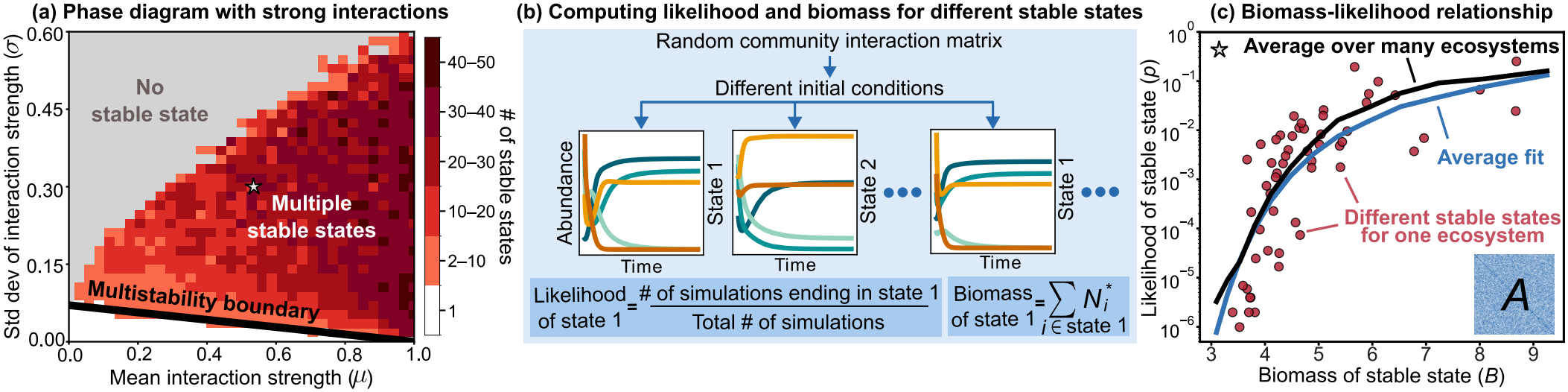
Multistability and biomass–likelihood relationship in the random GLV model with strong interactions. (a) Phase diagram showing the typical number of stable states for each interaction matrix *A*_*ij*_ with *S* = 100 species given its mean *µ* and standard deviation *σ*. Multistability is widespread (red region); black line shows the analytically-derived multistability boundary (Appendix J) separating the multistable and unique stable state (white) phase. (b) For a fixed *A*, varying initial conditions reveals different stable states. Shown are example dynamics (each species is a different color). Each state’s likelihood is estimated as the fraction of simulations that end in the state, while biomass is computed as the total abundance of surviving species. (c) The likelihood *p* of a stable state increases sharply with biomass *B*. Data in red show results from 10^6^ simulations with random initial conditions for a single matrix; black curve shows biomass-likelihood relationship averaged over 110 matrices with *µ* = 0.5, *σ* = 0.3. Blue curve shows fit to a hyperbolic relation: log(*p*) *∝* (*B*_*c*_ *− B*)^*−*1^ averaged over 110 matrices.

This shows that the multistable regime widens with increasing *µ*. Specifically, the critical *σ* at which multistability sets in decreases with *µ*, unlike for models with weak interactions, where it is constant [19]. The third phase (grey) represents a dynamically infeasible phase [19, 36] with unbounded growth and no stable states (Fig. 1a). Due to the prevalence of multistability, this model provides a natural setting to investigate the attractor landscape, including the relative likelihoods of stable states.

## Biomass–likelihood relationship

To probe the multistability landscape, we choose a random interaction matrix *A* with *S* = 100 species and simulate dynamics from 10^6^ random initial conditions. We sample each species’ initial abundance uniformly between 0 and 1, uncorrelated across species. Our central results are robust to other sampling schemes (Appendix C) and to the specific choice of matrix *A*. Each simulation converges to one of several possible stable states. We define the likelihood of a stable state as the fraction of initial conditions that converge to it (Fig. 1b, Appendix A). This allows us to numerically estimate ecological attractor basin sizes.

We find that the distribution of stable state likelihoods is highly skewed: the most likely state is over six orders of magnitude more probable than the least likely one observed (in many cases, there are further unobserved states that we might not detect; see Appendix A) . These dramatic differences in likelihood are strongly correlated with the total biomass of each state. Specifically, the likelihood of a stable state *p* increases sharply with its biomass *B*, with the sharpness progressively increasing at higher biomass (note the log scale in Fig. 1c). Averaging results from many random matrices, we obtain an approximation to the biomass-likelihood relationship (Fig. 1c; Appendix B), which roughly looks like a hyperbola in (−log *p, B*) space: log (*p*) ∝ (*B*_*c*_ −*B*)^*−*1^ (Fig. 1c, see Appendix K for a detailed discussion). The strong increase of the likelihood of a state with its biomass is robust to changes in the number of species, interaction parameters, carrying capacity variation, and degree of symmetry of *A* (Appendices C, K). These findings generalize earlier results on niche-structured ecosystems [43] (Appendix K for comparison), showing that biomass remains a strong predictor of the likelihood of different stable states in a broad range of complex ecosystems.

### Coarse-graining species to states

To understand the origin of this biomass–likelihood relationship, we focus on the interactions between species that coexist within the same stable state. Species that coexist in the same state inhibit each other much more weakly than expected by chance (Fig. 2a). Indeed, species in the same state can even be mutualistic and promote each other’s growth (*A*_*ij*_ *<* 0). This can be visualized as an emergent block structure in part of the interspecies interaction matrix (Figs. S26–S27, Appendix G). Together, these observations suggest that species within each state grow as a cohesive unit, motivating an effective model where different stable states compete with each other, and one state eventually emerges as the winner depending on initial conditions (Fig. 2b–c).

**FIG. 2:**
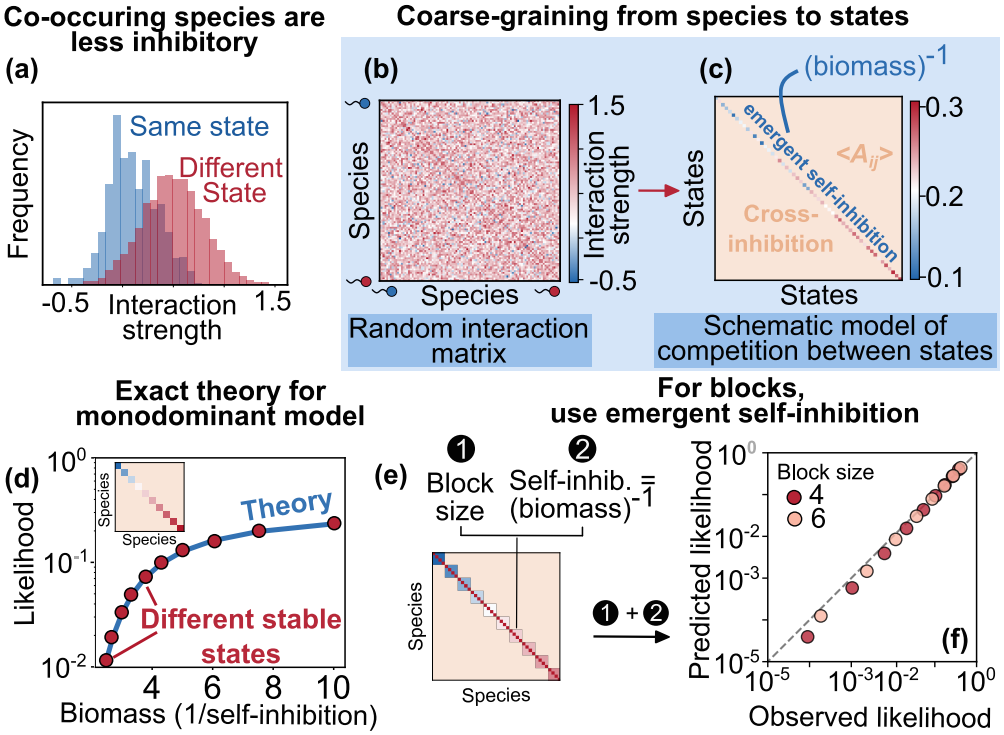
Coarse-graining states reveals emergent self-inhibition as a predictor of biomass and likelihood. (a) Species that interact weakly with each other form uninvadable stable states. The interactions among coexisting species are separated from the original interaction distribution, with a reduced mean. (b) Species-level interaction matrix. (c) We can conceptualize this as an effective matrix where different states compete with each other and one eventually emerges as the winner. (d) In a monodominant matrix where different stable states contain only a single species, both biomass and likelihood can be computed analytically from the self-inhibition (Appendix D). (e) In a block matrix where each different state corresponds to a block of *L* species, we can treat each block as an effective species with self-inhibition inversely proportional to its biomass. (f) Predictions combining this self-inhibition and block size accurately predict simulated likelihoods. Shown are block sizes *L* = {4, 6}.

We develop this picture in stages, starting with the simplest case and building toward random disordered matrices. We begin with a classic *monodominant* model [54] (also studied in this context in [43]), where individual species compete so strongly that only one species survives in each stable state. We then extend to block ecosystems with disjoint groups of coexisting species. Finally, we apply these results to random matrices, where we again think of states as blocks of species. Here, different states often overlap and contain species in common, which we can account for using a simple mean-field approximation under which the block model makes reasonable predictions even for random matrices (Appendix F). The monodominant model is exactly solvable. To construct it from Eq. (1), we consider a common interspecies interaction strength *A*_*ij*_ = *D* for all *i* ≠ *j* (cross-inhibition) and ensure that all self-inhibitions *A*_*ii*_ = *r*_*i*_*/K*_*i*_ are weaker than this: *A*_*ii*_ *< D* for all *i*. As in the random matrix case, here also we assume *r*_*i*_ = 1 (see Appendix C for generalization to variable growth rates *r*_*i*_). The per capita growth rate of each species then becomes

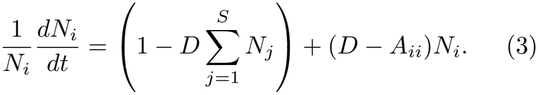

The first term in parentheses is common to all species, while the second is species-dependent. Specifically, the relative growth rate of each species depends on its abundance *N*_*i*_ dressed with a prefactor *χ*_*i*_ = *D*− *A*_*ii*_. If two species start at the same abundance, the one with larger *χ*_*i*_ will grow faster and win. In particular, we show that if all species start at abundances *N*_*i*_(0), the species with the largest value of the dressed initial condition *χ*_*i*_*N*_*i*_(0) will win (Fig. S14). For constant *D*, this condition implies that under uniform random initial conditions, species with lower self-inhibition *A*_*ii*_ will have a greater likelihood. Calculating this likelihood in general is rather challenging. With uniform initial conditions, the likelihood of each species winning is the fraction of the volume of initial conditions that maps to its attractor basin. For two species, calculating likelihoods is easy: a separatrix line partitions the two-dimensional phase space into two attractors. However, this method becomes computationally expensive when the number of species *S* is large. In this case, each pair of species defines a separatrix hyperplane, and the basin volumes correspond to *S*! high-dimensional solid cones subtended by the arrangement of these planes.

We circumvent this complexity by reformulating the problem as one of extreme value statistics. Rather than computing basin volumes directly, we consider the distributions of the dressed initial conditions *χ*_*i*_*N*_*i*_(0) and compute the probability that a given species has the maximum value. This simplification enables us to analytically calculate the likelihood *p*_*i*_ of species *i* winning in terms of *χ*_*i*_ (Appendix D) as:

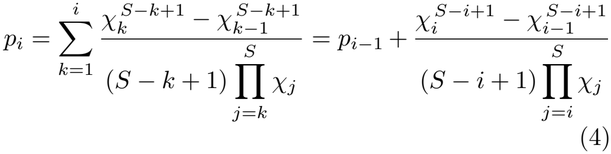

Here, species are indexed by *i* in increasing order of their *χ*_*i*_ (i.e., decreasing order of self-inhibition) and *χ*_0_ = 0. The recursive relation between *p*_*i*_ and *p*_*i−*1_ shows that decreasing self-inhibition (increasing *χ*) increases likelihood. Further, the dependence of *p*_*i*_ purely on *χ* highlights that self-inhibition *A*_*ii*_ is the key determinant of attractor likelihood in monodominant ecosystems. Notably, self-inhibition also directly sets biomass as *B*_*i*_ = 1*/A*_*ii*_. Thus, Eq.(4) can equivalently be expressed in terms of biomass *B*_*i*_ to reveal the biomass– likelihood relationship explicitly (Appendix D). Eq. (4) shows excellent agreement with simulations (Fig. 2d).

### Generalizing monodominant results to block ecosystems

The monodominant model predicts the likelihood of stable states containing only one species. In our random disordered ecosystems however, each state typically consists of several species. We thus extend our approach for computing state likelihoods to ecosystems with block-structured interaction matrices. We consider block ecosystems where each state *α* contains *L*_*α*_ species which inhibit each other with strength 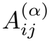 . Just like in the monodominant model, in a stable state one block outcompetes all others, but which block that will be might differ based on initial conditions. Species within each block *α* co-occur since their inhibition 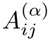 is weaker than inhibition from others outside the block *D* (Fig. 2e).

Similar to the monodominant model, to compute block likelihoods, we treat each block as an effective unit. In this context, we can again show that with a proper definition of effective self-inhibition (Appendix E), the block with the largest value of the dressed initial condition will win. The initial abundance for a block now involves summing the initial abundances of all *L*_*α*_ species within that block ∑_*i*∈*α*_ *N*_*i*_(0). By the central limit theorem, we expect this sum to be Gaussian-distributed for large *L*_*α*_. Further, each block’s initial abundance is dressed by a prefactor *χ*_*α*_ that depends on that block’s effective self-inhibition 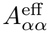 . Similar to the monodominant model, this quantity is the inverse of the block’s biomass *B*_*α*_. Combined, the dressed initial condition for block *α* becomes *χ*_*α*_ ∑_*i*∈*α*_ *N*_*i*_(0), where *χ*_*α*_ = *D −* 1*/B*_*α*_. This is well-approximated by a Gaussian with mean *m*_*α*_ = *L*_*α*_*χ*_*α*_*/*2 and variance 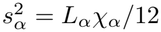(Appendix E). Following a similar calculation as the monodominant case, the likelihood *p*_*α*_ of block state *α* winning becomes:

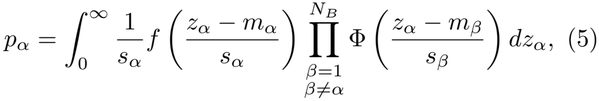

where *f* is the standard normal PDF, Φ is the standard normal CDF, and *N*_*B*_ is the total number of blocks. This integral has a rather simple interpretation—it computes the probability that block *α* has the largest dressed initial condition *χ*_*α*_ ∑_*i*∈*α*_ *N*_*i*_(0) among all blocks for a random uniform initial condition. To see this, notice that *f*_*α*_ gives the density of block *α*’s dressed initial condition being an arbitrary *z*_*α*_. The remaining product goes over all other blocks *β* ≠ *α* and demands that their dressed initial conditions be less than *z*_*α*_. We then simply integrate over all possible values of *z*_*α*_. Thus, Eq. (5) is really a generalization of the monodominant Eq. (4) to block states containing multiple species. This expression predicts the likelihoods observed in simulations extremely well for a range of block sizes *L*_*α*_ (Fig. 2e). Thus, for block-structured ecosystems, each state’s likelihood can be predicted using only the inter-state inhibition *D* and each state’s steady-state (equilibrium) biomass *B*_*α*_ and diversity *L*_*α*_.

### Predicting likelihoods for disordered ecosystems

Motivated by these results, we return to the case of random unstructured matrices from Fig. 1. We ask whether we can use the framework developed so far to predict state likelihoods in this case. Can we treat each observed state as an effective block which competes with other states? To map disordered matrices to the block model, we need to account for two major differences between these settings: (1) in disordered matrices, species often overlap between different states, unlike the block model with no overlaps; and (2) the inter-state inhibitions are now random variables. To this end, we generalized our block model to include random overlaps between blocks (Appendix H). We find that when overlaps are sufficiently widespread and randomly distributed, larger blocks tend to experience greater overall self-inhibition while smaller blocks tend to experience lower self-inhibition. This lowers the likelihood of larger blocks while increasing the likelihood of smaller blocks, thus reducing the effect of diversity variation between blocks (Fig. S28). This motivates the following mean-field approximation: we replace the size *L*_*α*_ of each block with the mean diversity ⟨*L*_*α*_⟩ = *L* across all *N*_*B*_ observed stable states, and the inter-block inhibition *D* with the mean interspecies interaction strength *µ* =⟨ *A*_*ij*_⟩ . For each block’s biomass *B*_*α*_, we use the biomass of each observed state.

Using these assumptions, our model accurately predicts the biomass–likelihood relationship in disordered interaction matrices (Fig. 3a). The predictions match simulations across a broad range of the multistable phase (Appendix I, Fig. S32). Indeed, prediction error decreases roughly as a power-law as we sample more matrices, supporting the accuracy of our theory (Fig. 3a, inset; Appendix B . Furthermore, for a particular ecosystem, we can predict the likelihood of each observed state using only its total biomass and typical species richness (Appendix I). These predictions align with simulated likelihoods for the majority of states (Fig. 3b), with some deviations at low-likelihoods. Together, these results demonstrate that the attractor landscape of complex random ecosystems is largely predictable without detailed knowledge of species-level interactions.

**FIG. 3:**
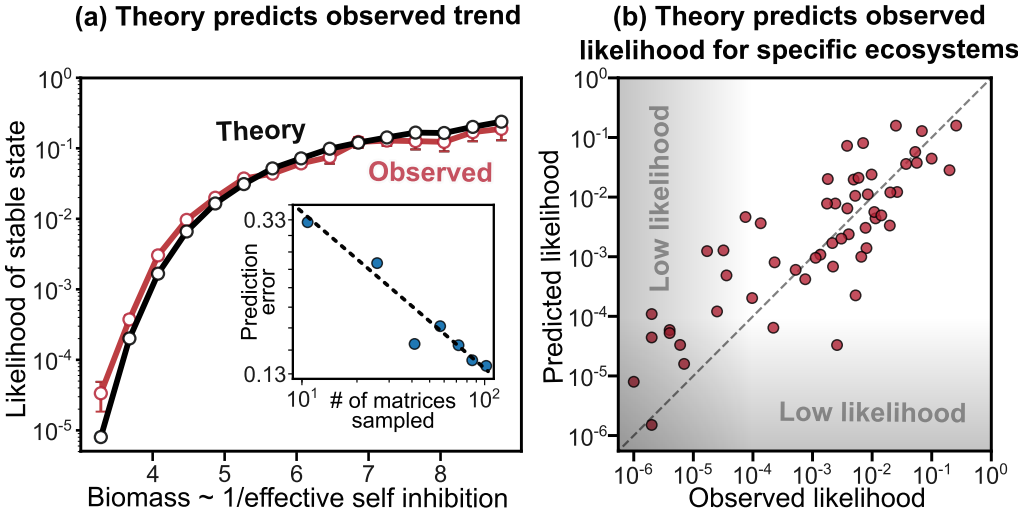
Biomass predicts state likelihoods in ecosystems with disordered interactions. (a) Using the biomass of each state as a measure of its emergent (inverse) self-inhibition, our theory (black) can predict the over-all biomass–likelihood relationship (red). Inset shows how prediction error decreases with number of sampled matrices (roughly power-law with exponent *−*0.4; Appendix B). For states with dramatically low likelihoods, our theory slightly deviates in likelihood predictions (gray region) (Appendix I). (b) For a specific ecosystem, the predicted likelihoods of each state, without any detailed knowledge of interactions, match the observed likelihoods for most states.

## Discussion

We have found a strong relationship between biomass and basin of attraction size for multiple stable states in a broad class of GLV models including monodominant, and block monodominant models, as well as random matrix models with predominantly competitive and nearly symmetric interactions, generalizing the observations of Ref. [43] for niche models to a much broader class of systems. Our findings furthermore give a quantitative underpinning to the idea that the likelihood of observing a set of species in a community is governed by how they collectively inhibit each other during growth. This collective self-inhibition of a community is a complex, emergent quantity that can be estimated as the inverse of the community’s final biomass. This leads to a robust positive correlation between the biomass and likelihood of a state. Note that here we focus on the relative probabilities of distinct stable states, giving the fundamental probability distribution on states. Since there are more states with intermediate biomass, computing properties of a “typical” state under this distribution would involve a balance of entropic factors favoring intermediate biomass and domain size, favoring larger biomass. Our results motivate experimental validation in natural microbial communities. In particular, communities derived from parallel enrichments—e.g., of soils, human microbiomes or plant rhizospheres—often give rise to multiple stable states under identical conditions [10–13, 55]. Quantifying the frequency of states across replicates enables one to estimate their likelihoods. Our results predict that these frequencies should be predictable only using each state’s total biomass (e.g., via cell counts or optical density) and species diversity (e.g., from 16S amplicon sequencing). Note that when individual species have different growth rates, slightly different measurements such as total community growth rate would be needed in our prediction method (Appendix C). We also note that our analysis applies directly to deterministic ecosystem dynamics and neglects stochastic fluctuations due to random births and deaths. While our results are robust to introducing such noise (Appendix C), it would be interesting to fully explore its consequences, especially how noise affects the interplay between the widths and depths of attractor basins. Another possible direction is predicting which species subsets form stable states directly from interactions rather than simulations, although this may be computationally difficult (Appendix K).

Beyond empirical validation, our findings open several directions for future work. Can we build on our framework to find ways to control ecosystems to be in states with low likelihoods, which might otherwise be healthy or functionally desirable? How can we predict the likelihood of being near a steady-state in chaotic ecological dynamics, which can occur due to nonreciprocal interactions [33, 35]? Does community evolution significantly deform the landscape of ecological stable states? Addressing these questions will be key to understanding how interspecies interactions collectively determine the state of a complex ecosystem.

## Acknowledgements

We thank J.P. O’Dwyer, A. Altieri, J. Gore, and S.Y. Li for valuable discussions. A.G. acknowledges support from the DAE under project no. RTI4001, as well as a Ramanujan Fellowship. The work of W.T. was supported in part by a grant from the Schmidt Futures Foundation, and W.T. would like to thank the Santa Fe Institute and the University of Auckland for hospitality while some of this work was done.

## Appendix A: Computing the likelihoods and biomass of stable states

In this appendix we describe our algorithm to compute likelihoods and biomasses for different uninvasible stable states corresponding to a chosen interaction matrix *A*. We first generate a random interaction matrix *A* with mean *µ* and standard deviation *σ*. Note that in most of the results, we use a symmetric interaction matrix, *A*_*ij*_ = *A*_*ji*_, but relax this assumption in Fig. S5. Symmetric interaction matrices yield a Lyapunov function for the dynamics in the GLV model [56, 57], given by

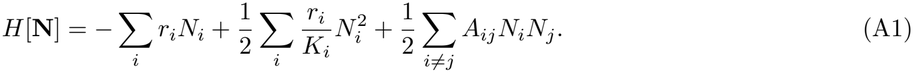

Here *N*_*i*_ is the abundance of species *i, r*_*i*_ is its intrinsic growth rate, *K*_*i*_ is its carrying capacity and *A*_*ij*_ represents the strength of the interaction between species *i* and *j*. The existence of this Lyapunov function guarantees that at long time, the dynamics always flow to a stable state, rather than having limit cycles or chaos [58]; this makes it very tractable to study multiple stable states. In this setting, each stable state is a local minima of this function, and this allows us to use the Lyapunov function to envision a landscape of stable states [22]. We also note that in what follows, we will not attempt to find every stable state corresponding to a given matrix *A*. This is because there are a substantially large number of states with exceedingly small basins of attraction, and while it is in many cases possible to identify these states (see Appendix F), statistically determining their likelihoods requires a computationally infeasible number of simulations when *A* is a large matrix (in our case, typically 100 *×* 100). We typically use 10^6^ initial conditions, while we estimate that computing likelihoods for all states might require roughly 10^10^ initial conditions (see Appendix F). Thus we analyze probabilities for a large but not complete subset of all the stable states; for example, we encounter 55 states in our simulations for the matrix in Fig. 1c while the total number of states is around 80 (Fig. S25).

To find stable states, we start by initializing all species with abundances sampled from a uniform distribution between 0 and 1. We then solve the GLV equations using the Radau method, which has adaptive time steps. After running the simulation for sufficient time until *t* = 300, we check whether we have reached a stable, uninvasible steady state in the following ways:

### Identifying surviving species

To determine steady state convergence, we first identify the surviving species. In our numerical simulations, species abundances never reach exactly zero, and species that are destined to go extinct may nevertheless take a long time to do so. To ascertain which species are on their way to extinction, we scan over extinction threshold abundances for surviving species over a wide range between 10^*−*5^ and 10^*−*1^. For most initial conditions, a threshold of 10^*−*5^ successfully identifies the surviving species. We begin by checking for feasibility. We start with a threshold of 10^*−*5^. We then classify all species with abundances above this threshold as surviving species. From the surviving species, we construct a reduced interaction matrix *A*_reduced_ that contains only the surviving species ; note that the diagonal elements of this matrix are *A*_*ii*_ = *r*_*i*_*/K*_*i*_. We then compute the steady-state abundances using the fixed point conditions of the GLV model in Eq. (1) as:

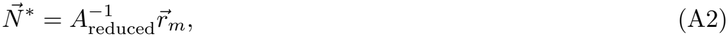

where *m* represents the number of surviving species and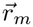 is the *m*-dimensional vector of growth rates of surviving species. For a given set of surviving species, the GLV equations have a unique fixed point, as given by Eq.(A2).

### Feasibility verification

If computing 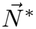 yields negative abundances for any species, then the set of surviving species is not feasible. If the set is feasible, then we ascertain if the abundances of survivors from simulations and Eq.(A2) are close enough (within numerical error). If we find that our guess of the surviving species is either infeasible or outside numerical error, then we assume that our extinction threshold was incorrect. In this case, we increase our threshold by a factor of 10 and repeat our procedure to identify surviving species. We do so until we reach a threshold abundance of 10^*−*1^. If we still fail to find a feasible steady state whose predicted abundances match our simulations, then we assume that we have not yet reached steady-state. In this case, we continue the simulation for additional time *t* = 1000 from this point, and repeat the process of identifying survivors after this additional time using the same procedure above. Once we find a feasible set of survivors with their steady-state abundances, we check their stability. We discard the small minority of simulations (approximately 0.1%) that fail to converge to a uninvasible steady state by *t* = 50300.

### Stability verification

To check stability, we calculate the Jacobian matrix at the steady state:

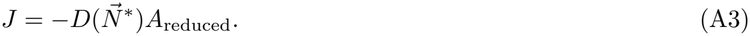

**FIG. S1:**
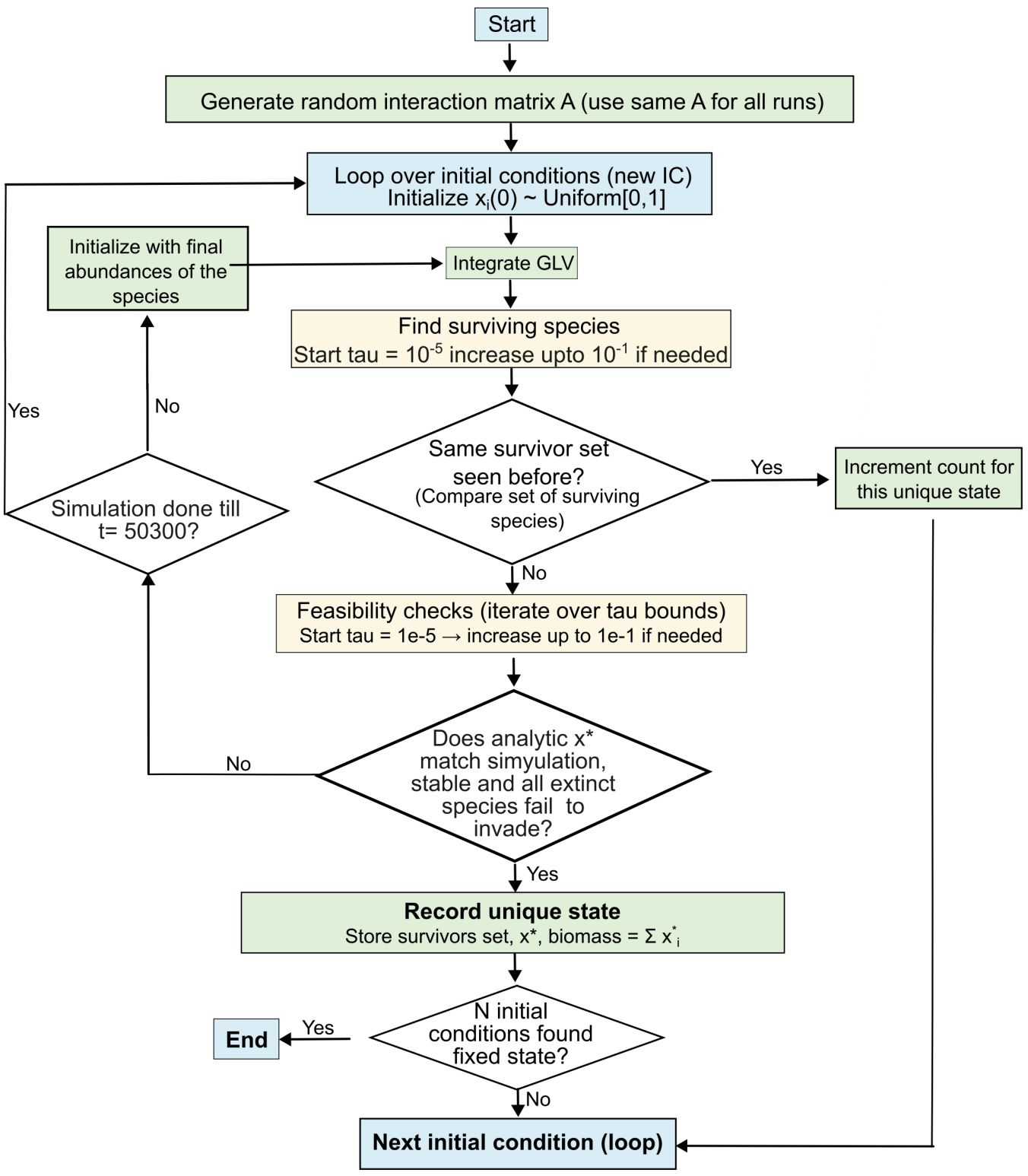
Flowchart of our algorithm to identify feasible, stable and uninvadable states for a given GLV interaction matrix in our model. Briefly, we choose a random initial condition, numerically integrate the dynamics, and detect surviving species using an self-consistently determined extinction threshold. We then finally ensure the resulting state is feasible and stable both to perturbations in surviving species abundances and to invasions of extinct species.

If the largest real part of all eigenvalues of *J* is negative, then the steady state is linearly stable to perturbations in abundances. We ensure that all steady states we obtain pass this test.

### Uninvasibility verification

We next check whether the stable steady state is uninvasible. Uninvasibility refers to the idea that extinct species should not be able to reinvade the community when introduced in small numbers. If *m* is the number of surviving species so that *S* − *m* species are extinct, then the state is uninvasible if all extinct species have negative invasion fitness, given by their per capita growth rate at low abundance:

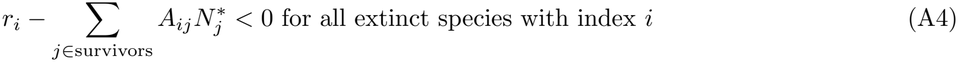

Once a steady-state passes even this criterion, it is certain to be a feasible, stable and uninvasible steady-state for the model with the given interaction matrix *A*. We add this state to our list of stable states.

### Cataloging unique states

We then start from a new randomly chosen initial condition, and repeat this entire process. After ascertaining which state this initial condition arrives at, we check whether the state is a new, unique state that we had not identified before, or if it matches an existing state. We track the number of random initial conditions that converge to each of these states. After repeating these simulations for 10^6^ random initial conditions, we collect all the unique stable states and the fraction of initial conditions that converge to them. Note that for convenience, we use 10^6^ initial conditions in the main text, but for several SI figures, we use 10^5^ initial conditions instead.

### Computing likelihoods and biomass

We define the likelihood of each unique stable state as:

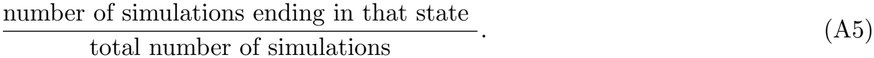

We calculate the biomass of each state as the sum of all steady-state species abundances for that state, i.e., 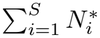 .

Using the Lyapunov function, the likelihood of a stable state is related to the size of its basin of attraction, while the biomass at steady-state is related to its depth [43, 58]. However, this Lyapunov function does not guarantee any clear relation between the size in abundance space and the depth of attractor basins. In this work, we find a robust positive biomass–likelihood relationship, showing that the size and depth of basins may indeed be related in the GLV model.

## Appendix B: Disorder averaging of the biomass–likelihood relationship

In the main text, we show that for a single random interaction matrix *A*, the likelihoods *p* of different stable states increase with biomass *B* and are well-described by our block-model. However, for any single realization of *A*, we observe error due to disorder in the interaction matrices, which are drawn randomly. To isolate the average biomass-likelihood relationship, we perform a quenched disorder average over many independent matrices drawn with the same (*µ, σ*). Here we use *N*_*I*_ = 110 independent matrices.

### Averaging protocol

For each disordered matrix *A*^(*k*)^ (*k* = 1, …, *N*_*I*_), we bin the stable states by biomass *B* and compute the mean likelihood ⟨*p* | *B* ∈*b, A*^(*k*)^ ⟩ within each bin *b* (see Appendix L for details of binning procedure). The disorder-averaged conditional likelihood is then

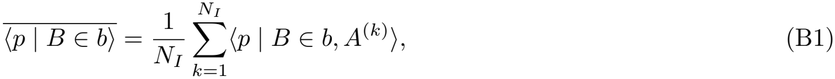

where

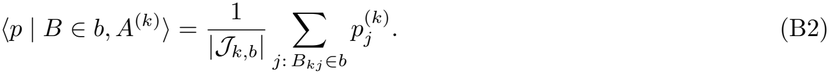

Here, |*J*_*k,b*_| is the observed number of states for matrix *A*^(*k*)^ whose biomass falls in bin *b*, and *j* indexes the specific stable states with biomass *B*_*kj*_ observed for matrix *k*. An alternative is to pool all data points across interaction matrix realizations and compute a single conditional average:

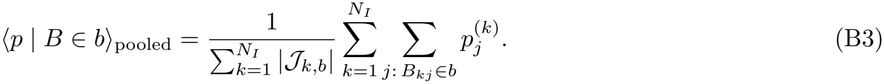

We find that both methods produce very similar results when the number of states per bin does not vary strongly across matrix interaction matrix realizations (Fig. S2), so we use the first method throughout.

**FIG. S2:**
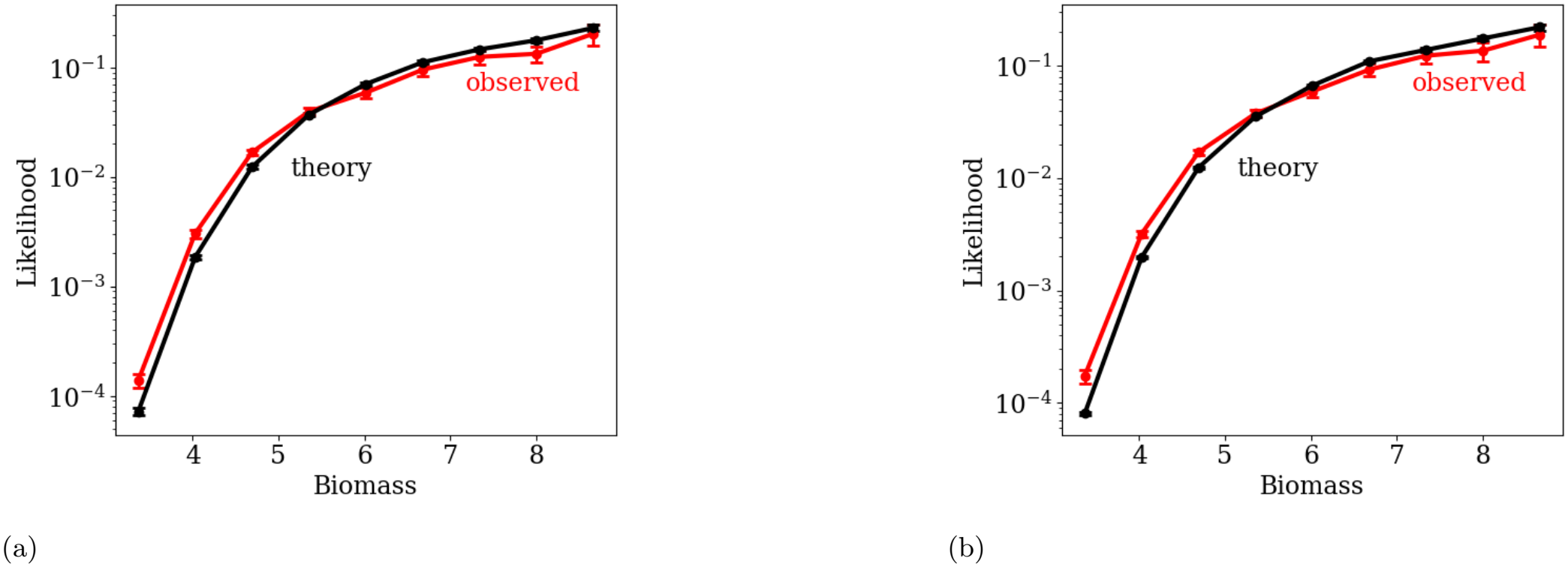
Comparison of averaging methods: (a) individual averaging, where the mean likelihood is first computed within each interaction matrix realization *A*^(*k*)^ for a given biomass bin and then averaged over matrices, and (b) pooled averaging, where all data points in each biomass bin are combined across interaction matrix realizations before averaging. Both methods produce the same mean trend and agree with the theoretical prediction, confirming that the result does not depend on the order of averaging. The error bars represent the standard error of the mean (SEM).

### Arithmetic vs. geometric mean

A subtlety arises in comparing disorder-averaged data with theory. Our block-model prediction yields the likelihood *p* directly, not log *p*. Since the scatter around the predicted value is approximately symmetric in *p* (not in log *p*), the appropriate comparison is between the theory and log ⟨*p*⟩, the logarithm of the arithmetic mean. Comparing instead with ⟨log *p*⟩ (the geometric mean) would overweight rare, low-likelihood states. In general, log⟨ *p*⟩ ≠ ⟨log *p*⟩ ; we use the arithmetic mean throughout this appendix.

### Comparison with theory

Fig. S3(a) shows the disorder-averaged likelihood ⟨*p*⟩ in each biomass bin, computed from *N*_*I*_ = 110 independent matrices with *µ* = 0.5, *σ* = 0.3, compared with the block-model prediction from Eq. (5) in the main text (using the mean block size as discussed in Appendix I). Error bars indicate the standard error of the mean, 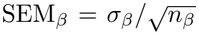, where *σ*_*β*_ is the standard deviation of likelihoods within biomass bin *β* and *n*_*β*_ is the number of contributing data points across all interaction matrix realizations. The averaged data closely follow the theoretical curve, with only minor deviations at the lowest biomass bins where states are rare and sampling is limited.

To quantify prediction error, we compute the root-mean-square-log error (RMSE_log_) between our theoretical prediction and observations from simulations across biomass bins:

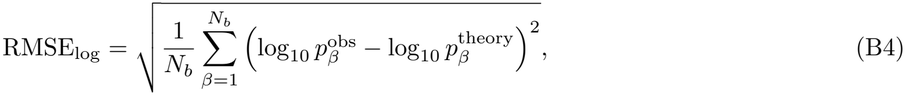

where *N*_*b*_ is the number of biomass bins and the index *β* labels bins. Here, 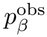 is the average likelihood obtained from simulations, while 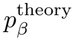 is the mean calculated likelihood across all states within that bin. This is sensitive to errors at low likelihoods. Since state likelihoods vary over orders of magnitude, one might prefer this as a metric of prediction error. We find that this error decrease monotonically as we sample more interaction matrices, i.e., as *N*_*I*_ increases (Fig. S3(c)). This confirms that our theory captures the average biomass-likelihood relationship rather accurately, and any scatter in likelihood estimates for a specific realization of an interaction matrix is likely a reflection of quenched disorder fluctuations.

**FIG. S3:**
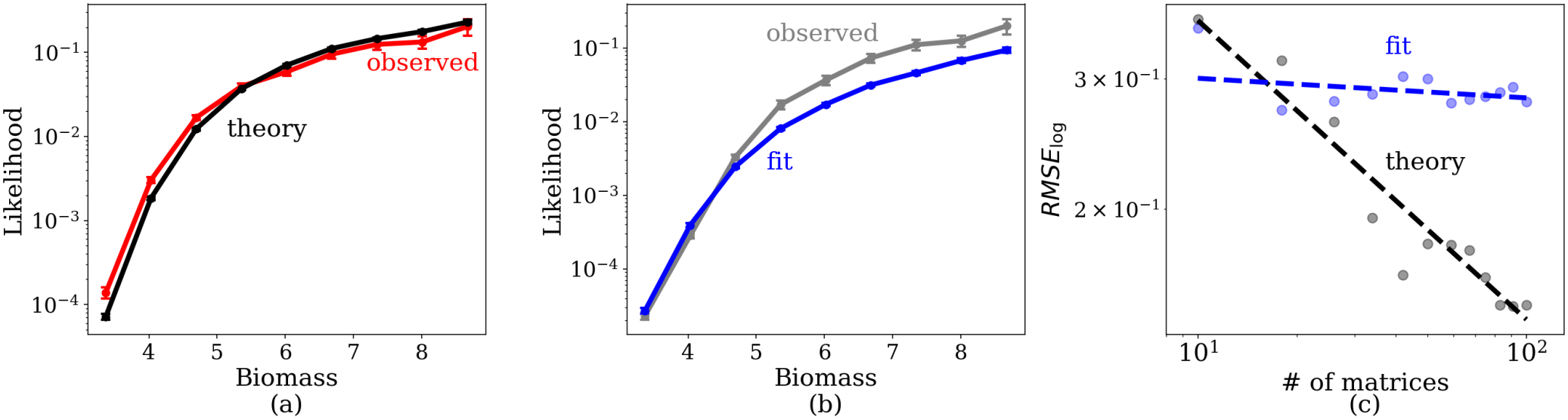
Our theory captures the biomass–likelihood relationship better than the log-hyperbolic fit. (a) The predicted block-model prediction (black) matches the disorder-averaged likelihood ⟨*p*⟩ (red) across biomass bins, with error bars showing the standard error of the mean. The theoretical curve captures the observed trend with only minor deviations at low-biomass. (b) The disorder-averaged log-hyperbolic fitted curve (blue) broadly follows the disorder-averaged log-likelihood log ⟨*p*⟩ (gray) across biomass bins, with error bars showing the standard error of the mean. The theoretical curve reproduces the overall trend, though some deviations are observed at high-biomass. (c) The RMSE_log_ as a function of number of matrices shows that block-model prediction (black dots) captures the trend more compared to the log-hyperbolic fit (blue dots), with error decreasing systematically faster as the number of matrices increases. The dashed lines are power-law fits to the data points, with exponent −0.4 (blue) and −0.03 (red). The disordered average is for the 110 matrices used in Fig. 1c.

### Comparison with log-hyperbolic fit

We can perform a similar disorder average for the log-hyperbolic fit log *p* ∝ (*B*_*c*_ − *B*)^*−*1^. Since this fit is performed in log-likelihood space, the appropriate comparison is between the averaged fit and log ⟨*p*⟩ . We find that the fit decently approximates the data at low likelihoods but deviates systematically for states with high likelihoods (Fig. S3(b)). Furthermore, in stark contrast with the prediction from our theory, the RMSE for the log-hyperbolic fit decreases very slowly compared to the theory, showing that our theory continues to predict better with increase in number of matrix realizations (Fig. S3(c)). This shows that the log-hyperbolic fit, while a simplified phenomenological approximation, does not capture the true biomass-likelihood relationship nearly as accurately as our theory.

### Visualizing the trend across many matrices

When all data points from *N*_*I*_ = 110 matrices are plotted together, the large spread of biomass values across interaction matrix realizations makes it difficult to discern the underlying trend by eye (Fig. S4a). To make the trend visible, we plot a kernel density estimate (KDE) of the likelihood within each biomass bin, normalizing the distribution separately in each bin. This visualization clearly reveals the predicted biomass–likelihood relationship even in the raw multi-matrix data (Fig. S4b).

**FIG. S4:**
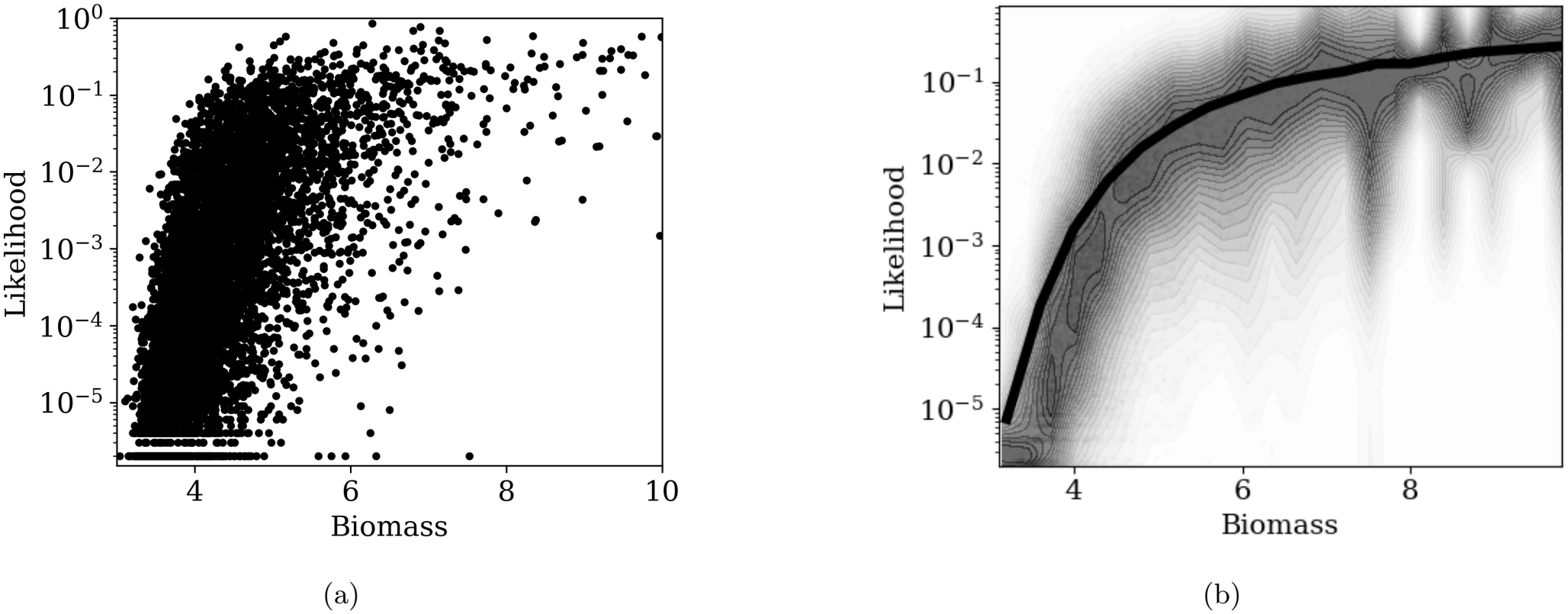
(a) Likelihood versus biomass for all stable states from *N*_*I*_ = 110 independent matrices at *µ* = 0.5, *σ* = 0.3. The large spread of biomass values across realizations obscures the underlying trend. (b) KDE of the likelihood in each biomass bin, with each bin normalized separately. The predicted biomass–likelihood trend is clearly visible in this representation. The black curve shows our theoretical prediction, also shown in 3(a)

## Appendix C: Robustness of biomass–likelihood relationship

In the main text, all our results have been shown for symmetric interaction matrices, fixed growth rates *r*_*i*_ = 1 and carrying capacities *K*_*i*_ = 1, number of species in the pool *S* = 100 and uniform, random initial conditions between 0 and 1. Further, we also assume deterministic dynamics with no demographic noise. Here we systematically relax these assumption and show that our central result—namely the biomass-likelihood relationsip—is qualitatively robust to including interaction asymmetry, allowing hetereogeneity in growth rates and carrying capacities, modifying the number of species in the pool, sampling initial species abundances from a truncated Gaussian, and including demographic noise in the dynamics. Towards the end of this Appendix, we also discuss the reliability and uncertainty in our likelihood estimates for different stable states.

First, we discuss interaction asymmetry. We repeat our analysis in Fig. 1c with a slightly asymmetric matrix, i.e., with across-diagonal correlation corr(*A*_*ij*_, *A*_*ji*_) = {0.9, 0.8 }. Just as in symmetric matrices, in these cases we also observe that states with higher biomass are much more likely (Fig. S5a–b).

In general, we expect that as long as asymmetries are not so big that the qualitative nature of the dynamics changes, such as by introducing limit cycles or chaos, asymmetries will be either neutral or enhance the tendency to favor states with lower self-inhibition. For example, in the block models studied in Appendix E, introducing an asymmetry in a matrix entry between blocks will favor one block over the other, but will not affect the biomass/self-inhibition of either block, which is determined solely by the in-block matrix elements, so such a change is neutral to the probability/biomass correlation. On the other hand, introducing a slight asymmetry of order *ϵ* in a given block has the effect of slightly increasing (by order *ϵ*^2^) the self-inhibition of the block and correspondingly decreasing the biomass. An analysis similar to that of the next Appendix shows that this correlates with a slight deviation of the separatrix from other blocks that disfavors the block with the added asymmetry; thus, in such a case the asymmetry causes same-sign changes in biomass and probability, maintaining the correlation between these quantities.

Next, instead of fixed carrying capacity *K*_*i*_ = 1 for all species, we introduce species-specific heterogeneity in *K*_*i*_ by sampling them from a uniform distribution on [1 −*ϵ*_*k*_, 1 + *ϵ*_*k*_]. Note that this change affects the self-inhibition *A*_*ii*_ = *r*_*i*_*/K*_*i*_ of each species. Thus, now both the self-inhibition *A*_*ii*_ and interspecies inhibition *A*_*ij*_ are disordered in the interaction matrix. Here *ϵ*_*k*_ is the measure of variation in the carrying capacities. In the main text, we study the case *ϵ*_*k*_ = 0. We observe that for *ϵ*_*k*_ ={ 0.01, 0.05 }, the biomass–likelihood relationship still shows a positive correlation and is qualitatively robust to species-specific *K*_*i*_ (Fig. S6).

**FIG. S5:**
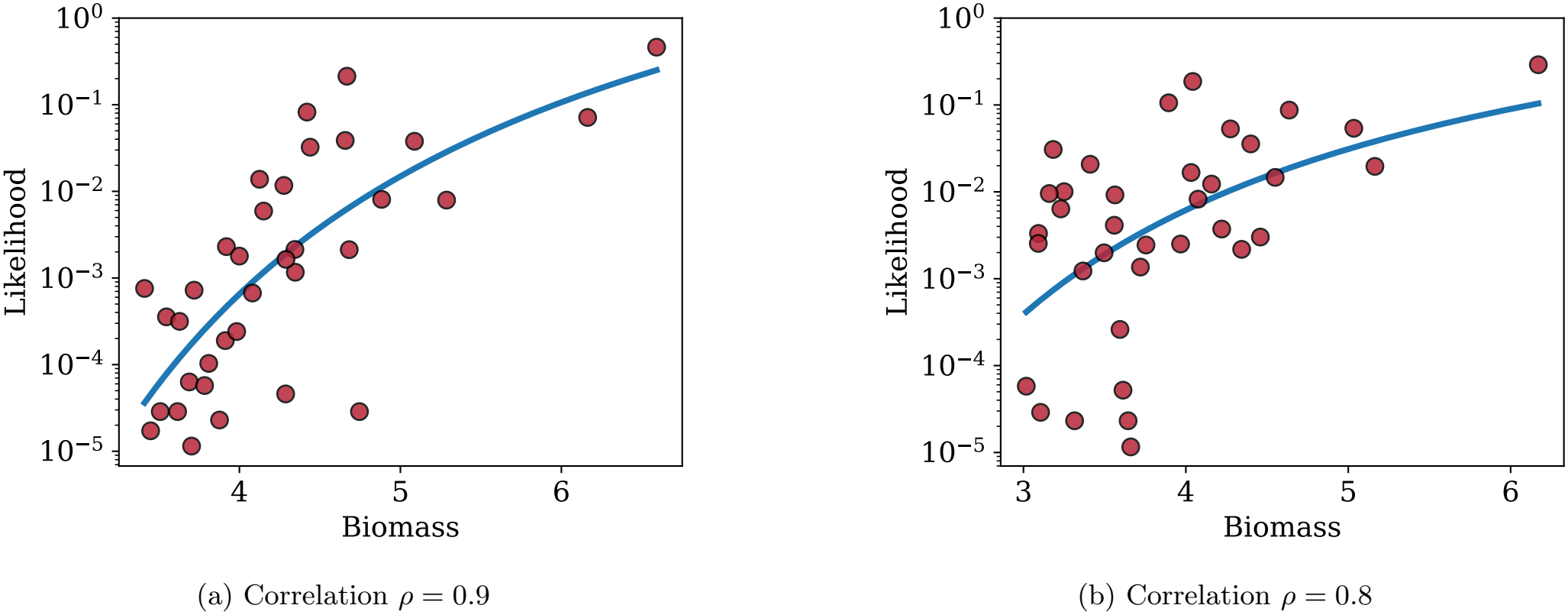
Biomass–likelihood distributions for asymmetric interaction matrices. (a) Correlation *ρ* = 0.9, (b) correlation *ρ* = 0.8. In both cases a positive correlation between likelihood and biomass is observed, although the strength decreases as *ρ* becomes smaller. Curves indicate fits to the log-hyperbolic relation log (*p*) ∝ (*B*_*c*_ − *B*)^*−*1^.

**FIG. S6:**
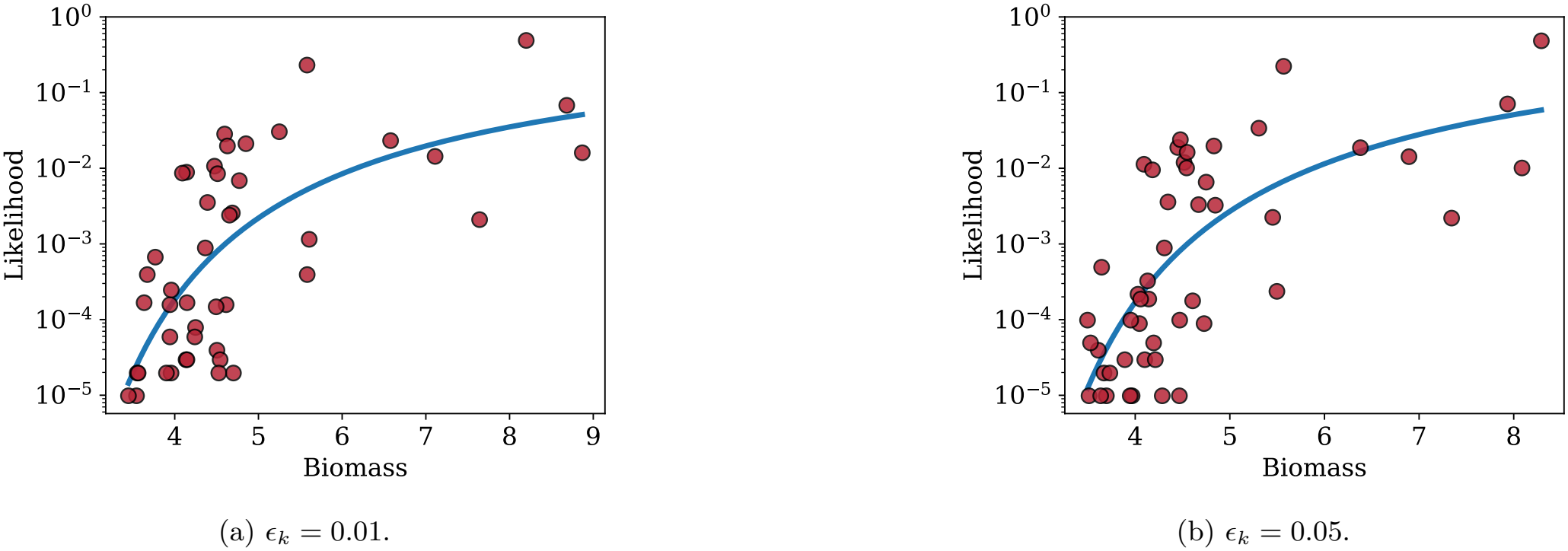
Biomass–likelihood relation when carrying capacity *K*_*i*_ varies around 1 with small variation *ϵ*_*k*_. In both cases, the positive correlation of likelihood biomass relation remains. Curves indicate fits to the log-hyperbolic relation log (*p*) ∝ (*B*_*c*_ − *B*)^*−*1^.

Next, instead of fixed *S* = 100, we take different system size We observe that for *S* = {50, 150}, the biomass– likelihood relationship still shows a positive correlation. As we decrease the system size the number of stable states decreases.(Fig. S7).

Next, in the main text, we sample each species’ initial abundance from a uniform distribution between 0 and 1. Here we show that the biomass–likelihood relationship is robust to another reasonable sampling scheme: that of sampling the initial conditions from a Gaussian centred at 0 with standard deviation 0.3, truncated to the positive orthant to ensure nonnegative abundances. We still find that the biomass–likelihood is qualitatively similar (S8). Even though sampling initial conditions differently does change the likelihoods of some states and we gain few extra states of low likelihood, the overall relationship is robust.

We also test robustness to heterogeneous growth rates *r*_*i*_ across species. In the main text, we set *r*_*i*_ = *K*_*i*_ = 1; here we relax this by drawing *r*_*i*_ from a uniform distribution on [1 − *ϵ*_*r*_, 1 + *ϵ*_*r*_]. As not ed in [43], when the *r*_*i*_ values are different, the value of the Lyapunov function in a local extremum becomes 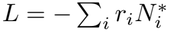 (weighted biomass), so it is actually this quantity rather than biomass that we expect to be more closely correlated with basin of attraction size. This quantity, which represents an alternative weighting over abundances, can also be interpreted as the net growth rate of the stable subset community in the absence of inhibition. We observe that for *ϵ*_*r*_ = {0.01, 0.05 }, the weighted biomass–likelihood relationship shows a positive correlation (Fig. S9). With increasing *ϵ*_*r*_, the correlation only slightly decreases and eventually saturates at roughly 0.5, still significantly positive (Fig. S9(c)). Further, this correlation persists both when considering the biomass of each state as well as the weighted biomass, with the latter showing a slightly stronger correlation (Fig. S9(c)). These results show that the biomass–likelihood relationship is robust to the inclusion of growth-rate variation between species.

**FIG. S7:**
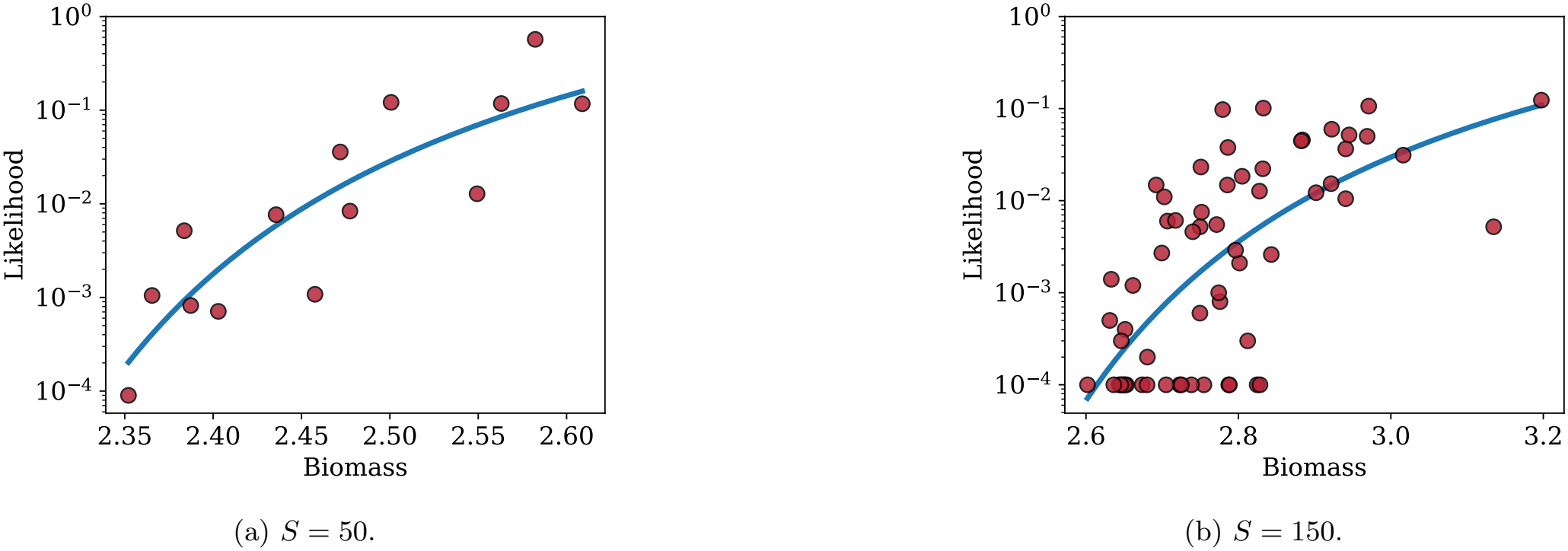
Biomass–likelihood relation when number of species (*S*) varies. In both cases, we continue to observe a positive correlation between likelihood and biomass. Curves show fits to the log-hyperbolic relation log (*p*) ∝ (*B*_*c*_ − *B*)^*−*1^.

**FIG. S8:**
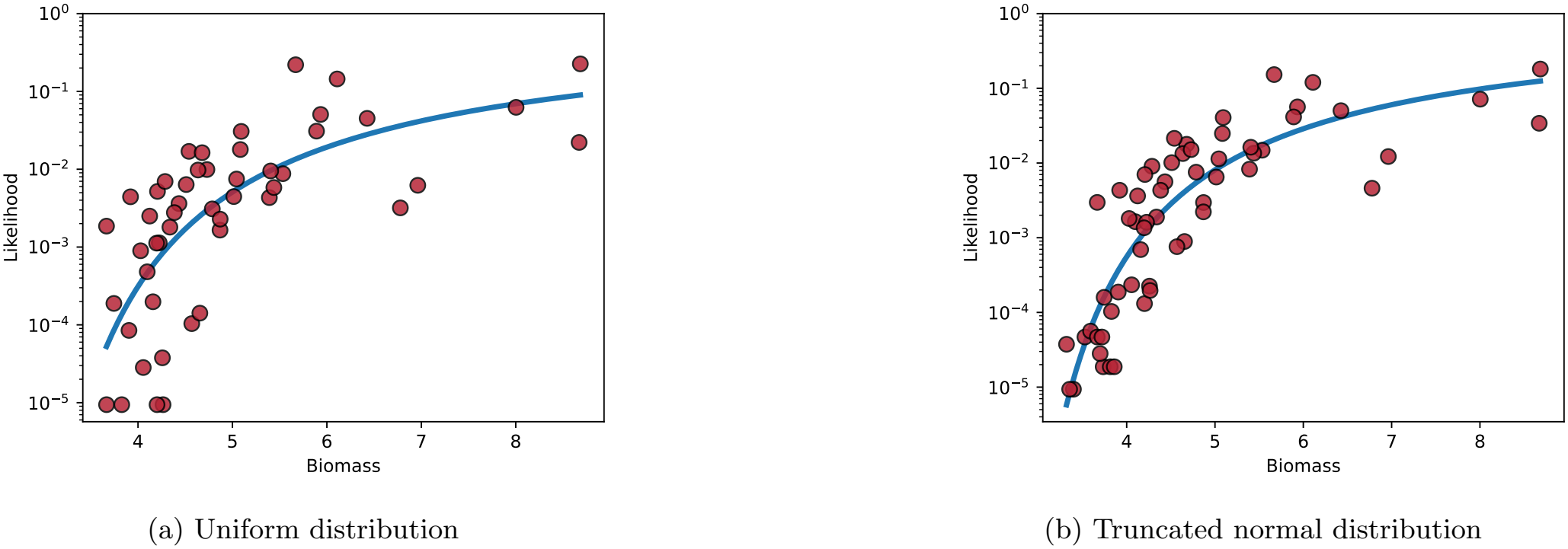
Our results are robust to different initial condition sampling schemes: shown are both (a) initial conditions sampled from a uniform distribution in [0, 1] and (b) a standard normal distribution truncated below 0. Both initial condition sampling schemes show a similar likelihood-–biomass relationship. Curves show fits to the log-hyperbolic relation log (*p*) ∝ (*B*_*c*_ − *B*)^*−*1^.

To confirm that the biomass–likelihood relationship is not specific to the particular (*µ, σ*) used in the main text, we sweep over a grid of values spanning the multistable phase and compute *γ*, the Pearson correlation between biomass and log-likelihood, for each. We find that *γ* remains positive throughout the multistable regime (Fig. S10), confirming that the positive relationship between biomass and likelihood is a generic feature of this phase.

To understand how reliable our likelihood estimates are, we measured the uncertainty in them when repeating our simulations for the same interaction matrix. We repeated the simulation procedure described in Appendix A 5 times for the same interaction matrix *A* used in the main text Figs. 1c and 3, each time sampling a different set of 10^6^ random initial conditions. We quantified uncertainty using the standard error in the mean (SEM) in the predicted likelihoods for each state. We observed a negligible uncertainty (*<* 1%) for the majority of states. Only for states with the lowest likelihoods (≤ 10^*−*5^) did we observe noticeable uncertainty (Fig. S11).

**FIG. S9:**
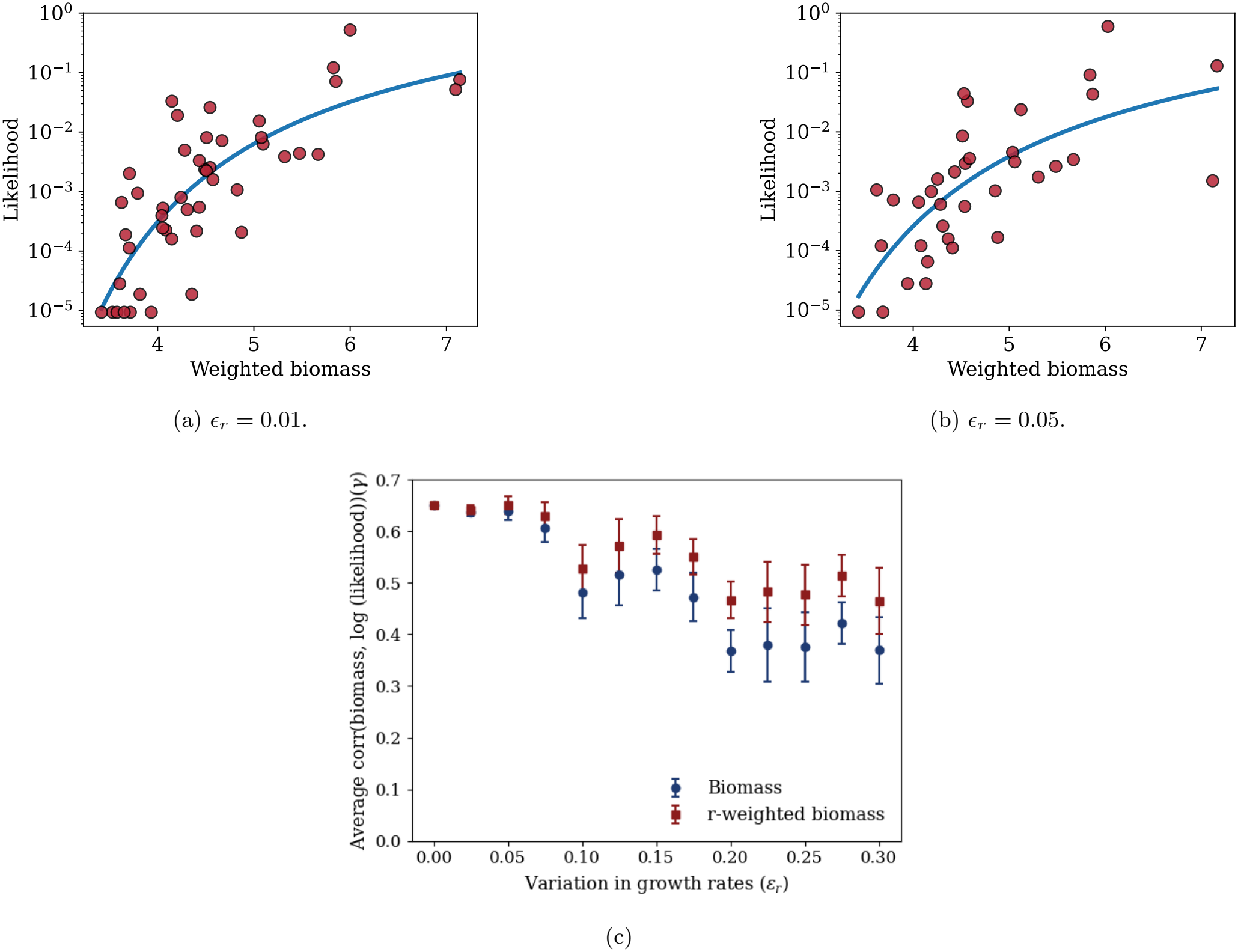
Biomass–likelihood relation when growth rate *r*_*i*_ varies around 1 with small variation *ϵ*_*r*_.(a,b) In both the case likelihood and growth-rate weighted biomass 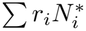 shows positive correlation. Curves indicate fits to the log-hyperbolic relation log (*p*) ∝ (*B*_*c*_ − *B*)^*−*1^. (c) Increasing variation in growth rates *r*_*i*_ maintains a positive Pearson correlation *γ* between biomass and log-likelihood (blue) for random interaction matrices, while slightly decreasing its magnitude. The correlation eventually saturates to roughly 0.4. Using growth rate weighted biomass 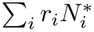 results in a slightly stronger correlation (roughly 0.5, red). Results are shown for 5 independent realizations of sampling *r*_*i*_ for each *ϵ*_*r*_, while keeping the interaction matrix *A* random but fixed for all realizations. Error bars indicate SEM across realizations.

Finally, we added demographic noise to the GLV dynamics to study the effect of fluctuations on the likelihood– biomass relationship. We incorporated noise into the dynamics through

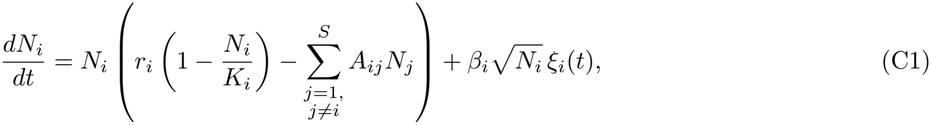

where *β*_*i*_ ≥ 0 is the noise amplitude of species *i*, 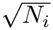_*i*_ reflects the Poisson scaling of birth–death fluctuations (vanishing at *N*_*i*_ = 0, preserving the extinction boundary), and *ξ*_*i*_(*t*) is Gaussian white noise with ⟨*ξ*_*i*_(*t*) ⟩ = 0 and ⟨*ξ*_*i*_(*t*) *ξ*_*j*_(*t*′)⟩= *δ*_*ij*_ *δ*(*t* − *t*′). We still observed a positive likelihood–biomass relationship after introducing noise, showing that high-biomass states generally remain more likely. The likelihood–biomass relationship remains qualitatively similar after adding noise, showing that low-biomass states do not become significantly less likely due to fluctuations (Fig. S12).

We also directly compared the likelihoods of the same states with and without fluctuations, and found that lowbiomass states can also be surprisingly resilient to fluctuations. In some cases, fluctuations actually increased the likelihood of the low-biomass states (Fig. S13), making them even more resilient than high-biomass states. These observations may reflect the interplay between the widths and depths of basins of attraction, but understanding this in detail requires further study, which we leave for future work.

**FIG. S10:**
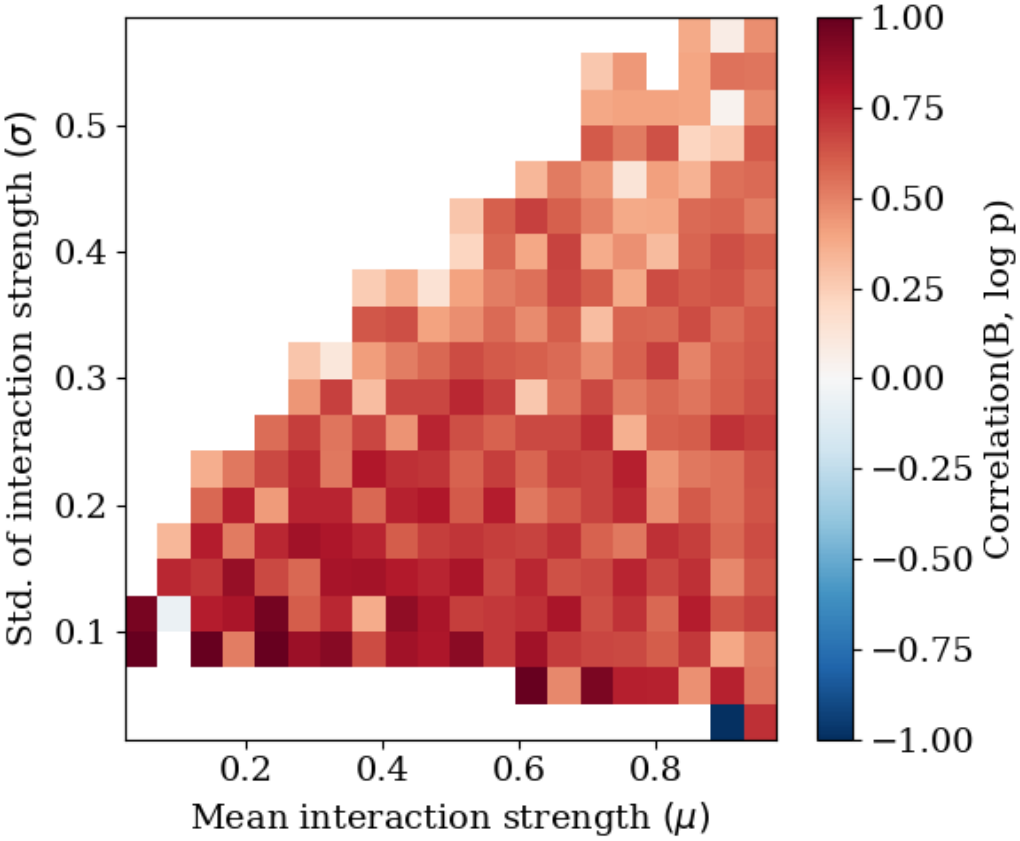
The Pearson correlation *γ* between biomass and log-likelihood across the multistable phase space is positive. Each point corresponds to a different (*µ, σ*) value; *γ* remains positive throughout, confirming that the biomass–likelihood relationship is a robust feature of the multistable regime independent of the specific parameter choice.

**FIG. S11:**
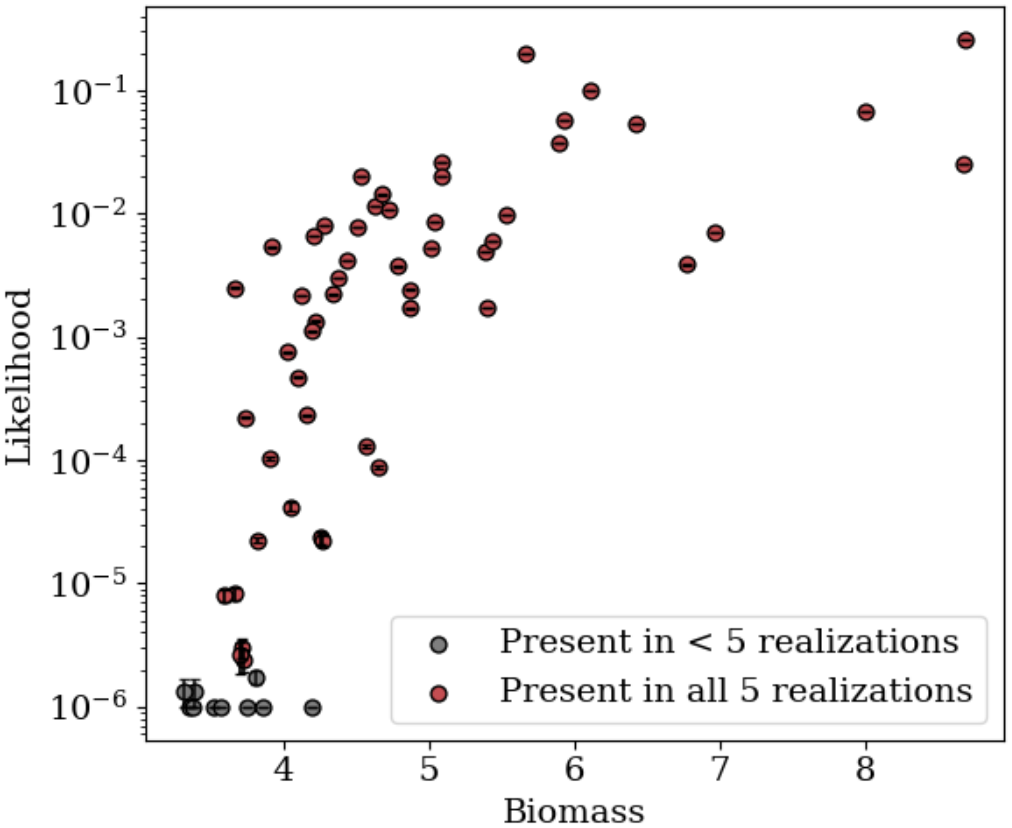
Uncertainty estimates for state likelihoods. Each point shows the likelihood of a state across multiple realizations of initial conditions for the same matrix. High-likelihood states appear in all five realizations (red) and show very small variation. In contrast, low-likelihood states show larger fluctuations, and some of them (gray) do not appear in all realizations. Results are shown for five different realizations. Error bars show the SEM across the realizations in which the state is observed.

**FIG. S12:**
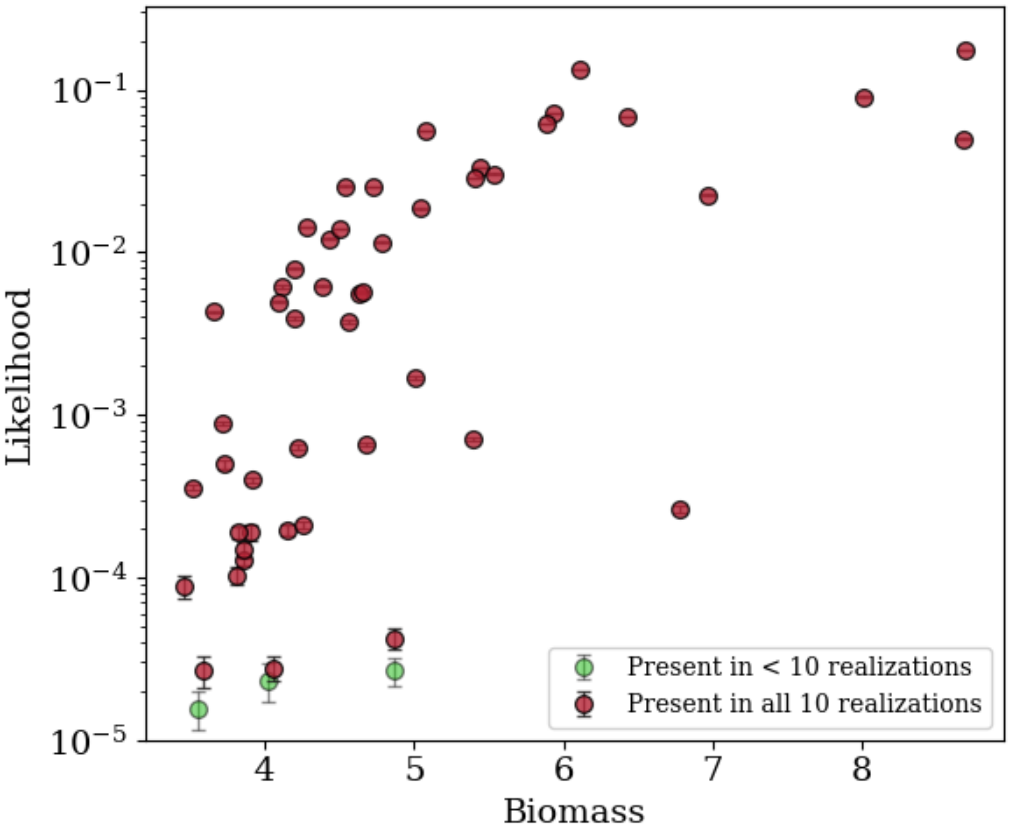
In the presence of demographic noise, the likelihood and biomass of different stable states are still positively correlated. We find that both low-biomass and high-biomass states are resilient to fluctuations. In all 10 realizations, we observe roughly the same set of low-biomass and high-biomass states. Each point shows the average likelihood of a state across 10 realizations of initial conditions for the same matrix and the same noise strength. Error bars show the SEM across the realizations in which the state is observed.

**FIG. S13:**
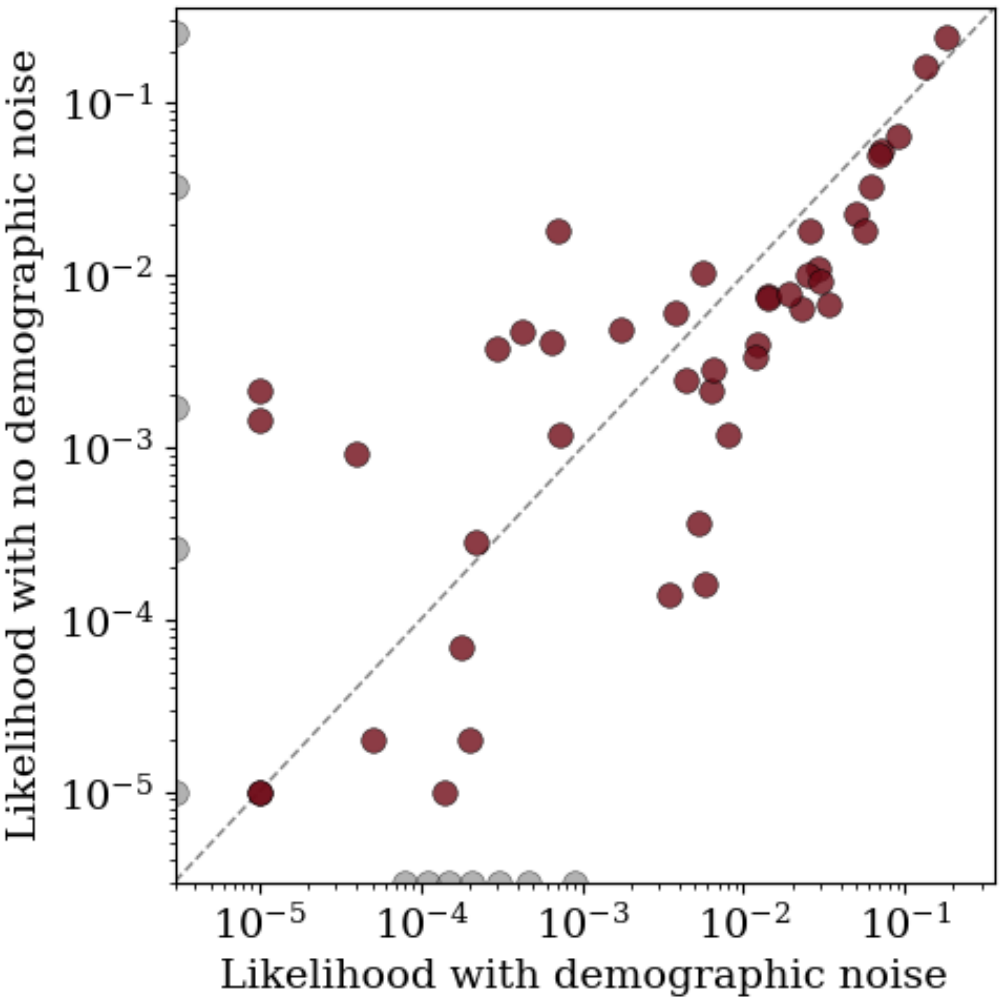
Demographic noise changes the likelihood of ecological states. States observed under both conditions (red) show that high-likelihood states remain highly likely even in the presence of noise, while low-likelihood states can shift, with some becoming more likely and others less likely. States that disappear under noise (gray, left edge) or appear only with noise (gray, bottom edge) suggest that demographic stochasticity can make rare states dominant and dominant states rare.

## Appendix D: Exact solution of likelihoods and biomasses in the monodominant model

In the monodominant model, the likelihood and biomass can be calculated exactly, which is not possible for random matrices. Since the monodominant model is the simplest one in which we can compute the likelihood–biomass relationship exactly, we start there. Afterwards, we will extend our calculation to block-structured interaction matrices, where more than one species can survive. Although random matrices do not obviously show clear block patterns, species that coexist in a state derived from a random disordered interaction matrix still contain block-like features, as we show explicitly in Appendix G. In the subsequent Appendices, we generalize the monodominant analysis to block models, and use the block model as a framework for developing a technique to predict likelihoods for such random matrices. As we show in the main text, this allows us to predict likelihoods for random matrices reasonably well using only the biomass and diversity of each state, along with the mean interaction strength *µ* which is a coarse parameter.

In the monodominant model all species *i* interact with the same interspecific strength *D*, while they differ in self-inhibition *K*_*i*_. For simplicity, we take *r*_*i*_ = 1 for all species (though we relax this in a later subsection in this Appendix). Thus, the self-inhibition of species *i* is given by *A*_*ii*_ = 1*/K*_*i*_, and species with higher carrying capacity have weaker self-inhibition. To ensure that at most one species survives at steady state we consider only cases where *D > A*_*ii*_ for all *i*.

The GLV equation in the monodominant case can be written as

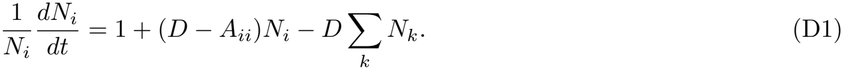

The last term −*D* ∑_*k*_ *N*_*k*_ is identical for all species. Thus the species-specific contribution is (*D* − *A*_*ii*_)*N*_*i*_. Define

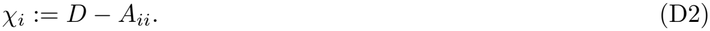

From (D1), we can see that the hyperplane defined by *χ*_*i*_*N*_*i*_ = *χ*_*j*_*N*_*j*_ is a fixed locus of the dynamics. Assuming this relation holds at any time *t*, we have *χ*_*i*_*dN*_*i*_*/dt* = *χ*_*j*_*dN*_*j*_*/dt*, since the right-hand side of (D1) is the same for both species. This shows that this hyperplane is a separatrix between the two species, and any initial condition with *χ*_*i*_*N*_*i*_ *> χ*_*j*_*N*_*j*_ must end in a steady-state solution where *N*_*j*_ = 0. Because this geometric relation holds for every pair of species, for any initial condition the species that dominates must be the one with the largest value of *χ*_*i*_*N*_*i*_(0), which we call the dressed initial condition.

Geometrically, we can interpret this as a division of the positive orthant by *S*(*S*− 1)*/*2 hyperplanes into *S*! cones, each associated with an ordering of the *χ*_*i*_*N*_*i*_ values. Within each cone, all initial conditions lead to the final steady state *i* with the largest value of *χ*_*i*_*N*_*i*_. To compute the probability of a given final state for a given distribution of initial conditions, we need to determine the fraction of initial conditions associated with the set of cones that lead to the given final state. (Note that while this general cone structure was described in [43], the exactness of this geometric decomposition for the monodominant systems was not realized there.)

In the subsequent analysis we consider this question from two different perspectives: first, given the initial conditions on each species taken from a uniform distribution from 0 to 1, giving a hypercube of initial conditions within which we calculate the fraction in each domain of attraction; second, scaling the initial conditions by *χ*_*i*_, so the sides of the initial condition boxes vary, but such that we simply determine the largest value of *χ*_*i*_*N*_*i*_(0) directly.

### Two species

Consider two species with self-inhibition parameters satisfying *A*_11_ *> A*_22_ (so *χ*_1_ *< χ*_2_). In our simulations we draw initial abundances for both species from the same uniform distribution between 0 and 1, so the set of initial conditions is a square in the (*N*_1_, *N*_2_) plane (see Fig. S14a). The separatrix can be found analytically; it passes through the point ((*D* −*A*_22_), (*D* −*A*_11_)) in the coordinate system used in the simulation and partitions the square into two basins.

Viewed in the dressed-initial-condition picture, set *X*_1_ ∼ Uniform(0, *χ*_1_) and *X*_2_ ∼Uniform(0, *χ*_2_) where *X*_*i*_ = *χ*_*i*_*N*_*i*_(0). The separatrix is then given by the line *χ*_1_*N*_1_(0) = *χ*_2_*N*_2_(0), which in the dressed (*X*_1_, *X*_2_) coordinates is simply *X*_1_ = *X*_2_ (see Fig. S14b). The question “what is the likelihood of species 2 winning?” reduces to computing

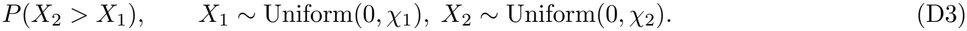

We compute *P*(*X*_2_ *> X*_1_) by splitting into cases. If *X*_2_ lies between *χ*_1_ and *χ*_2_ then *X*_2_ is always larger than *X*_1_. If *X*_2_ lies between 0 and *χ*_1_ then *X*_2_ is larger than *X*_1_ with probability 1*/*2 (by symmetry of two identical Uniform(0, *χ*_1_) draws). Therefore

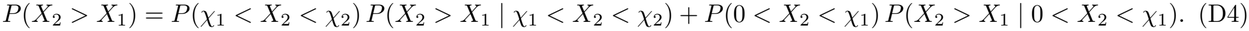

We also have that

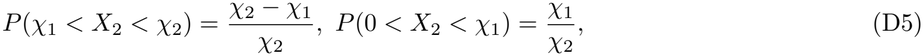

and the conditional probabilities are 1 and 1*/*2 respectively. Hence

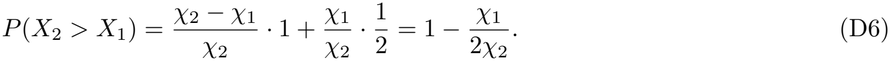

**FIG. S14:**
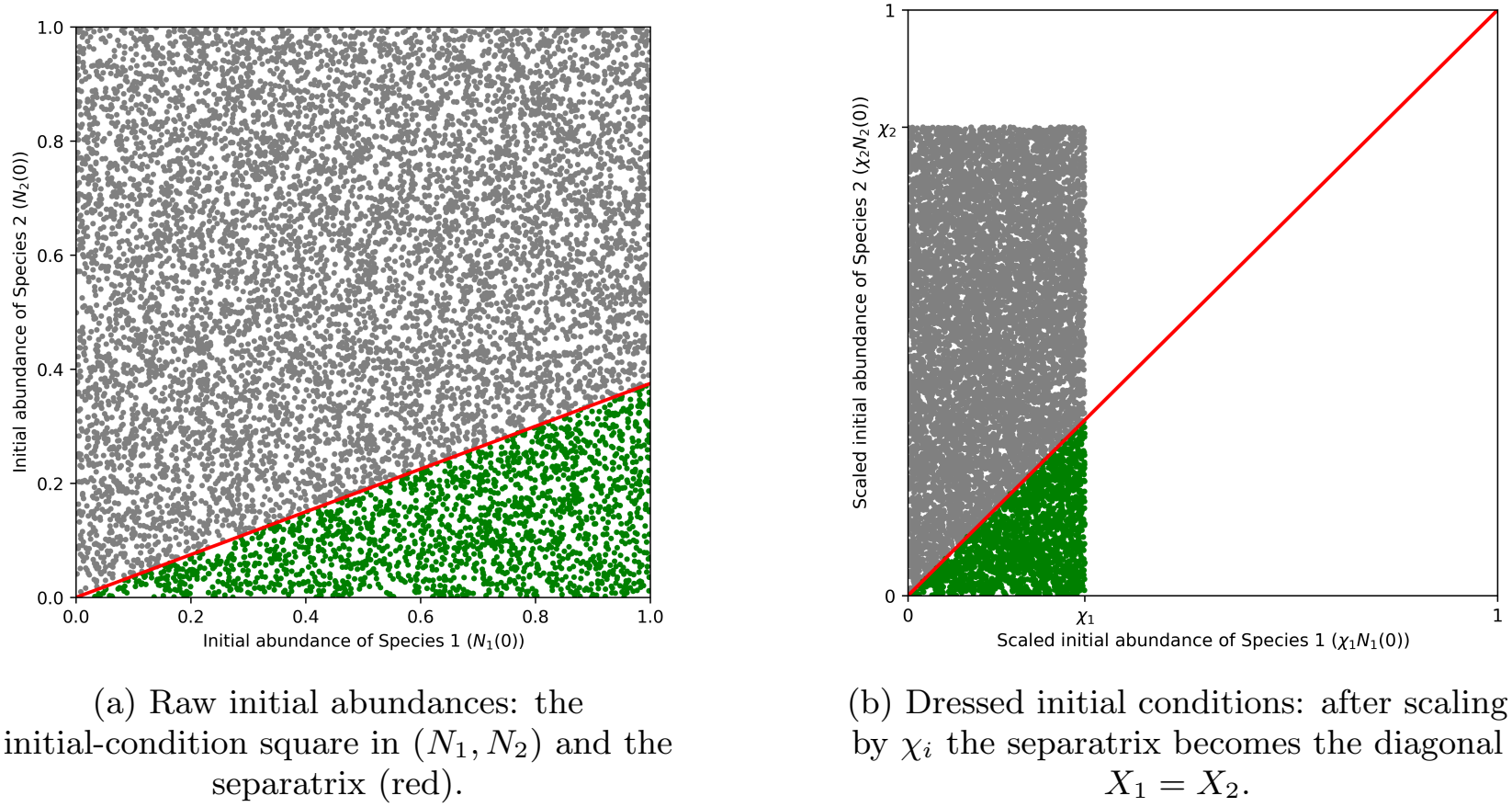
Green regions indicate outcomes where species 1 wins; gray regions indicate outcomes where species 2 wins. In panel (a) the separatrix has slope *χ*_1_*/χ*_2_ in the (*N*_1_, *N*_2_) plane. In panel (b) the dressed variables (*X*_1_, *X*_2_) make the separatrix the diagonal *X*_1_ = *X*_2_.

**FIG. S15:**
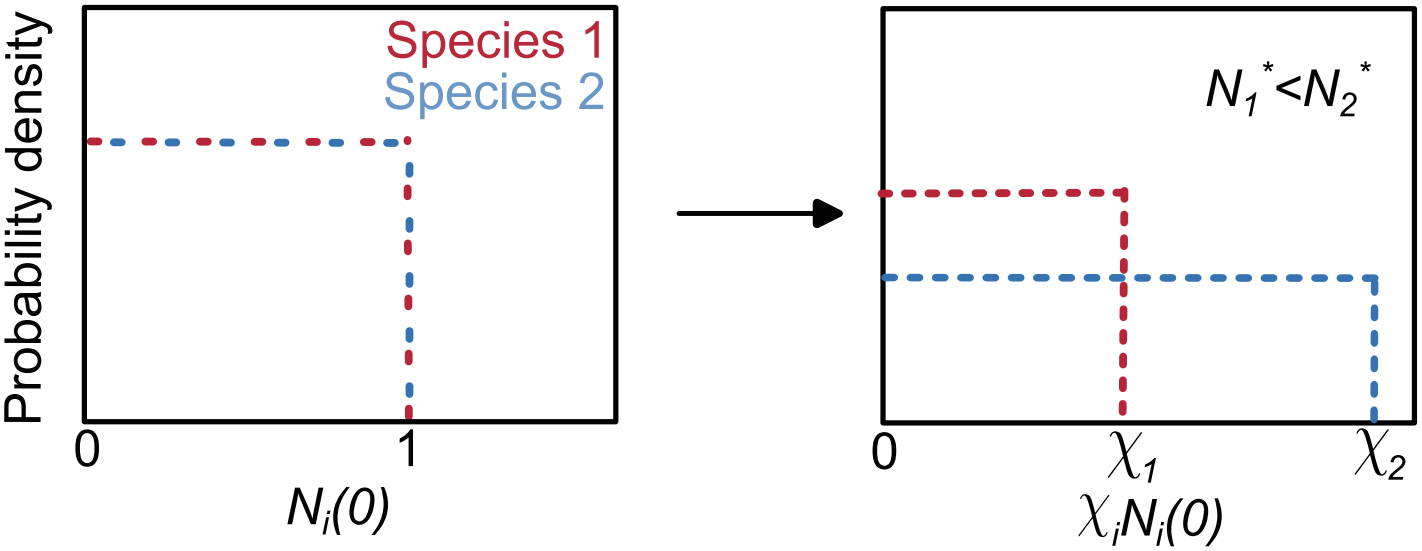
Left: PDFs of the raw initial abundances for species 1 and 2 (both drawn from Uniform(0, 1)) are identical. Right: PDFs of the dressed initial conditions *X*_*i*_ = *χ*_*i*_*N*_*i*_(0) are uniform on [0, *χ*_*i*_]. Because *χ*_2_ *> χ*_1_, species 2 has extra support on the interval (*χ*_1_, *χ*_2_] that species 1 cannot access, giving species 2 an advantage in the maximum.

By complementarity,

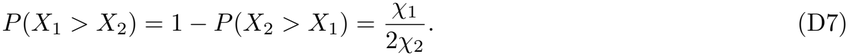

### Many species

Consider *S* species with self-inhibition values ordered as *A*_11_ *> A*_22_ *> … > A*_*SS*_. Define *χ*_*i*_ = *D* − *A*_*ii*_ so that

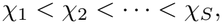

i.e. species with smaller self-inhibition have larger *χ*_*i*_.

**FIG. S16:**
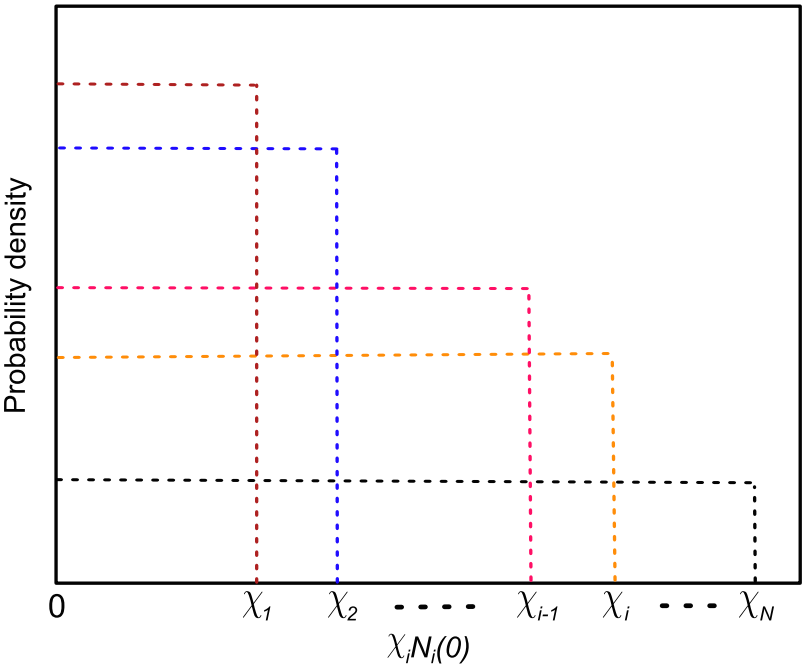
PDF of dressed initial conditions for *S* species. The self-inhibition values are ordered as *A*_11_ *> A*_22_ *>…> A*_*SS*_, so the corresponding upper bounds satisfy *χ*_1_ *< χ*_2_ *< … < χ*_*S*_. A larger *χ*_*i*_ (smaller self-inhibition) gives species *i* a wider range of dressed initial values and thus a larger chance to be the maximum.

For each species *j* let

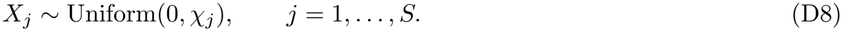

The density of *X*_*i*_ is

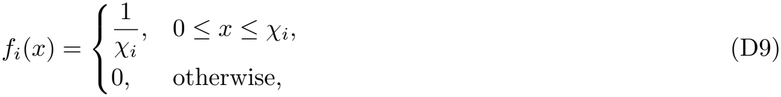

and the cumulative distribution of *X*_*j*_ is

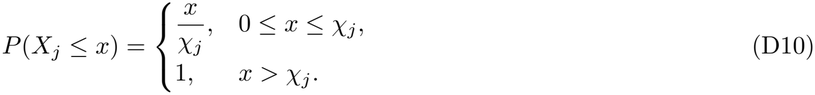

The probability that *X*_*i*_ is the largest among the *S* variables is

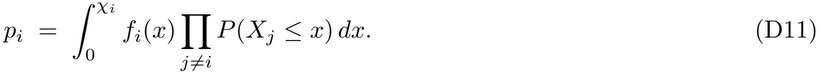

Because *P*(*X*_*j*_ ≤ *x*) changes form at the breakpoints *χ*_*j*_, partition the interval [0, *χ*_*i*_] using

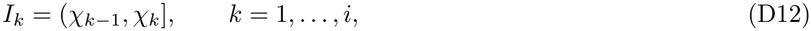

with the convention *χ*_0_ = 0. On the subinterval *I*_*k*_ exactly the first *k* − 1 variables have reached their full support (so *P*(*X*_*j*_ ≤*x*) = 1 for those), while the remaining variables satisfy *P*(*X*_*j*_ ≤*x*) = *x/χ*_*j*_. Hence on *I*_*k*_ the product becomes

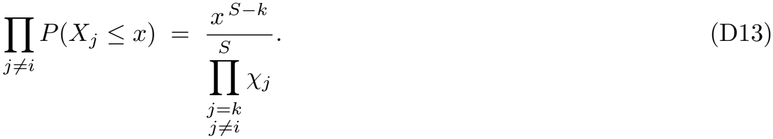

Using *f*_*i*_(*x*) = 1*/χ*_*i*_, the contribution of the interval *I*_*k*_ to *p*_*i*_ is

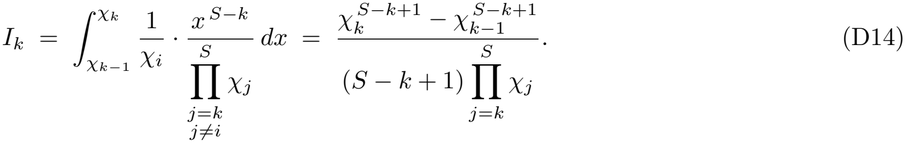

Note that this contribution can be understood in a simple geometric fashion: 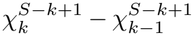 is the volume of the region in which all values of *χ* ≤ *χ*, for *i* ≥ *k*, removing the volume where all *χ*_*i*_ *< χ*_*k−*1_. The product over *χ*_*j*_’s is the appropriate normalization factor for this volume. Within this volume there are *S*− *k* + 1 species that are equally likely to have the greatest value of *χ*_*i*_*N*_*i*_. So this volume gives an equal contribution to each of those species for the probability to be greatest.

Therefore the exact probability that species *i* is the largest is the sum over these contributions:

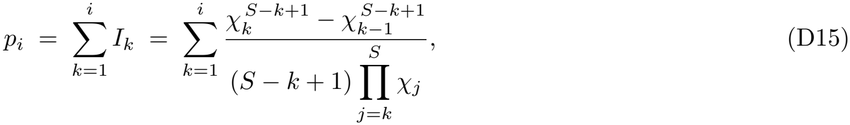

The difference in probability between species *i* and *i* − 1 is

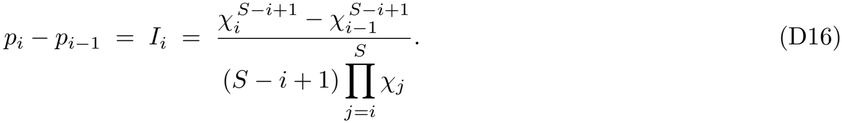

The above expression shows that when species *i* has lower self-inhibition than species *i* − 1, i.e. *A*_(*i−*1)(*i−*1)_ *> A*_*ii*_ (or equivalently *χ*_*i*_ *> χ*_*i−*1_), its likelihood always increases.

### Extension of the monodominant model to heterogeneous growth rates

In the main text and throughout the analysis above, we set *r*_*i*_ = 1 for all species (except see Appendix C, Fig. S9). Here we extend the monodominant model to allow species-specific growth rates *r*_*i*_, which is important because in the GLV model (Eq. (1) in the main text) the growth rate and carrying capacity can play distinct roles. With heterogeneous *r*_*i*_, the monodominant GLV equation becomes

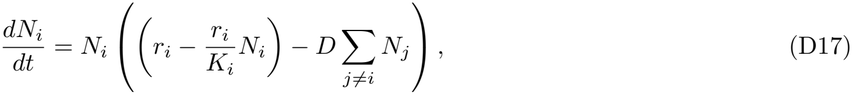

To test if we can quantitatively predict the winner in this modified monodominant model, we attempt to use a similar approach as before: that of defining an appropriate dressed initial condition. Rewriting Eq.(D17) in terms of the per capita growth rate of species *i* we get

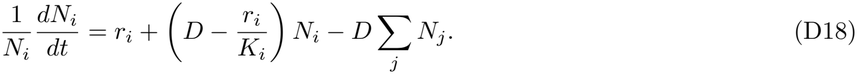

The last term is again common to all species, but the species-dependent part now involves *r*_*i*_ explicitly. We can now define a modified version of the dressed initial condition to account for growth rate differences as:

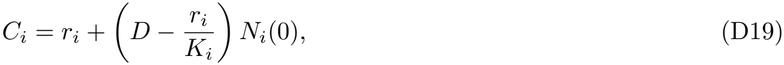

where *K*_*i*_ is the steady-state abundance or biomass of species *i*. When *r*_*i*_ = 1, we get *C*_*i*_ = 1 + *χ*_*i*_*N*_*i*_(0), and like in the previous section, the species with the largest *χ*_*i*_*N*_*i*_(0) wins. However, when *r*_*i*_ varies between species, the modified dressed initial condition *C*_*i*_ acquires a complicated dependence on both *r*_*i*_ and *K*_*i*_. The species with the largest *C*_*i*_ has the largest per capita growth rate at *t* = 0.

To test whether the modified dressed initial condition *C*_*i*_ can quantitatively predict the winner in this modified monodominant model, we perform simulations in which we systematically control the extent of growth rate variation. We introduce a parameter *ε* and for each species *i* we draw its growth rate *r*_*i*_ from a uniform distribution centered at 1 with width 2*ε*, such that *r*_*i*_ ∈ [1 − *ε*, 1 + *ε*]. When *ε* = 0, all species have the same growth rate *r*_*i*_ = 1 and we recover the original monodominant model, Eq.(D1). Increasing *ε* increases the extent of growth rate variation.

In the presence of growth rate variation between species, we find that the modified dressed initial condition *C*_*i*_ only approximately predicts which species wins, as shown in Fig. S17. Specifically, while *C*_*i*_ perfectly predicts the winner in the absence of growth rate variation, in the presence of substantial growth rate variation, *C*_*i*_ correctly predicts the winner only ∼ 50% of the time. This is still significantly better than the null expectation of 1*/S* = 5% for *S* = 20 species. Furthermore, the winner predicted using the largest *C*_*i*_ is still typically among the top 2 or 4 species (70% and 85% of the time, respectively), significantly better than the null expectation (dashed lines in Fig. S17). This shows that the modified dressed initial condition *C*_*i*_ remains a somewhat good approximation of the winner. This reduction in predictive power might arise to the fact that in the presence of growth rate variation, the separatrices between the different basins of attraction are no longer straight lines, but become curved. This may result in the transient nonlinear dynamics mattering a lot more in predicting the winner, which cannot be captured by the initial conditions alone. We leave a full exploration of this for future work.

**FIG. S17:**
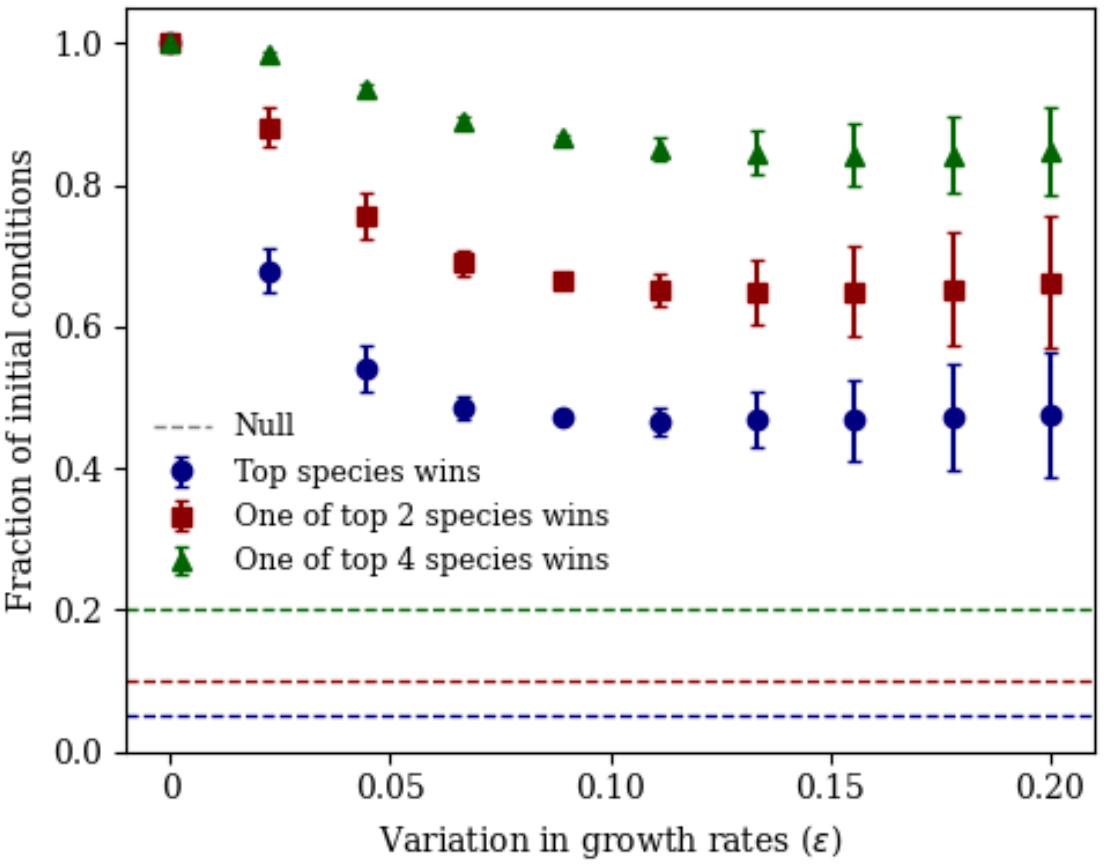
The species with the largest *C*_*i*_(0) remains a good approximation of the winner in the monodominant model with heterogeneous growth rates. The plot shows the fraction of initial conditions in which the winning species is the one with the largest *C*_*i*_(0) (blue), one of the two largest (red), or one of the four largest (green), as a function of the variation in growth rates *ϵ* among a total of *S* = 20 species. Here, *r*_*i*_ is drawn from a uniform distribution on [1 − *ϵ*, 1 + *ϵ*]. Dashed lines of the same colors show the corresponding null expectation, *k/S* for the top *k* species, obtained if the winner were chosen uniformly at random among all *S* species. Error bars indicate SEM for 30 monodominant interaction matrix realizations.

Beyond predicting the winning species, we also examine how the correlation between biomass and likelihood depends on the growth rate variation. We find that in the monodominant model, with increasing growth rate variation, the average correlation between biomass and likelihood *γ* = corr(biomass, log(likelihood)) decreases significantly from *γ* ∼ 0.85 and saturates to *γ* ∼ 0.15 around *ε* = 0.20 (Fig. S18). No tably, while its magnitude decreas es, the correlation still remains positive. Using the growth rate-weighted biomass 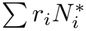 instead of the true biomass 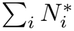 increases this correlation to *γ*∼ 0.35 in the presence of substantial growth rate variation. These results show that the correlation between biomass and likelihood in the monodominant model always remains positive, but its magnitude is sensitive to growth rate variation.

Note that this decrease in the correlation is qualitatively similar to what we observed in the case of random disordered matrices (Fig. S9(c)). However, the effect is much more pronounced in the monodominant model than the random matrix case, where the decrease in the correlation is relatively small. This might be because the monodominant model is much more sensitive to the growth rates of individual species, since each stable state contains only one species, while stable states in the random disordered matrices contain multiple species. We leave a full investigation of this effect for future work.

**FIG. S18:**
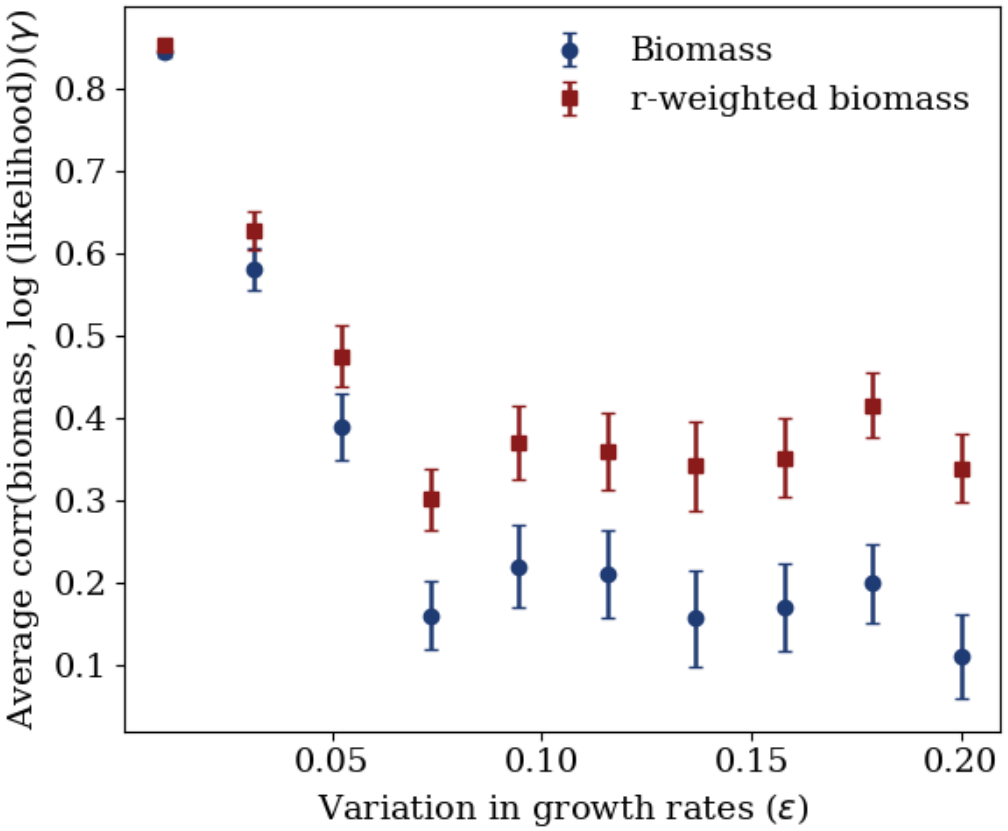
Increasing variation in growth rates *r*_*i*_ maintains a positive Pearson correlation *γ* between biomass and log-likelihood in the monodominant model, but decreases its magnitude. Here, *r*_*i*_ is drawn from a uniform distribution on [1 − *ϵ*, 1 + *ϵ*] in the monodominant model. The correlatio n saturates eventually, but to a positive value and never becomes negative. Using growth rate weighted biomass 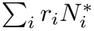 produces systematically higher correlations, especially as the variation in growth rates increases. Error bars indicate SEM for 30 monodominant interaction matrix realizations.

## Appendix E: Solution of likelihoods and biomasses for block models

In the monodominant model, the chance of a species winning depends on its self-inhibition *A*_*ii*_ (or equivalently its eventual biomass *B*_*i*_ = 1*/A*_*ii*_). If all species’ initial conditions are the same uniform distribution, then this difference in self-inhibitions causes changes in the slope of the separatrices dividing their basins of attraction. But, a random disordered matrix case does not map easily to the monodominant model since in each state, the former has several coexisting species. This motivates us move on to an enhanced variant of the monodominant model that allows more than one species to coexist in a state.

The simplest such model is to have the interaction matrix consist of blocks, where species within a block *α* inhibit both each other and themselves with a common strength *A*_*αα*_, and species in different blocks inhibit each other with a different, stronger strength *D*. We assume no overlap between blocks, which means that any species belongs exclusively to one block state. In this case only one block can win given any set of initial conditions, and all species within the winning block survive. We call this a “neutral block” model since each species in a block is identical to all others and thus in ecology jargon, all species within a block are “neutral”. We can show that the neutral block model maps exactly to a monodominant model. To see this, note that we can write down the GLV equations governing the dynamics of the total abundance (biomass) *B*_*α*_ =∑_*i∈α*_ *N*_*i*_ of each block *α* as

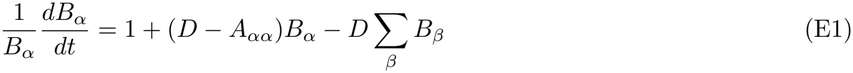

At steady state the final abundance of the winning block *α* will be 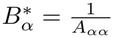. Eq.(E1) maps exactly to Eq.(D1). The steady state abundance of each block *α*, now the inverse of *A*_*αα*_, replaces the steady-state abundance of each species, which was the inverse of *A*_*ii*_. Following the monodominant results, the winning block will be the one with the largest value of the dresssed initial condition, now *χ*_*α*_*B*_*α*_, where *χ*_*α*_ = *D* − *A*_*αα*_ and *B*_*α*_ = ∑_*i ∈ α*_ *N*_*i*_ being the sum of initial abundances of species within block *α*. If each block *α* contains *L*_*α*_ species, then given uniform initial conditions for each species’ abundances, the block’s initial abundance follows the Irwin–Hall distribution. At sufficiently large *L*_*α*_ (here, 5), this distribution agrees extremely well with a Gaussian with mean ^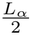^ and standard deviation ^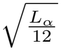^ (Fig. S19). Thus, for blocks of sufficient size (as shown, 5 is enough), the key difference between this neutral block model and the monodominant model is that the distribution of dressed initial conditions for each block will be a Gaussian rather than a uniform distribution. This will affect the likelihood of each block quantitatively, but not change the central insight that the dressed initial condition, which as we have shown depends crucially on the self-inhibition, perfectly predicts which block wins. Note that this choice of initial distribution will strongly favor larger blocks; with truly neutral interactions within each block, depending upon the context another distribution such as uniform across *B*_*α*_(0) might be natural, but in the context of the random matrix systems for which we are developing this theory, where each species is treated as a priori the same regarding initial conditions, the Irwin-Hall distribution is what we want to analyze. Note also that if the distribution on each species is taken to be something else that is the same across species, such as a Gaussian truncated to the positive orthant, for blocks of sufficient size we will again have a Gaussian distribution on the dressed initial conditions.

**FIG. S19:**
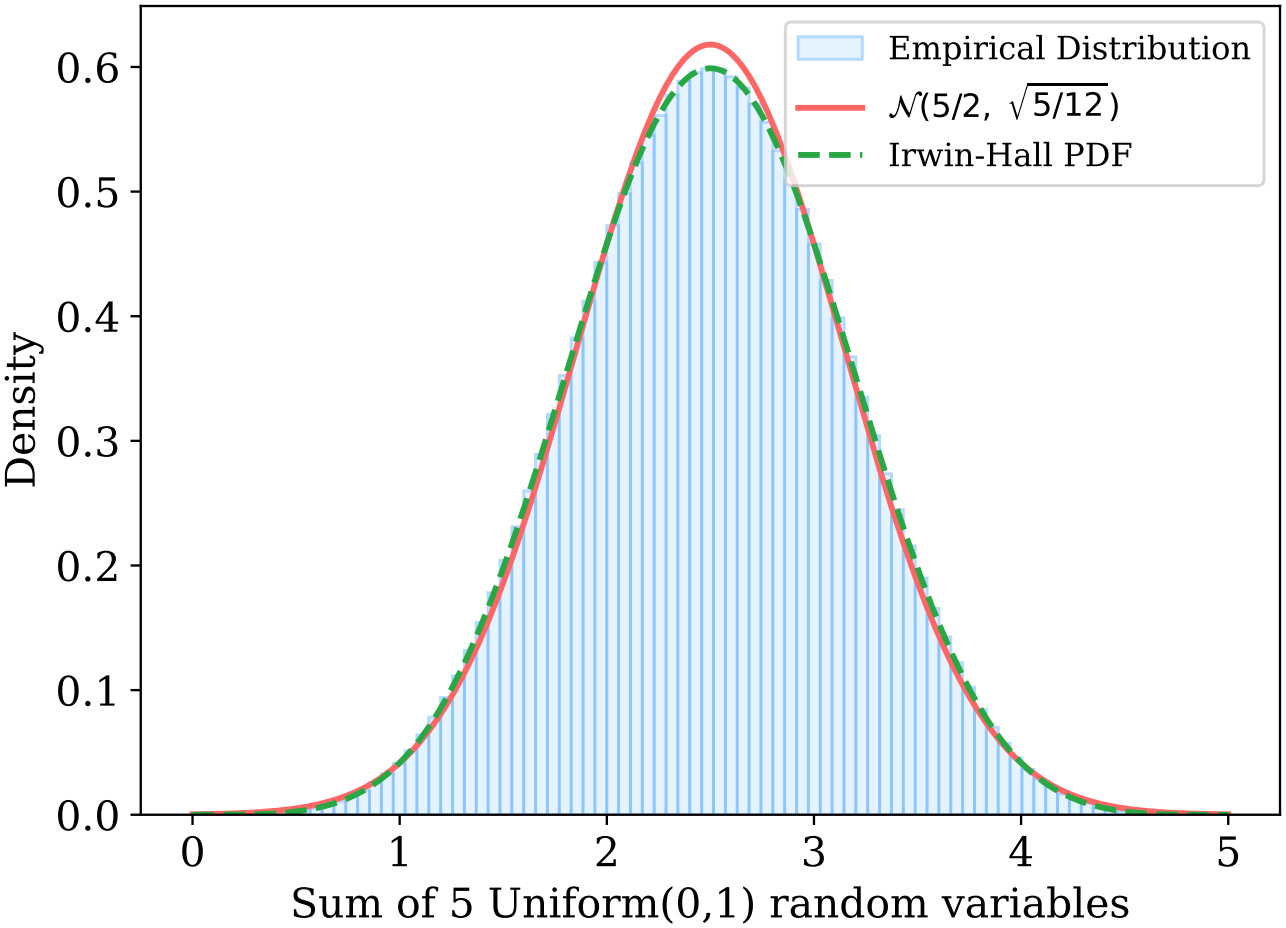
The sum of 5 independent uniform random variables (blue) compared with a Gaussian distribution with mean 5*/*2 and standard deviation 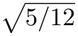 (red). The Irwin–Hall distribution matches the Gaussian approximation very well. The empirical distribution contains 10^4^ samples.

The neutral block model assumes that all species’ self-inhibition is the same as their inhibition of each other species in the same block, but in our random disordered matrices all species’ self-inhibitions are 1. To account for this, we now consider a different block interaction matrix, which only differs from the neutral block in that the self-inhibitions of all species is 1. All other interspecies inhibitions of species in the same block *α* are *A*_*αα*_ and across blocks are *D*. This is the block model we use in the main text. We will show that likelihoods in this block model are determined in a very similar fashion to that of the completely neutral block model, with an effective self-inhibition that again corresponds to the inverse of the equilibrium biomass for each block. To see this, let us begin with the species-level dynamics. The abundances of each species *N*_*i*_ evolve according to:

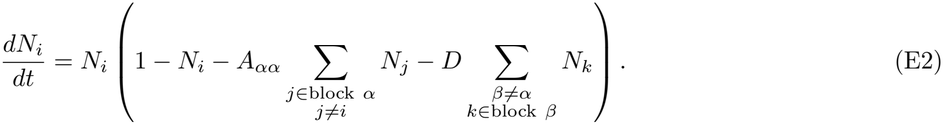

At steady state, all species in block *α* have the same abundance, i.e. 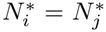 for all *i, j* ∈ block *α*. The steady state condition becomes

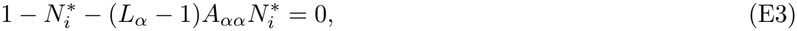

which gives

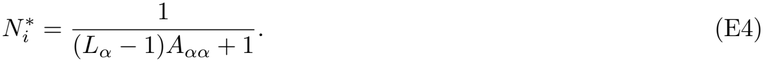

Hence the steady state biomass of block *α* is

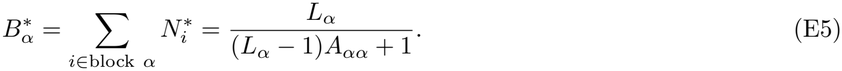

Summing equation (E2) over all *i* in block *α*, we obtain the effective dynamics for the blocks in terms of their current abundances *B*_*α*_ as

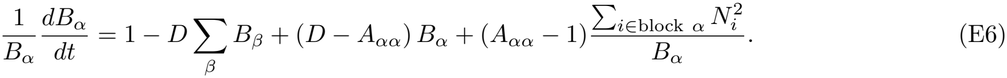

Note that the dynamics within each block in this model is no longer neutral, but tends to equilibrate species within each block when 1 *> A*_*αα*_, so that for a given total block biomass *B*_*α*_ the species abundances are driven within each block to the local quasi-equilibrium

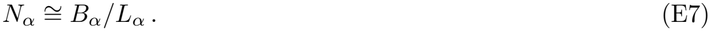

It is thus helpful to rewrite Eq.(E6) in the form

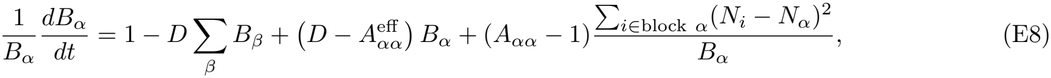

where the effective self-inhibition

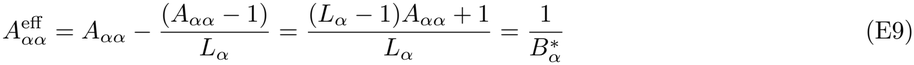

is exactly the inverse of the equilibrium biomass for each block. The last term in Eq.(E8) corresponds to statistical fluctuations from the initial conditions that rapidly dissipate over time as each block locally equilibrates, and add some noise but should not significantly impact the relative probabilities of ending up in different equilibria with random initial conditions, over most ranges of the parameters. Thus, we see that the effective dynamics even with the unit self-inhibition is essentially identical to that of the block monodominant model with neutral conditions within each block, with the proper value of the effective self-inhibition matching the inverse biomass.

As an illustration, consider two blocks. Just as in the monodominant model, we might describe the dynamics in terms of a phase portrait where we follow the abundances of the two blocks *B*_1_ and *B*_2_. The basins of attraction for the two blocks will be separated by a separatrix in the (*B*_1_, *B*_2_) plane given by the line:

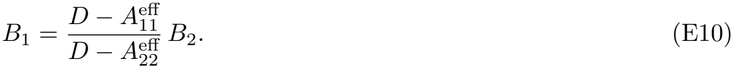

Based on our argument above, we take the effective self-inhibition of each block as the inverse of its equilibrium biomass. From Fig. S20b and Fig. S21a we see that this works well. If both blocks have the same size *L* but different equilibrium biomass, the initial distribution is Gaussian with a roughly circular spread with its center on the line *B*_1_ = *B*_2_, while the separatrix is tilted towards the *x*-axis compared to this line (Fig. S20a). On the other hand, if the blocks have equal equilibrium biomass but different sizes, the separatrix coincides with *B*_1_ = *B*_2_, but the initial distribution bivariate Gaussian has a spread which is both shifted and stretched due to differences in block size (Fig. S20b). This shows that, in block models, both the equilibrium biomass and size of a block affect its likelihood: equilibrium biomass tilts the separatrix, while block size moves and streches a block’s initial abundance distribution.

Similar to our treatment of the monodominant model, we can simplify the analysis by scaling the initial condition distribution in an appropriate way while also avoiding the need to tilt the separatrix. The key insight is that—when both the biomass and block size vary (Fig. S21a)—their combined effect can be captured by scaling a block’s total initial abundance by the factor

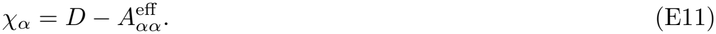

As shown earlier in this Appendix, for sufficiently large blocks, a block’s total initial abundance can be well-approximated by a Gaussian with a mean ^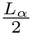^ and standard deviation 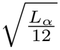 . The block with the largest dressed initial abundance is the winner (Fig. S21). Thus, the distribution of dressed initial condition *X*_*α*_ for block *α* also follows a Gaussian with

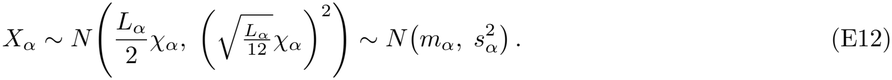

**FIG. S20:**
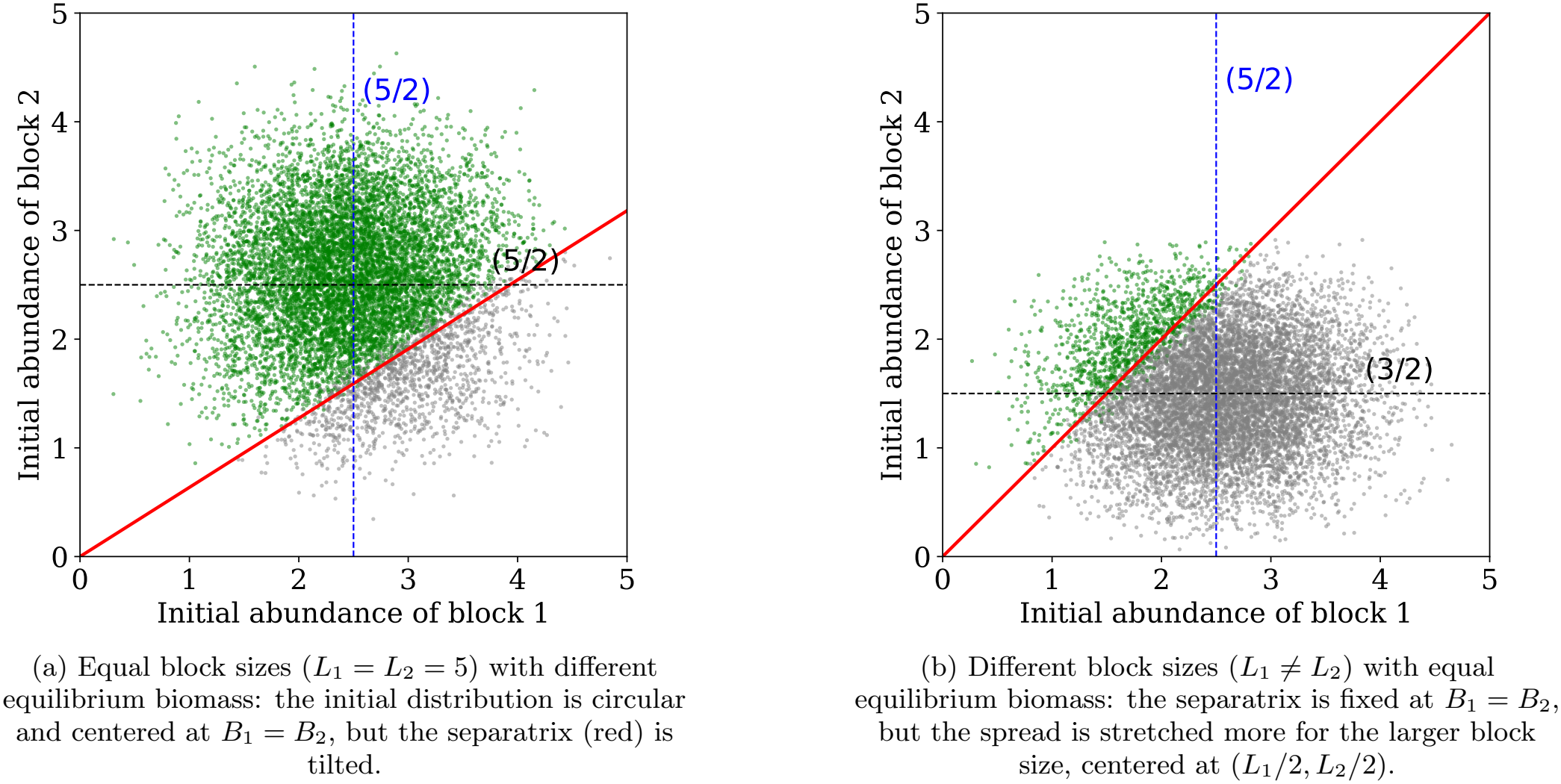
Effect of equilibrium biomass and block size on the likelihood. Green regions show outcomes where block 1 wins; gray regions where block 2 wins.

**FIG. S21:**
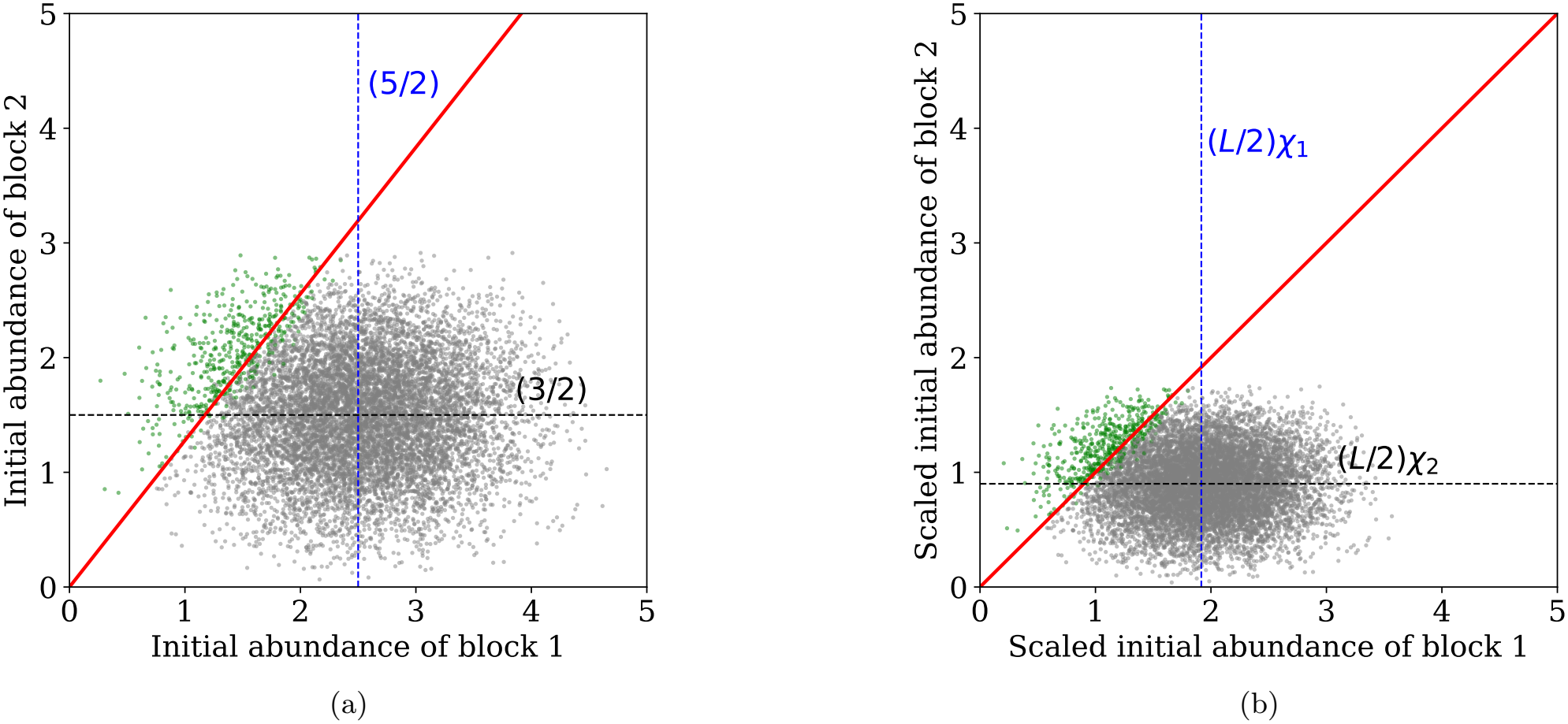
Two different ways of graphically depicting likelihoods of different blocks surviving in a case with different block sizes and different equilibrium biomasses. Scaling block abundances captures the effects of both equilibrium biomass and block size in determining relative likelihoods. Block with largest dressed initial biomass wins. Green regions: block 1 wins; gray: block 2 wins. (a) Without scaling, the separatrix has slope *χ*_1_*/χ*_2_, with initial abundance centered at (*L*_1_*/*2, *L*_2_*/*2), with *L*_1_ = 5 and *L*_2_ = 3. (b) After scaling, the separatrix becomes *X*_1_ = *X*_2_,with dressed initial abundance centered at (*χ*_1_*L*_1_*/*2, *χ*_2_*L*_2_*/*2).

**FIG. S22:**
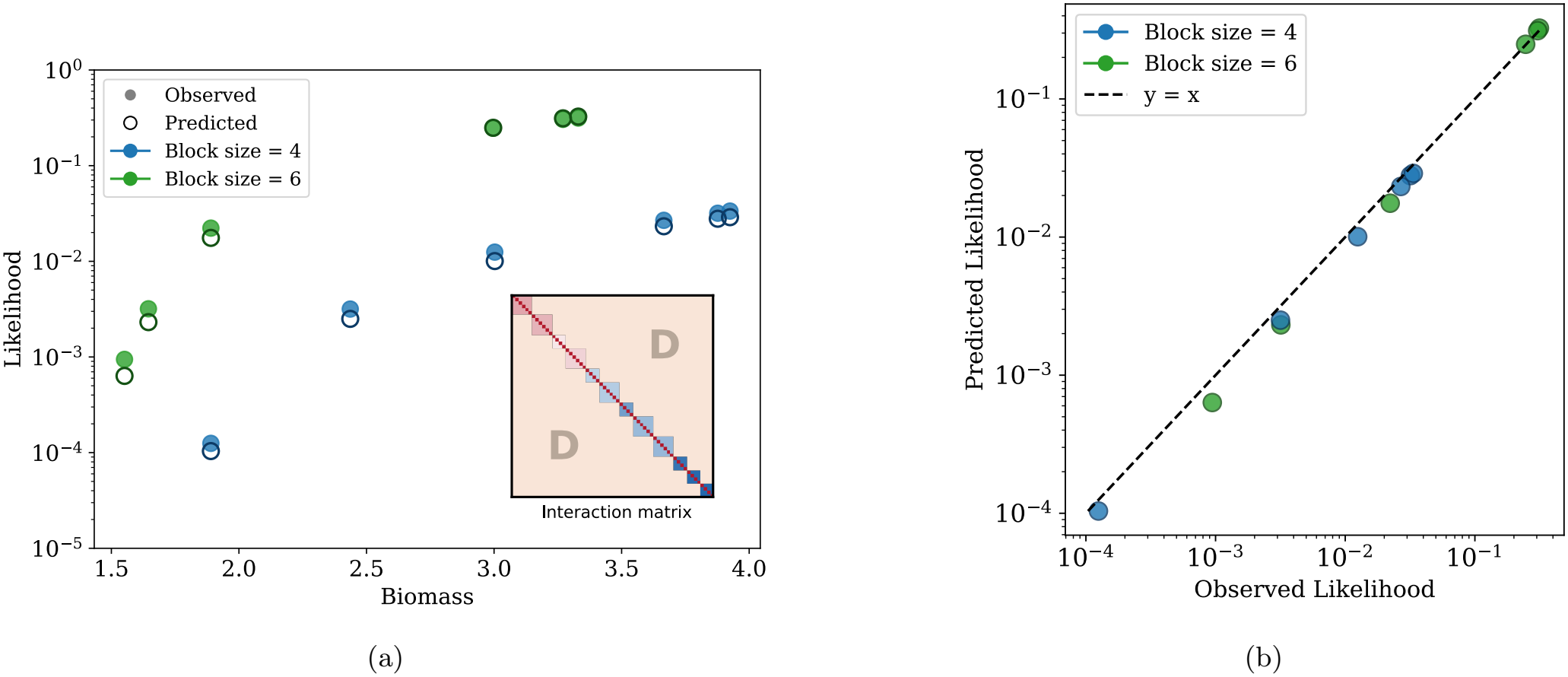
(a) Likelihood versus biomass for different block sizes, using an interaction matrix with 12 blocks (inset). Blocks were either of size 4 or 6, with randomly chosen self-inhibitions and inter-block inhibition *D* = 1.1. Some blocks had high likelihood despite lower biomass due to their larger size. Eq.(E18) predicted likelihoods extremely well. (b) Comparison of observed likelihoods with predicted likelihoods for the same matrix as in (a).

Now consider *N*_*B*_ independent blocks with dressed initial conditions 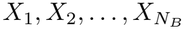, where

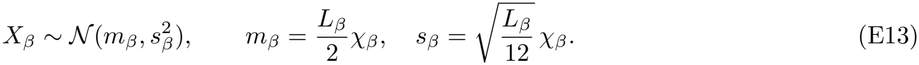

Following the same approach as in the monodominant many-species case, the probability that block *α* has the largest dressed initial condition *X*_*α*_ is

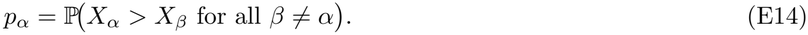

This can be expressed as the following integral

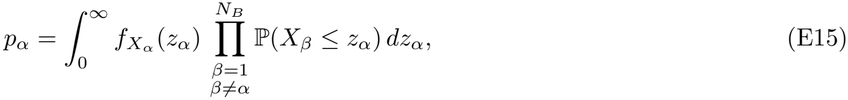

where 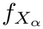 is the pdf of *X*_*α*_. Explicitly, this reads as

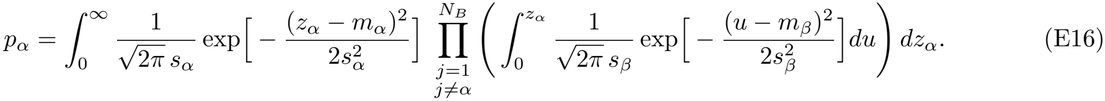

Equivalently,

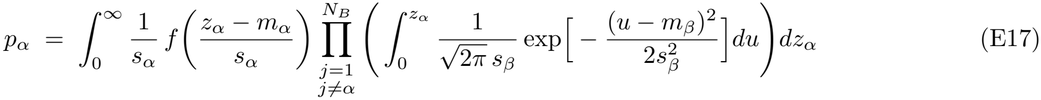

where 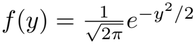 is the standard normal density. Note that the lower limit of the inner integrals in Eq.(E17) can be replaced by −∞ (for blocks of size *L* ∼5 or larger) without significantly affecting the quantitative contribution from these integrals. This is because virtually all of their mass is concentrated in the positive space of abundances. Thus, for convenience we may replace the inner integrals with the CDF of the Gaussian distribution, using which we get

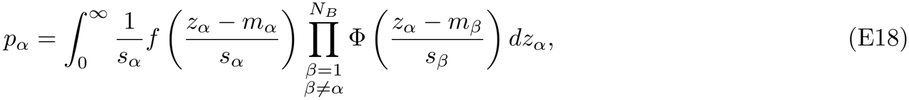

which is Eq. (5) in the main text. Here

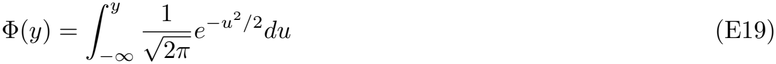

is the Gaussian CDF. In Fig. S22b–c, we show that the prediction from Eq.(E18) agrees well with the observed likelihoods for an example block matrix shown in Fig. S22a. This matrix has 12 blocks, each of size either 4 or 6 (chosen randomly). We set the inter-block inhibition *D* = 1.1 and all diagonal entries (representing species’ self-inhibitions) to 1. To assign within-block self-inhibitions *A*_*αα*_ for each block *α*, we randomly picked a biomass values uniformly randomly between 1.5 and 4. We then used Eq.(E5) to compute the corresponding within-block inhibition *A*_*αα*_ to get that equilibrium biomass. We then computed the likelihoods for each block numerically using simulations (Fig. S22a, closed circles) and compared them with the predicted likelihoods from Eq.(E18) (Fig. S22a, open circles). When the inter-state inhibition in a block-structured matrix was not constant, as in the random matrix case, we used *D* = ⟨*A*_*ij*_⟩ in Eq.(E18) to predict the likelihood of the blocks (Fig. S23). Overall, our formula captures the likelihoods well, with some minor deviations at low likelihoods, likely due to numerical errors. Replacing our Gaussian approximation with the exact Irwin–Hall distribution for block initial conditions yields very similar results. Taken together, we have obtained an analytic formula for the likelihood of a block state, which is analogous to the many-species monodominant case. This formula in Eq.(E18) accounts for blocks (states) with multiple species and Gaussian initial distributions. Each block’s probability to win depends on both its biomass and its block size, whereas in the monodominant model it depends only on biomass.

**FIG. S23:**
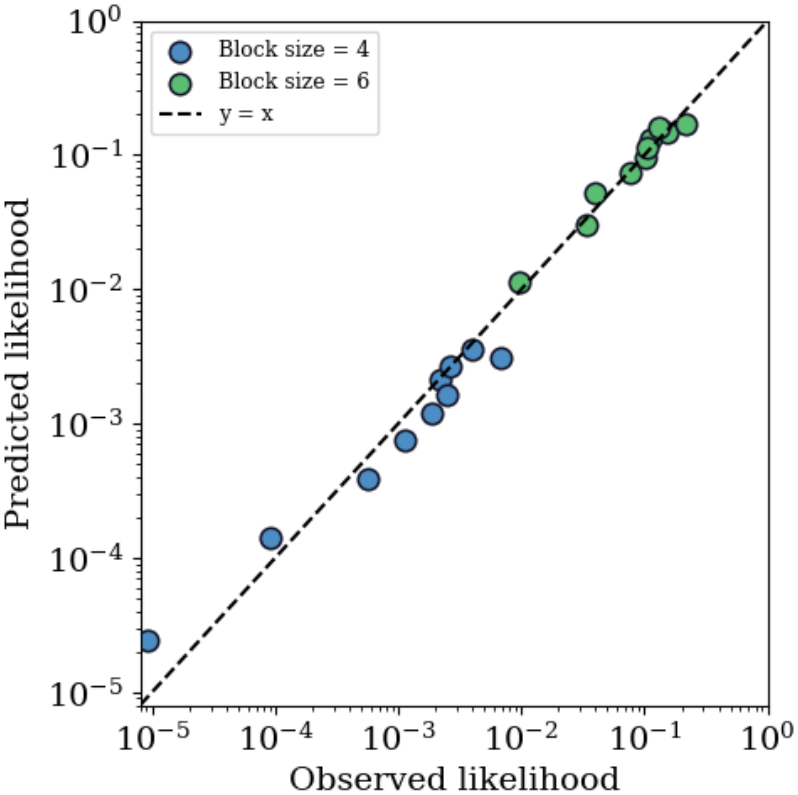
Variability in inter-state inhibitions did not significantly affect the predictions of our block model. We constructed a matrix with 10 blocks of size 4 and 10 blocks of size 6, with 10 biomass values linearly spaced from 3 to 7. We took the inter-state interaction from a gaussian distribution *N* (1.1, 0.1). We used *D* = ⟨*A*_*ij*_⟩ = 1.1 in Eq.(E18) to compute the predicted likelihoods. Our predictions agreed well with the observed likelihoods across both block sizes, with only minor deviations .

## Appendix F: Overlaps between states and a simple statistical model of states in ecosystems with random disordered interaction matrices

In this Appendix we describe the overlaps between states in these random disordered ecosystems, and describe a simple statistical model that captures some basic aspects of the number and size of the individual block states in these models

**FIG. S24:**
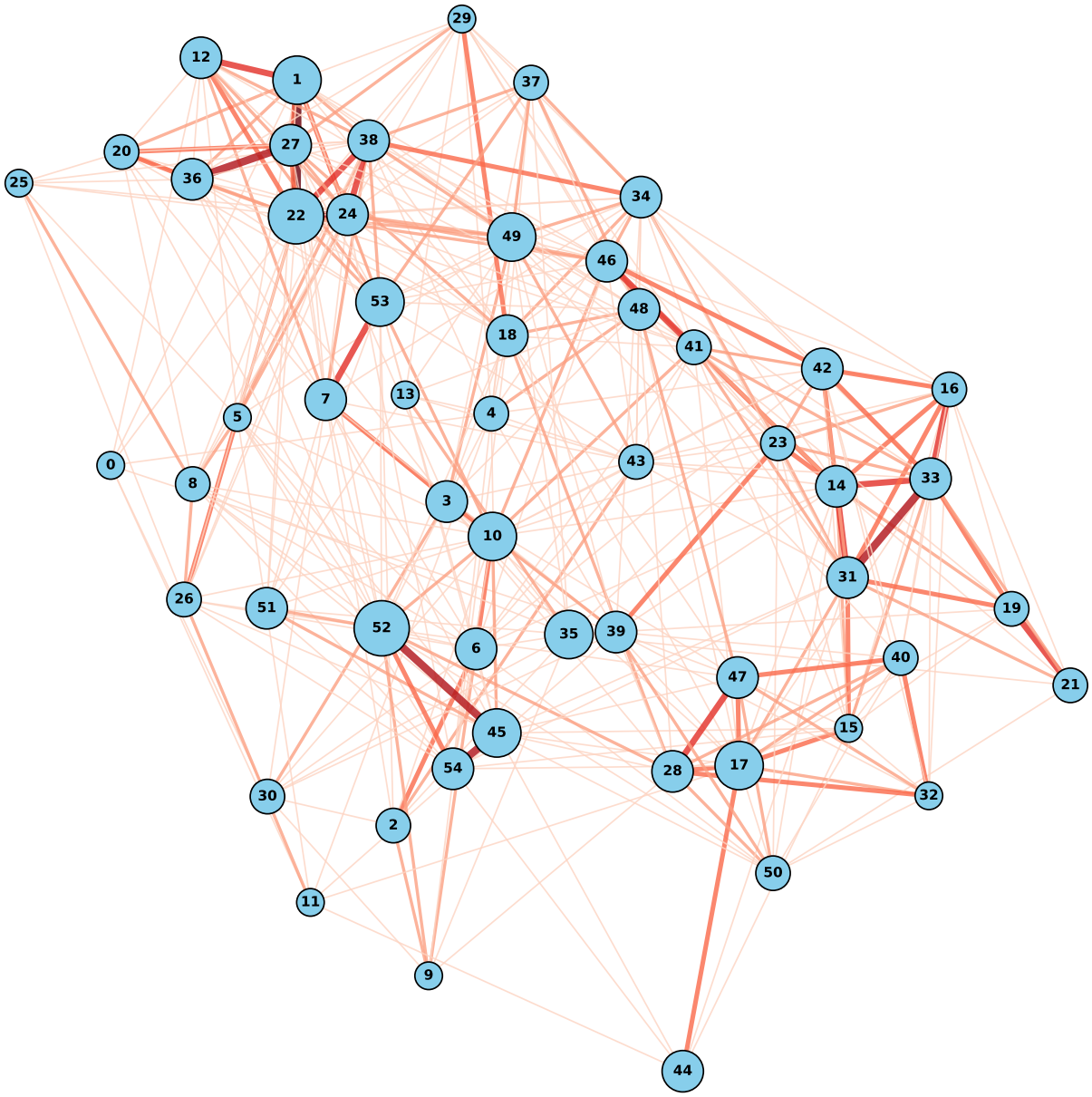
Each node represents a state and each edge shows that two states share common species. Thickness of the edge shows the number of common species between the states. The overlapping goes from 1 to 6 species. The node index follows the order of likelihood.The size of the nodes shows the diversity of the state,which goes from 4 to 8 species. We can see that states overlap with each other, so the random matrix does not form a perfect block structure.

When we predict the likelihood of different states from a random matrix, we apply the theory of block-structured matrices developed in Appendix E. In a perfect block-structured matrix, each species belongs to only one state. As we show in Fig. 2a, the theoretical motivation for treating states as blocks comes from the observation that species coexisting within the same stable state interact much more weakly with each other than expected from the overall interaction distribution. This reduced intra-state inhibition suggests that species within each state should form cohesive, block-like units.

In random matrices, we observe a richer structure where states naturally overlap through shared species. We can visualize this by constructing a network where each state represents a node, and we connect two nodes with an edge if they share at least one common species (Fig. S24). The thickness of each edge indicates the number of species shared between connected states. This network reveals the intricate interconnectedness among states, with many states sharing species across multiple connections. Rather than the discrete blocks described in Appendix E, we find a complex overlapping structure that reflects the natural complexity of random disordered ecosystems.

One direction in which the analysis of this paper could be extended is by taking further account of the detailed structure of individual blocks and the connectivity between blocks. This could likely lead to a more refined statistical model. As a first step in this direction, and as a way of understanding the appearance of blocks, we outline here a simple statistical model based on *k*-cliques in random graphs that seems to capture the structure of the size and number of block/states for the random ecosystems considered here

We begin by noting that even though in the random interaction matrix each interaction is chosen independently according to a given mean and variance, the combinations of species that persist in a stable block in general have smaller inter-species interactions, as illustrated in Fig. 2. Thus, to estimate for example the number of stable blocks of size *k* in a given system of size *S*, we can make a rough model in which a block is persistent when all inter-species interactions within the block lie below some specific value *a*. If we denote the probability that this occurs for a given interaction under the normal distribution on *A*_*ij*_ by *p*, then the probability that any specific subset of *k* species has all inter-species interactions less than or equal to *a* is 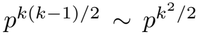, and the number of such subsets is *S*!*/*((*S k*)!*k*!) ∼*S*^*k*^, where the asymptotics are at large *k, S*, with *k* ≪*S*. This is essentially the problem of finding *k*-cliques in a random graph of size *S*.

In this simple model, the expected number of blocks of size *k* is given by

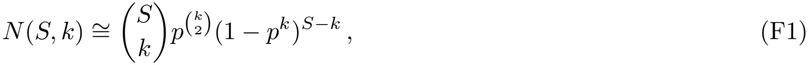

where the last factor comes from the condition that the block is uninvadable, hence is not a sub-block of a larger feasible stable block. Each of the (*S* −*k*) species outside the block can be added to it only if all *k* of its interactions with the block lie below *a*, which occurs with probability *p*^*k*^. The largest clique/block in this context has size *k*∼ 2 ln *S/* ln(1*/p*), which can be determined by finding the largest *k* such that the expected number of blocks of size *k* is *O* (1); this is a classic result in graph theory [59]. Similarly, a saddle point analysis shows that the typical maximal clique/block has size *k*^*\**^∼ ln *S/* ln(1*/p*), and the number of maximal cliques/blocks goes as 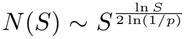. This argument explains why the stable states that we find in random matrices of size, e.g., *S* = 100 are relatively small (*k* ∼ 6).

This simple model for the number of maximal blocks matches very well with explicit computations; see Figure S25. Because in the parameter ranges we are considering here, the number and size of stable equilibrium states is relatively small, a simple pruned tree algorithm can enumerate all possible stable states for e.g. *S* = 100, *µ* = 0.5, *σ* = 0.3, where across an ensemble of 10 matrices the number of distinct equilibrium states is *N*_eq_ ≡83*±* 4. The pruning can be done efficiently using the fact that if a subset *S*^*′*^ has an unstable *A* matrix (negative eigenvalue), then there cannot be any stable subset *S*^*′′*^ where *S*^*′*^ ⊂*S*^*′′*^. Note that for this kind of system the standard ecological method of predicting the number of states based on a limited probabilistic sampling using, e.g., the Chao1 estimator [60, 61] based on the Good-Turing discovery rate [62–64] may not be trustworthy, particularly when the sampling only touches the large-probability tail of the distribution.

**FIG. S25:**
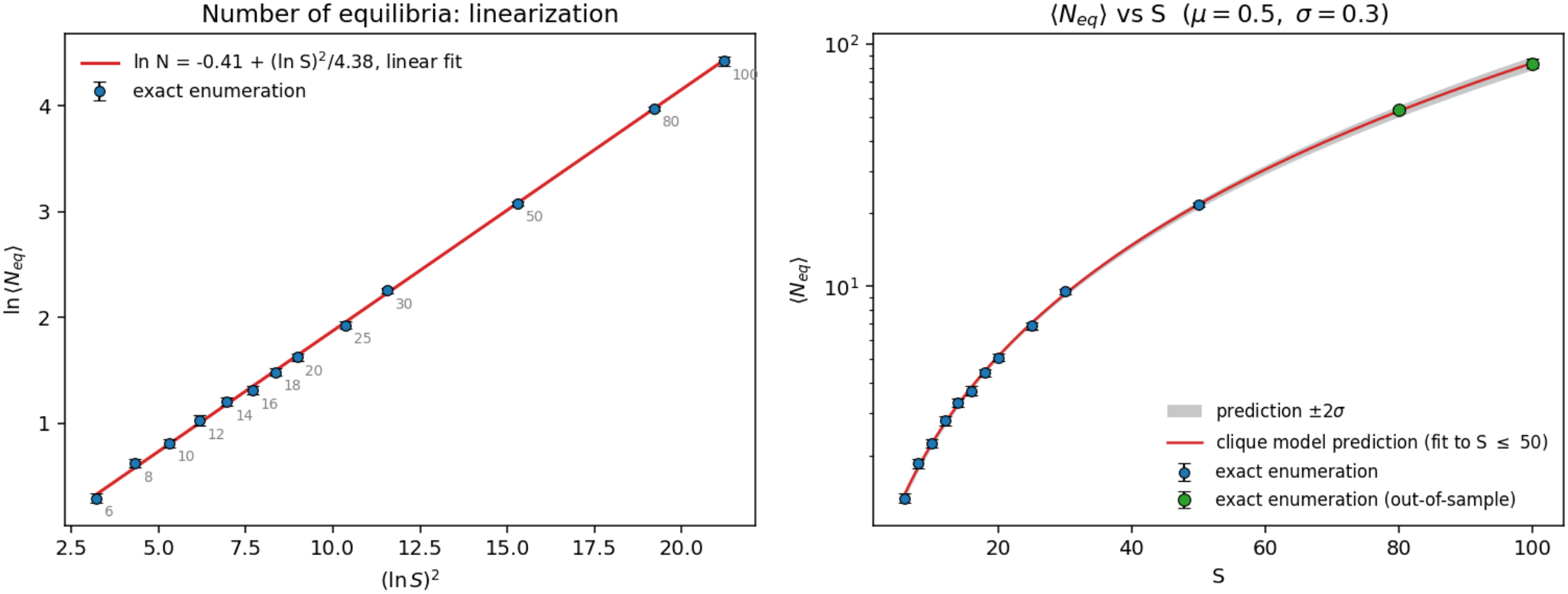
The clique model for equilibrium state distribution suggests a number of equilibria going as 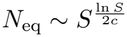, where *c* is a constant depending on the matrix ensemble. This matches explicit computation very closely. Left: a linear fit between ln *N*_eq_ and (ln *S*)^2^ matches data for number of equilibria with *S* from 4 to 100, explicitly calculated exactly for a sample of matrices using a pruned tree search; Right: a fit with numerical data from *S* ≤50 (80 matrices each at lower values, 100 matrices at *S* = 50 gives a precise prediction of values at *S* = 80, 100 (estimated from exact enumeration over 50, 10 matrices respectively).

Using similar logic, a similar estimate can be given for, e.g., the number of pairs of cliques of size *k, k*^*′*^ that overlap in *s* species. It would be interesting to develop this kind of model further to understand in more detail the expected structure of interactions between the stable states in this kind of random system and use that structure to make more precise predictions about domain of attraction sizes, but we leave further analysis in that direction for future work.

## Appendix G: Method to reveal emergent block structure in a random interaction matrix

The same species can appear in multiple stable states for any given random disordered interaction matrix, and as we show in Appendix F, the interaction matrix does not form a perfect block structure as assumed in our theory (Appendix E) due to overlapping states. In this appendix, we show that nevertheless, there is an emergent block-like organization when we reorder species in specific ways. Particularly, here we detail a possible systematic approach— which we use—that focuses on states that do not overlap.

Note that in this analysis, we use blocks that are identified as stable solutions from simulations with random initial conditions. An alternative approach that would be worth exploring is to try to use statistical properties of the matrix to identify stable blocks using *a priori* rubrics such as finding subsets of species where all inter-species interactions within the subset are relatively weak. As discussed in Appendix F, this is analogous to the problem of identifying *k*-cliques in a random graph, which is a computationally difficult, NP-hard problem. The problem of identifying all stable states in a given random system is likely at least this hard. As discussed also in [43], the simulation approach, which samples stable states with a given probability distribution *p*_*s*_ is much more effective at computing things like ensemble averages over the set of states with that probability weighting than over the full ensemble with equal weighting, which is computationally much more difficult and may miss low-probability states without exponential effort. The simulation approach that samples with distribution *p*_*s*_ is also more compatible with an experimental approach that would give states with a similar distribution.

**FIG. S26:**
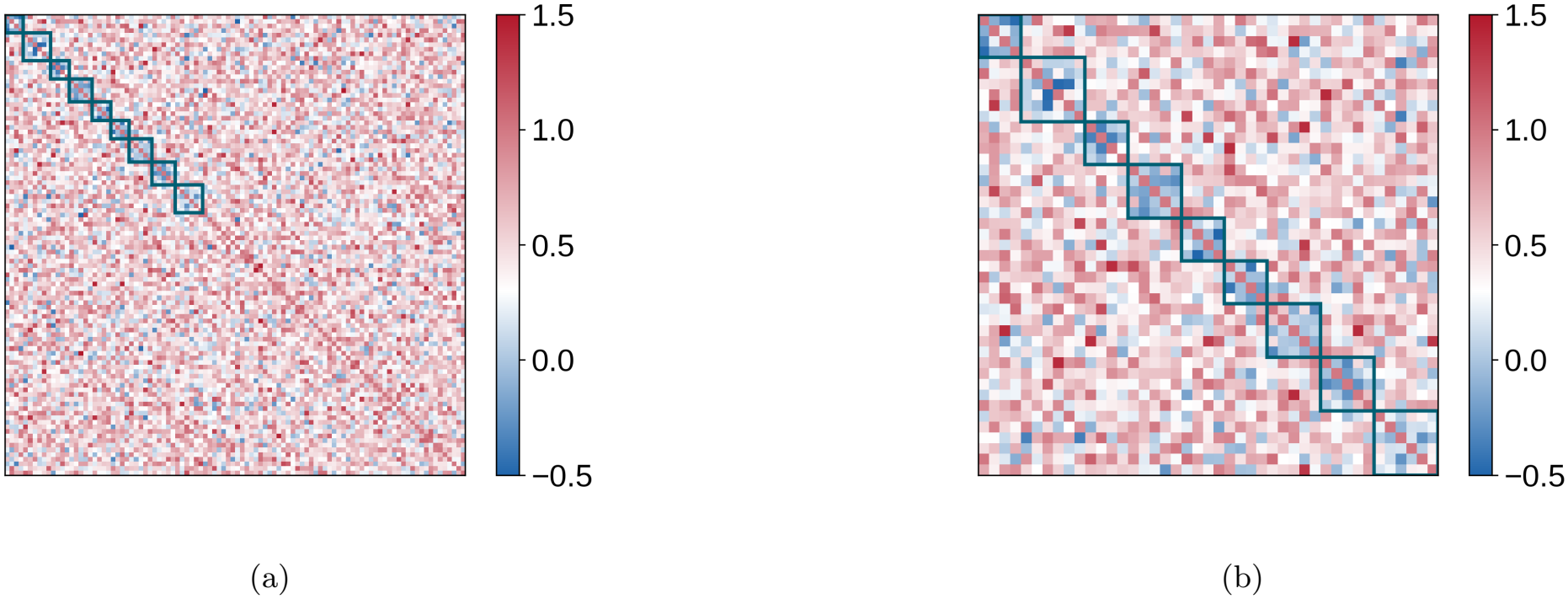
Emergent block structure revealed by selecting non-overlapping states. (a) Full interaction matrix. The chosen non-overlapping states are placed first, followed by the remaining species. (b) Subset of the matrix showing only the chosen states.

We begin by ordering all states in descending order of their likelihood. We then select all species from the most likely state and reorder the matrix to have rows and columns corresponding to them appear first. Then we choose species in the next most likely state. We check whether it shares any species with the previously selected state. If there is even one species that overlaps, we discard this state and move to the next one. If there is no overlap of species between states, we then move the rows and columns corresponding to species in this state right after those belonging to the first state. In this way we iteratively build a submatrix of the full interaction matrix until we have considered all states. This submatrix contains a set of states with no overlaps in species. This submatrix of the full interaction matrix reveals a clear block structure (Fig. S26a–b).

This method emphasizes high-likelihood states and reveals a clear emergent block structure in at least part of the full interaction matrix. However, this emergent block structure appears not just for highly likely states but also those with low likelihood. We demonstrate this by selecting states uniformly randomly rather than following descending order of likelihood as in the previous scheme. That is, we begin by picking a random state regardless of its likelihood and proceed iteratively. This procedure yields different sets of non-overlapping states each time, yet a clear block structure emerges in all cases (Fig. S27). Importantly, we do not alter any species interactions—we only reorganize their ordering within the matrix to reveal the latent structure.

These results show that block-like organization represents a fundamental property of our random ecological matrices, not merely an artifact of selecting high-likelihood states. The consistent emergence of block patterns across different selection schemes validates our theoretical approach of treating states as effective blocks in competition with each other.

**FIG. S27:**
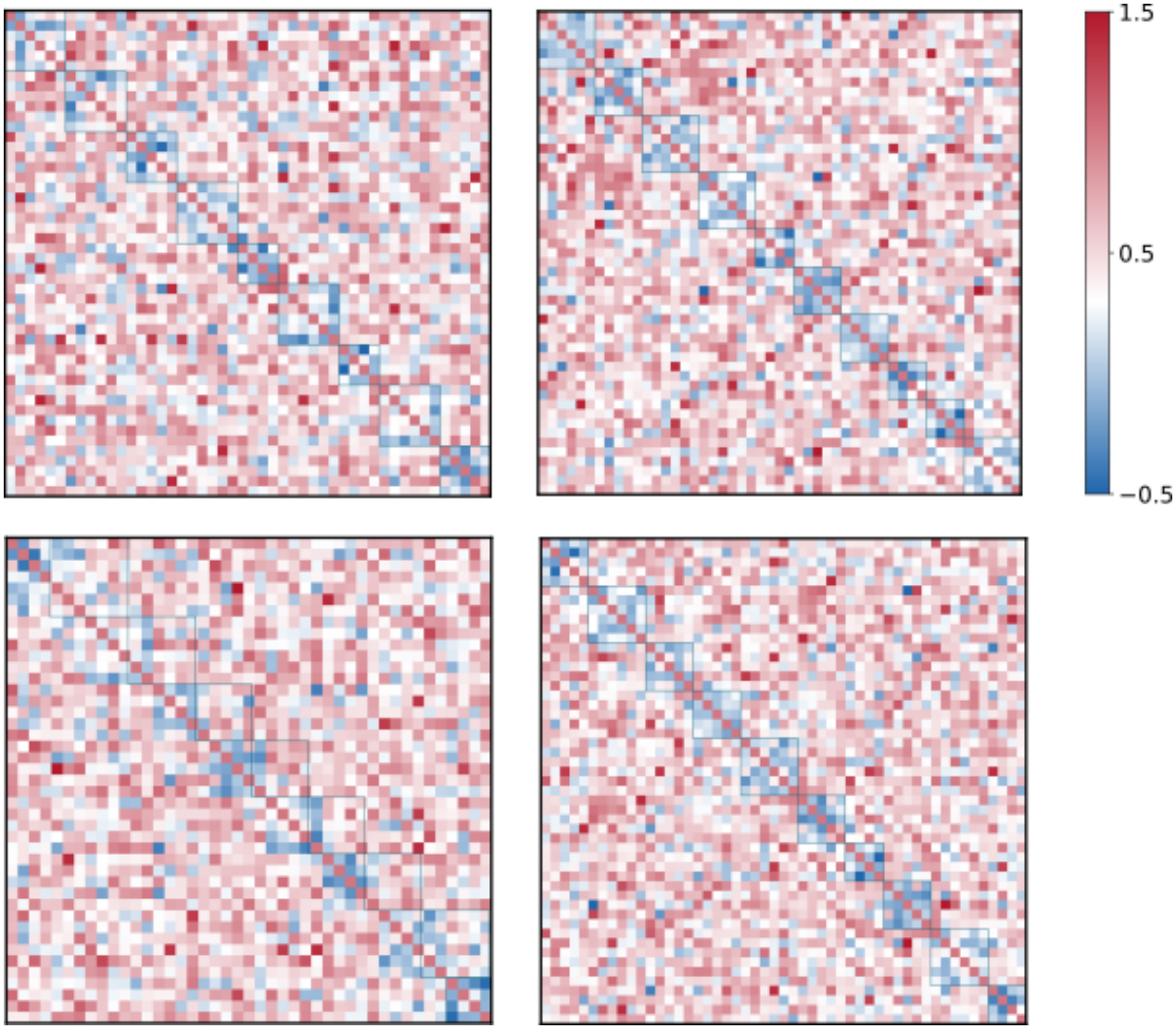
Robustness of block structure to reordering species in different ways. Shown are reordered subsets of the same interaction matrix. Each example shows a different reordered sub-matrix obtained by choosing different non-overlapping sets of states (see text in Appendix G). Although the species and states selected in each submatrix vary, we always observe that a clear block structure emerges when we cluster species from the same state together—at least in the submatrices containing only species found exclusively in one state.

## Appendix H: Accounting for overlaps in the block model

In the block models analyzed in Appendix E, each species belongs to exactly one block. In random disordered matrices, however, states overlap through shared species (Appendix F). Here, we study our block model to systematically vary the extent of overlap between blocks while keeping the number of states fixed. We then use this model to show that when the extent of overlap is sufficiently high, the true block size is a worse predictor of block likelihoods than the mean block size. We also find that the random matrices we studied in the main text indeed have sufficiently large overlap such that using the mean block size for all blocks would serve as a better predictor. This justifies the mean-field approximation of the block model to predict likelihoods in random disordered matrices, which we use in the main text (Fig. 3) and in Appendix I.

To build intuition, consider building an interaction matrix for an ecosystem with 20 blocks or states. Say we want 10 blocks of size 6 and 10 blocks of size 4. The total number of species slots required to fill these blocks is 10*×* 6 + 10 *×*4 = 100. In a perfect block matrix, all 100 slots are filled by 100 distinct species, each species appearing in exactly 1 state. On the other hand, if there are only 50 species in the pool to form the same 20 blocks, each species must participate in 2 states on average. We use this idea to define the *extent of overlap m* as the average number of stable states in which a species participates. Note that an extent of overlap equal to *m* = 1 corresponds to the perfect block structure with no overlaps we studied before. Larger values *m >* 1 introduce progressively more overlap between states.

### Constructing block models with controlled overlap

To verify that our results are robust to the specific way in which we introduce overlaps, we used two different methods to construct overlapping block matrices with a given mean extent of overlap *m*, for a fixed set of blocks (number of stable states) with fixed sizes and biomass values. We systematically increased *m* and studied the prediction error of two block models (as described in Appendix E): one where we used the actual block size of each respective block, and the other where we used the average block size for all blocks.

Both methods rely on the same key idea to introduce overlaps. Namely, if the number of species in the pool is smaller than the total number of species slots Σ _*α*_ *L*_*α*_, where *L*_*α*_ is the size of block *α*, some species must appear in more than one block, since there are not enough distinct species to fill all slots without repetition. The fewer the species in the pool, the more frequently each species must be reused across blocks, and the greater the overlap between states. We thus used a reduced species pool of size Σ _*α*_ *L*_*α*_*/m*, so that on average each species fills *m* slots, giving a mean extent of overlap equal to *m*. In both methods, we assigned within-block interactions in the same way. For species belonging to block *α*, we set the within-block interaction strength to 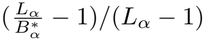, which is weaker than *D*, ensuring that species within the block could coexist with required biomass 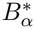 (Eq.(E5)). We set all diagonal elements to 1 and all off-diagonal elements outside any block to a constant *D*. If a pair of species appeared together in more than one block, we set their interaction to the average of the values assigned by those blocks. In the first method, we ensured that all blocks had exactly the same extent of overlap (“constant overlap”), while in the second, we allowed the extent of overlap to vary, while controlling the mean extent of overlap (“variable overlap”).

#### Method 1 (Constant overlap)

We started with a pool of ⌊Σ _*α*_ *L*_*α*_*/m*⌋ species. We went through each block sequentially, and for each block *α*, we randomly picked *L*_*α*_ species from the pool. Once a species was picked *m* times, we removed it from the pool. If Σ _*α*_ *L*_*α*_ is not divisible by *m*, we could not fill the remaining *r* = Σ _*α*_ *L*_*α*_ mod *m* slots from the pool, so we drew species again from the full set of already-used species. In such cases, some species may be used more or fewer than *m* times, but the deviation was negligible and the mean extent of overlap remained effectively equal to *m*.

#### Method 2 (variable overlap)

We filled each block by randomly drawing *L*_*α*_ species from the reduced pool, without any constraint on how many times a species could be selected. Since the pool was smaller than the total number of slots, we necessarily selected species multiple times, causing them to participate in multiple states. This resulted in different species having different extents of overlap, with a mean extent of overlap equal to *m*.

For random matrices, we observe that the extent of overlap of a species does not depend on the diversity of the states it is present in. Therefore, in both methods, we constructed our overlapping block matrices such that the extent of overlap of a species is independent of the block size it participates in.

**FIG. S28:**
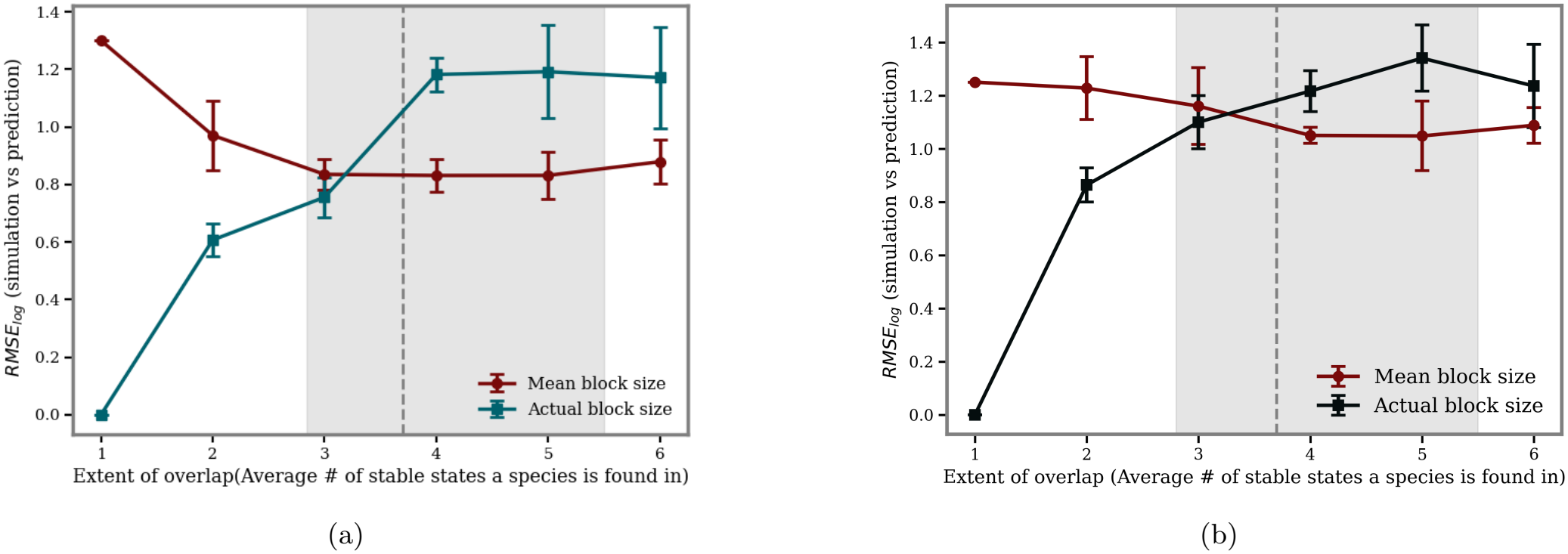
Using mean block size for likelihood prediction gives better results in the presence of overlapping states. As overlap increases, the prediction error using actual block sizes grows, while the error using the mean block size decreases. Beyond an extent of overlap of approximately 3, the mean block size gives better predictions. (a) Fixed overlap: each species participates in a fixed number of blocks. (b) Variable overlap: Different species participates in different number of blocks. The gray shaded region indicates the range of overlap corresponding to the random interaction matrix studied in the main text; the dashed line marks the measured extent of overlap for that matrix. Error bars indicate SEM of multiple realizations of the interaction matrix at fixed overlap structure.

We compare two prediction schemes across varying extents of overlap: using each state’s actual diversity *L*_*α*_, and using the mean diversity *L* = ⟨*L*_*α*_⟩ for all states. At zero overlap (*m* = 1), we find that using the actual block size gives better predictions, as expected from the exact block theory. However, as the extent of overlap increases, the error for the actual-block-size prediction grows, while the error for the mean-block-size prediction decreases. Beyond an extent of overlap of roughly *m* = 3, the mean block size consistently outperforms the actual block size in predictability (Fig. S28). We also measured the extent of overlap metric for the random interaction matrices we studied in the main text and found that they fall within this high-overlap regime (grey regions in Fig. S28).

### A single overlap between blocks increases both their likelihoods

To understand why overlaps change predictions from the block model, here we study how introducing overlaps changes block likelihoods. For this, we first consider the simplest case: a perfect block-structured matrix with no overlaps, into which we introduce a single overlap between exactly two blocks by having them share a single species. We find that the blocks participating in the overlap now have higher likelihood compared to when these blocks did not overlap (Fig. S29). Consequently, the likelihoods of all other non-overlapping blocks decreases a bit to compensate. This is simply because the total likelihood is conserved (Σ_*α*_ *p*_*α*_ = 1). This effect shows that by sharing overlapping species, blocks can effectively boost their likelihood.

**FIG. S29:**
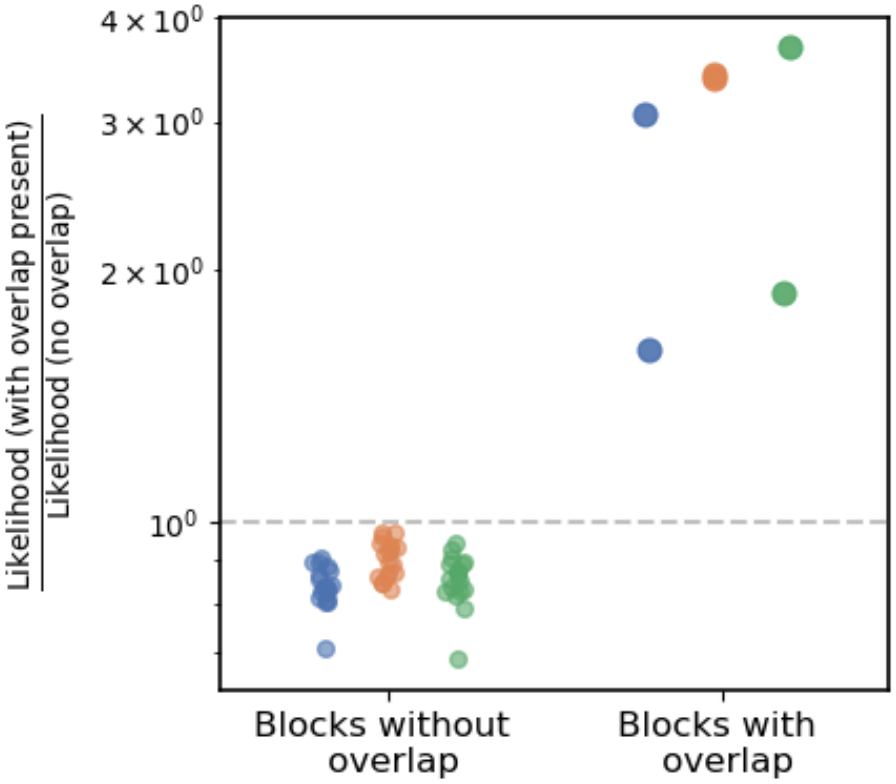
Overlapping blocks experience a mutualistic increase in likelihood. The *y*-axis shows the ratio of likelihood with overlap present to that without overlap. This ratio exceeds 1 for blocks participating in the overlap and is less than 1 for all other blocks. Different colors represent different matrices where two blocks are randomly chosen to overlap.

To gain further insight into this mathematically, we examine the block-level dynamics in the presence of overlap. When blocks *α* and *β* share species, the effective dynamics of block *α* acquires an additional term compared to Eq.(E8):

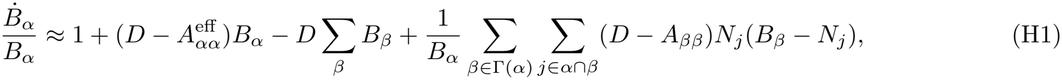

where Γ(*α*) denotes the set of blocks that overlap with block *α*, and the inner sum runs over species *j* shared between blocks *α* and *β*. The last term is the additional contribution arising from overlap. Absorbing it into the effective self-inhibition yields a modified value

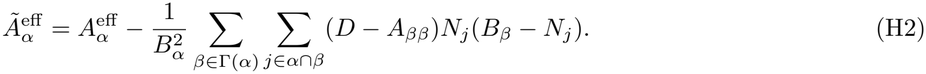

Since *D > A*_*ββ*_ and *B*_*β*_ *> N*_*j*_ for each shared species, the correction term is positive, making 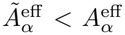. Blocks participating in overlap thus have a reduced effective self-inhibition, which increases their likelihood.

### When overlaps are randomly distributed and widespread, smaller blocks typically benefit from overlap at the expense of bigger blocks

We next consider the case where every block participates in overlap by randomly pairing blocks and having each pair share one species. We find that smaller blocks show a net increase in likelihood, while larger blocks show a net decrease (Fig. S30(a)). This asymmetry can be understood from Eq.(H2). Consider two blocks *α* and *β* with *B*_*β*_ *> B*_*α*_ that share a single species with abundance *N*_*k*_. The reductions in effective self-inhibition are

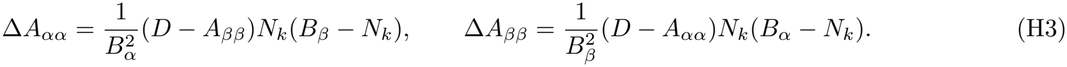

Since *B*_*α*_ *< B*_*β*_, the smaller block has a smaller denominator 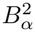 and a larger numerator factor (*B*_*β*_ *N*_*k*_), giving Δ*A*_*αα*_ *>* Δ*A*_*ββ*_: the smaller block experiences a larger reduction in self-inhibition and thus gains more benefit from the overlap. For equal-sized blocks sharing a small overlap (*N*_*k*_ *< B*), the benefit scales as Δ*A*∼ 1*/B*, confirming that smaller blocks gain disproportionately. Combining these results, the benefit from overlap follows the ordering

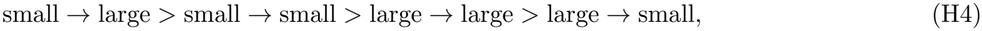

where “small→ large” denotes the benefit received by a small block from overlapping with a large block.

The above explanation is for the case where each block overlaps with one other block irrespective of block size. As discussed earlier, in a random ecosystem, the extent of overlap of a species is independent of block size. Furthermore, we observe that in our random disordered matrices, larger blocks overlap with more blocks compared to smaller blocks, presumably because they contain more species. After including this effect, we still find that smaller blocks become more likely, while larger blocks become less likely (Fig. S30(b)).

**FIG. S30:**
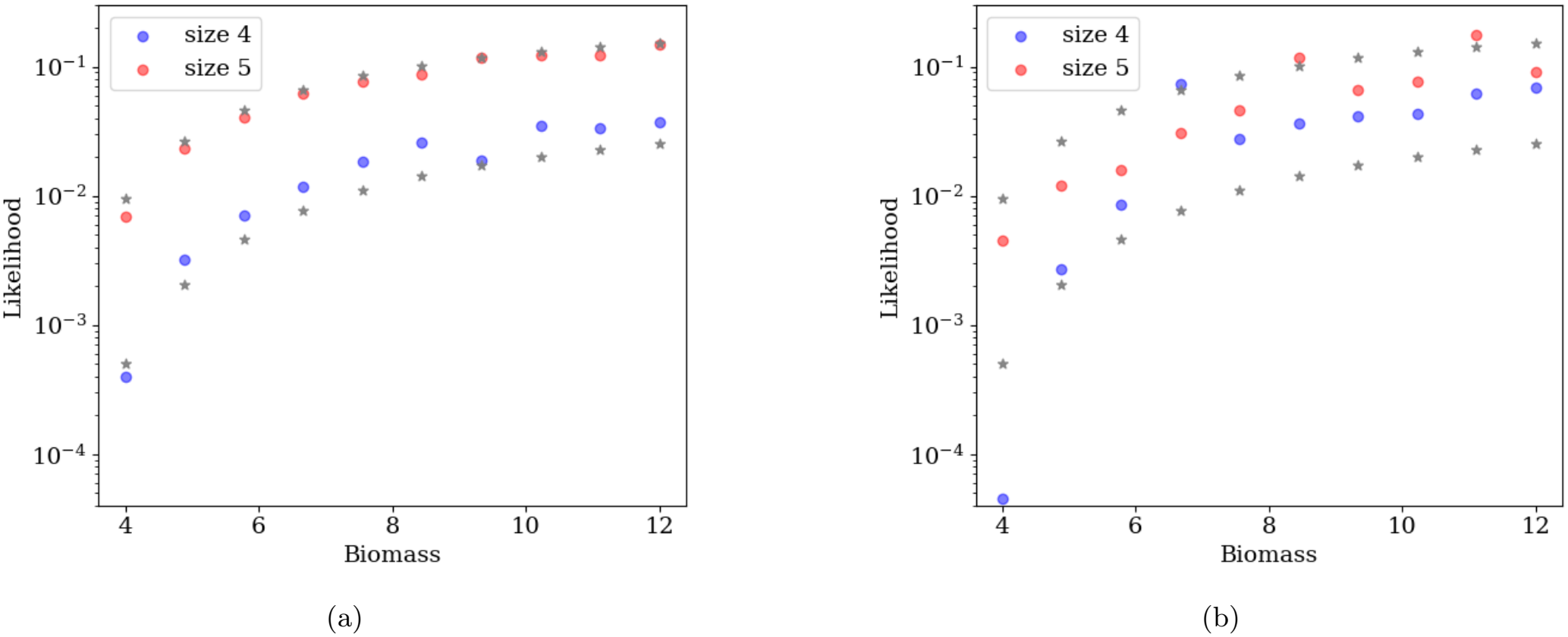
Overlap generally increases the likelihood of smaller blocks and decreases the likelihood of larger blocks. (a) We randomly pair each block with one other block and let them share a single species. Stars and circles represent likelihoods without overlap and with overlap, respectively. For each biomass value, two stars correspond to the two block sizes with the same biomass. The upper star represents the larger block, while the lower star represents the smaller block. For most states, smaller blocks (blue) show an increase in likelihood, while larger blocks (red) show a decrease in likelihood. (b) We pair each block of size 5 with two other blocks and let it share one species with each block, resulting in two distinct shared species, while blocks of size 4 share species with only one block. Even though the larger block shares more species, for most states smaller blocks (blue) show an increase in likelihood, while larger blocks (red) show a decrease in likelihood. In both (a) and (b), we use the same set of blocks, consisting of 10 blocks of size 4 and 10 blocks of size 5, with the same biomass values.

### Connection to the mean-field approximation

The disproportionate benefit to smaller blocks explains why the mean block size produces better predictions than the actual block size in the presence of widespread overlaps, at least for the ways of introducing overlaps that we tried. Such widespread and randomly distributed overlaps (which are equally likely for states roughly independent of their size/diversity) effectively change the likelihoods of small and large blocks closer to each other. In fact such an effect tends to make block likelihoods come closer to the likelihood they would have if they all had the mean size: small blocks are boosted in likelihoods while large blocks are suppressed. Using the actual *L*_*α*_ for each state in the block model therefore misses this redistribution. The mean-field approximation *L*_*α*_ →*L* is a simple and implicit way to capture this effect. This is why, we believe, it produces better predictions despite being a cruder description of the individual states.

## Appendix I: Calculation of likelihood in case of random interaction matrices

We saw in Appendix G that random disordered interspecies interaction matrices exhibit partial block-like organization. In perfectly block-structured matrices, we can calculate likelihood using only the steady-state biomass and block size, as developed in Appendix E. In this appendix, we apply this framework to random matrices, elaborating on the methodology summarized in the main text, as well as considering alternative approaches.

One option is to treat each state as an effective species — like a monodominant model — and use only equilibrium biomass information for predictions, while ignoring block size (diversity). So, we then assume the initial condition of each block to be a uniform distribution, similar to the monodominant model. We find that this approach tends to overpredict strongly for low-likelihood biomass. (Fig. S31a), indicating that biomass alone provides insufficient information. A second option is to incorporate both block size (diversity) and equilibrium biomass information — like the full-block model — using the distinct species diversity *L*_*α*_ of each state. Surprisingly, this approach also produces poor predictions (Fig. S31b). The overlapping nature of blocks in our random matrix (Appendix F) suggests that effective size might differ from the simple species count within each state. Moreover, the cross-state inhibitions are not constant, but have disordered fluctuations.

We find that we achieve most accurate predictions by using a hybrid approach: that of using the true equilibrium biomass of each state along with the average block size across all states (rather than using state-specific diversity values). As shown in Appendix H, as overlap increases, the mean block size becomes a better predictor of the likelihood. The predicted and observed likelihoods match reasonably well over a wide range (Figs. 3b, S32 and S33). Moreover, we find that this predictability persists throughout the entire multistable region of the model (Fig. S34, S35). For a random matrix, this coarse-grained information proves sufficient to estimate state likelihoods effectively. We observe some discrepancies at very low likelihoods for two reasons. First, our number of simulations, while extensive are limited, and do not sample these rare states sufficiently to provide reliable likelihood estimates. Second, we do not explicitly model the detailed effects of overlaps and the exact structure of each state in our mean-field approach. Nevertheless, across the multistable regime for different values of *µ* and *σ*, we successfully predict both individual state likelihoods and the overall biomass–likelihood relationship using only biomass and average species richness (Fig. S32). These results demonstrate that our theoretical framework may capture the essential dynamics of state-state competition even in the presence of structural complexity.

**FIG. S31:**
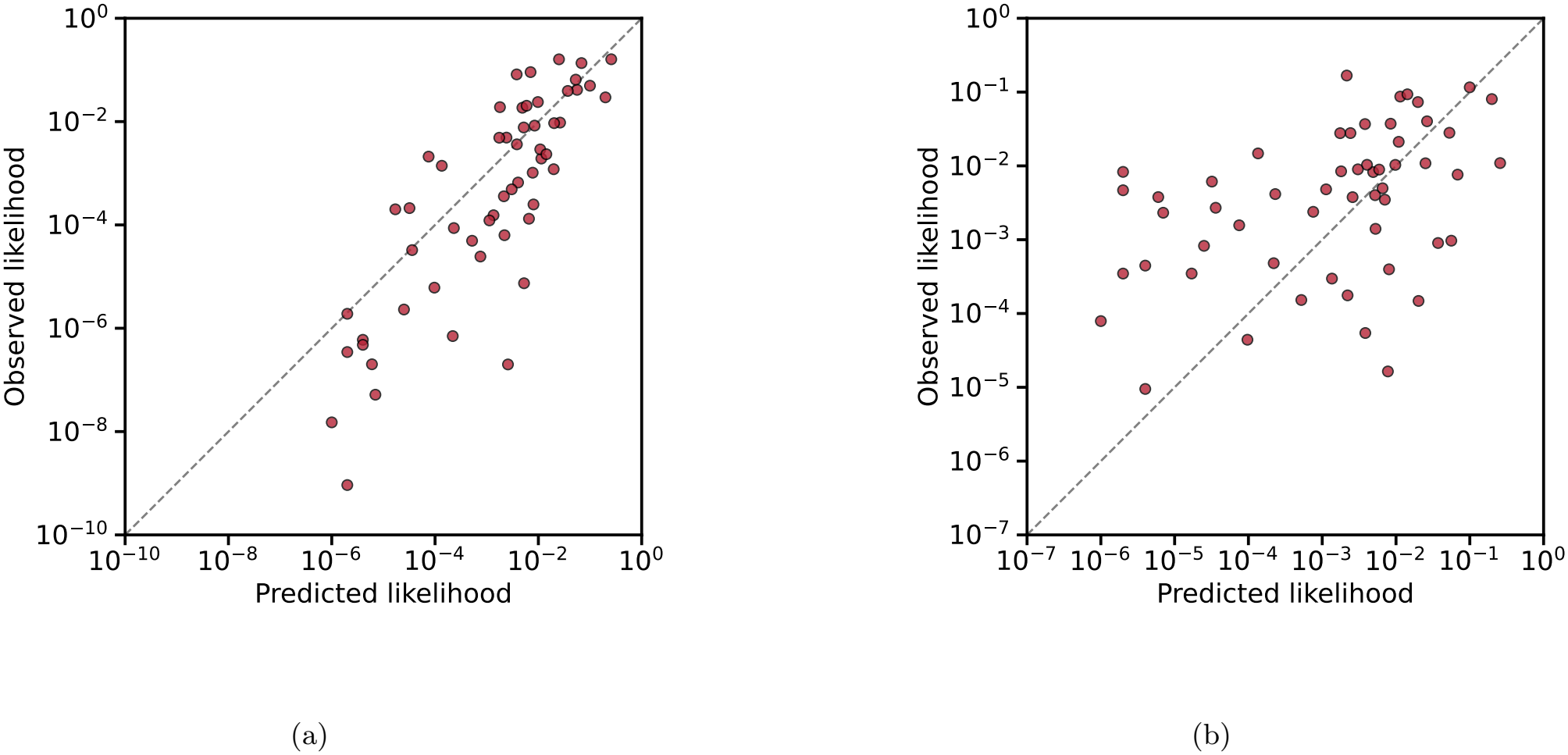
Observed vs. predicted likelihoods for the random matrix used in the main text Figs. 1c and 3, except here the predictions are made using different (less accurate) schemes. In the main text, we predicted likelihoods using the biomass of each state and the average species diversity as a common block size. In (a) we predict likelihoods using only the biomass of each state, setting the diversity of each state to 1, thus effectively using predictions from the monodominant model (Eq. (4) in the main text). In (b) we use the biomass of each state and the diversity of each state (rather than a common diversity equal to the mean). Both schemes produce worse predictions than in Fig. 3b. In panel (a) we systematically overpredict likelihoods across all biomass ranges. Surprisingly we also produce worse predictions for (b), where we used the actual diversity of each state rather than a mean-field approximation. Comparing goodness of fit using *χ*^2^ values, we notice that the method in the main text Fig. 3b produces a *χ*^2^ ≈12, whereas here we get significantly worse *χ*^2^ of (a) 29.15 and (b) 32.50, respectively.

**FIG. S32:**
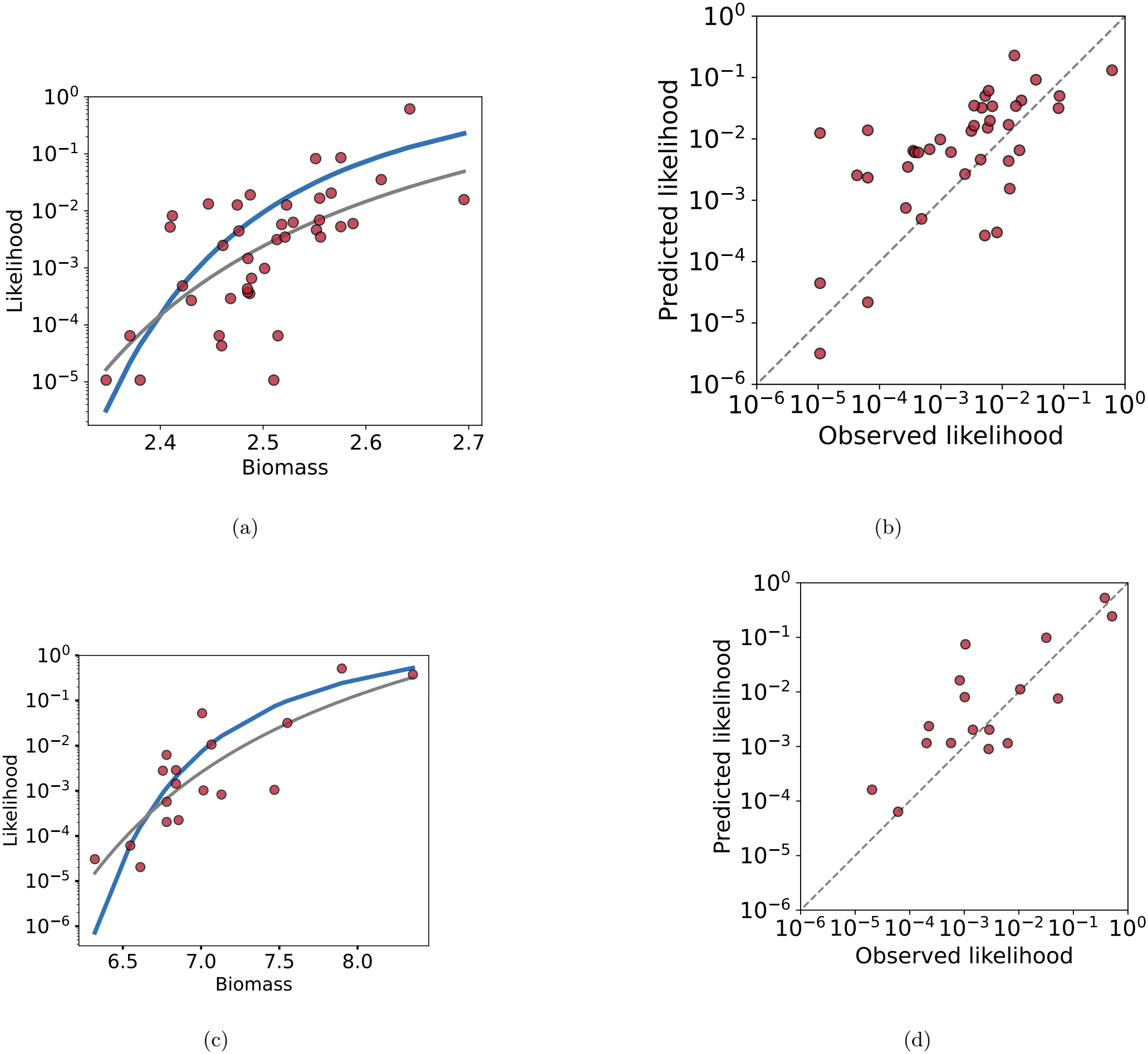
Biomass–likelihood relationship across different parameter regimes. The log-hyperbolic relationship emerges for different values of *µ* and *σ*: panels (a,b) show results for *µ* = 0.5 and *σ* = 0.2, while panels (c,d) show results for *µ* = 0.2 and *σ* = 0.15. We predict likelihoods using biomass and average diversity across all states. In panels (a) and (c), the blue curve shows predictions using Eq. (5) from the main text, while the gray curve shows the hyperbolic fit log(*p*) ∝ (*B*_*c*_ − *B*)^*−*1^. Panels (b) and (d) compare predicted versus observed likelihoods, demonstrating strong agreement across both parameter regimes. In both cases, we note that the predictions are slightly better than the fits in terms of their *χ*^2^ values. In (a)–(b), the fit has *χ*^2^ ≈ 19.05, while the prediction has *χ*^2^ ≈ 15.2. Similarly, in (c)–(d), the fit has *χ*^2^ ≈ 5.2, while the prediction has *χ*^2^ ≈ 4.6

**FIG. S33:**
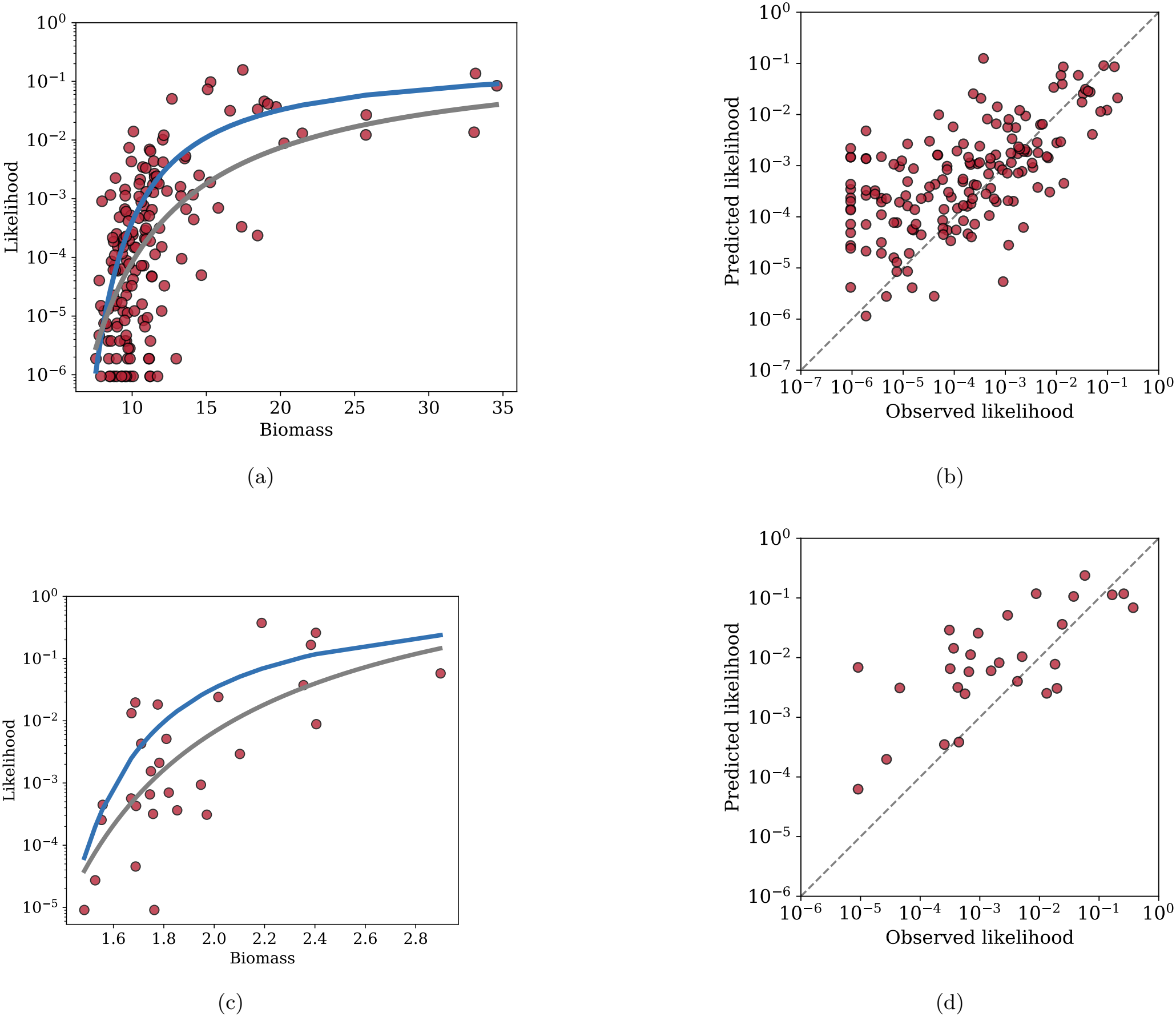
Biomass–likelihood relationship across different number of species. The log-hyperbolic relationship emerges for different values of *S*: panels (a,b) show results for *S* = 200 (*µ* = 0.25 and *σ* = 0.21), while panels (c,d) show results for *S* = 50 *µ* = 1 and *σ* = 0.4. We predict likelihoods using biomass and average diversity across all states. In panels (a) and (c), the blue curve shows predictions using Eq. (5) from the main text, while the gray curve shows the hyperbolic fit log(*p*) ∝ (*B*_*c*_ − *B*)^*−*1^. Panels (b) and (d) compare predicted versus observed likelihoods, demonstrating strong agreement across both parameter regimes. In both cases, we note that the predictions are slightly better than the fits in terms of their *χ*^2^ values. In (a)–(b), the fit has *χ*^2^ ≈ 58.2, while the prediction has *χ*^2^ ≈ 40.1. Similarly, in (c)–(d), the fit has *χ*^2^ ≈ 15.4, while the prediction has *χ*^2^ ≈ 12.2

**FIG. S34:**
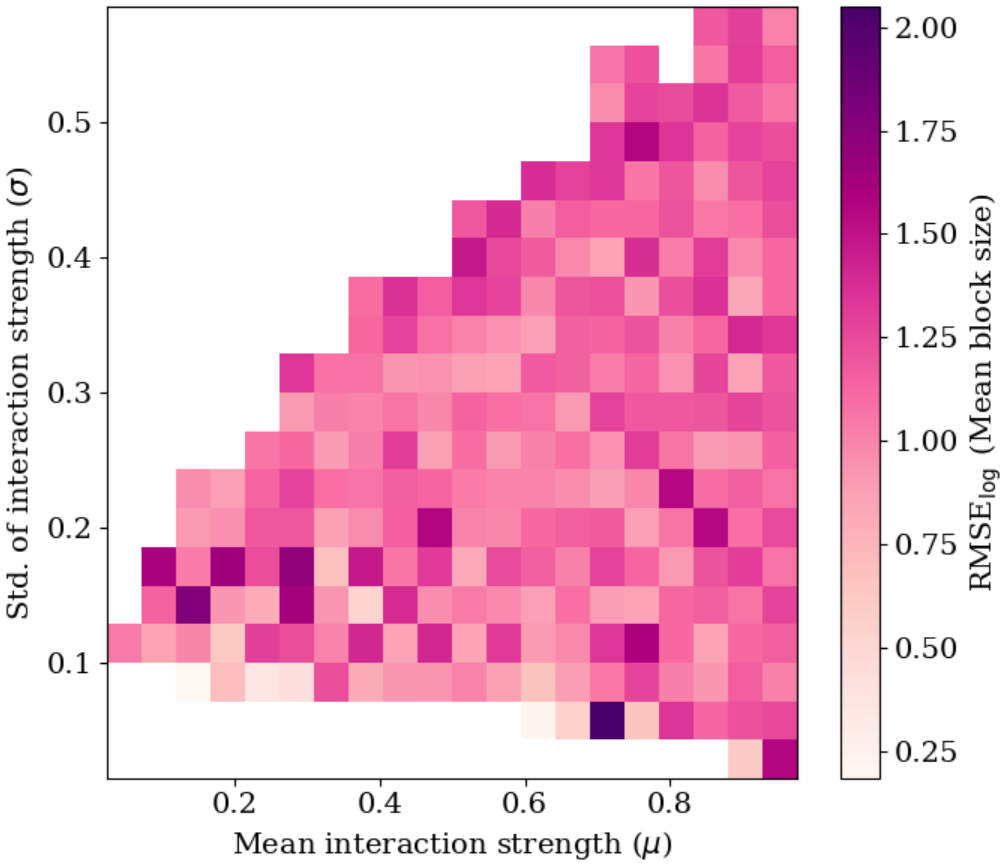
The prediction error of using mean block size in Eq. (5) to estimate likelihood of state remains consistent across the multistable regime. The error is similarly distributed across the parameter space, suggesting that our prediction works equally well throughout the multistability regime.

**FIG. S35:**
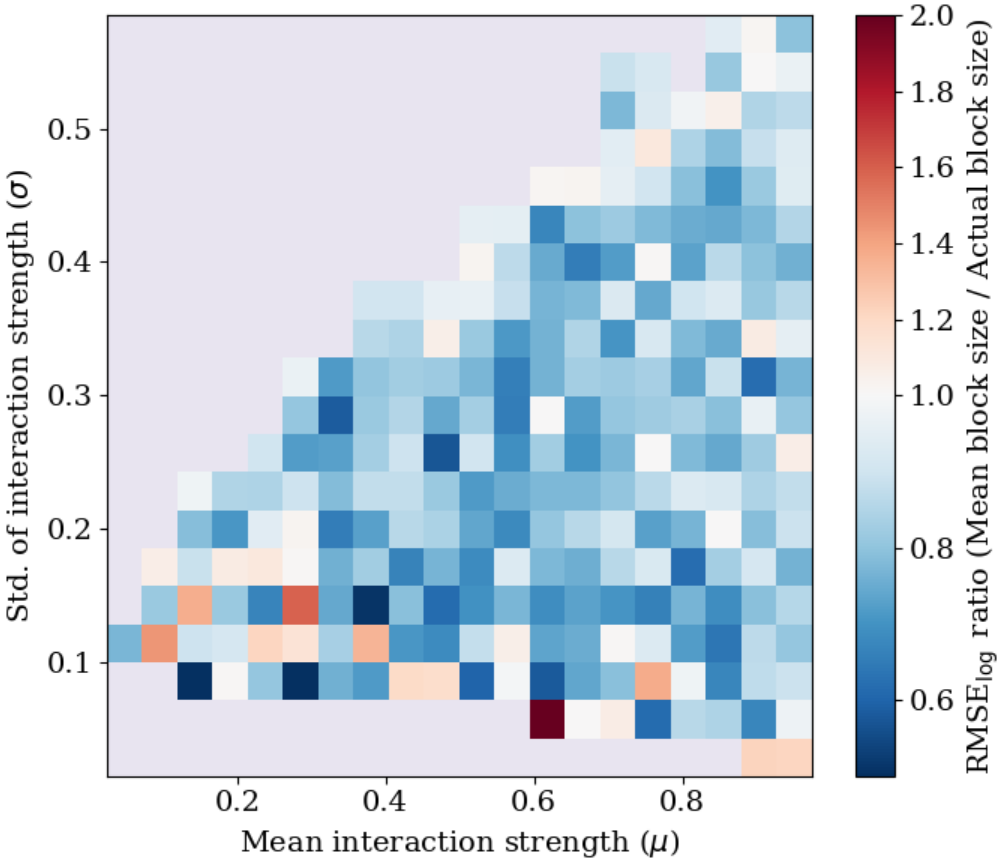
Using the mean block size gives better predictions for likelihood across the multistable regime. The colorbar shows the ratio RMSE_log_(mean block size)*/*RMSE_log_(actual block size). Most values are below 1, indicating that using the mean block size leads to lower prediction error.

## Appendix J: Derivation of the multistability boundary

In the main text, we show that the GLV model with strong interactions exhibits a transition from a single stable state to multiple stable states. This transition occurs at a critical boundary that we derive here analytically and display as the black line in Fig. 1a.

We will follow the approach of previous work [19], which derived this boundary under weak interaction scaling where 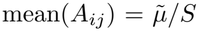 and 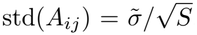. The transition from single to multiple stable states for symmetric interaction matrices occurs at:

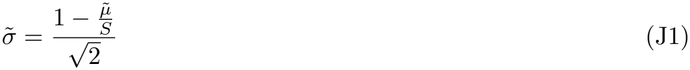

In this manuscript, we consider strong interactions that do not scale with the number of species *S*. To derive the multistability boundary in this regime, we remove the original scaling by substituting 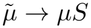 and 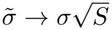. This transformation yields:

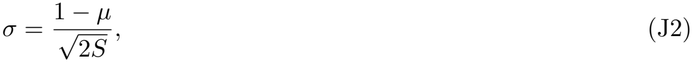

which corresponds to Eq. (2) in the main text. This expression reveals important features of the strong interaction regime. For *µ >* 1, no value of *σ* produces a single stable state—the system always exhibits multistability. In the feasible parameter region, multistability dominates most of the phase space. Our numerical simulations confirm excellent agreement with this analytical prediction (Fig. 1a), validating our approach despite the approximations involved in extending the weak interaction theory to the strong interaction case (in particular, linear response which is central to the original approach is no longer valid, leading one to expect higher-order corrections to be important).

The boundary demonstrates that increasing the mean interaction strength *µ* expands the multistable regime by reducing the critical *σ* required for the transition. This contrasts sharply with weak interaction models where the critical *σ* remains constant [19], highlighting fundamental differences between these two interaction regimes in determining stability.

## Appendix K: The shape of the biomass–likelihood relationship

The relationship between biomass and likelihood exhibits different curvatures depending on the distribution of self-inhibition values across species. In our monodominant framework, diagonal self-inhibition terms *A*_*ii*_ range between 0 and the cross-inhibition strength *D*. Since biomass scales inversely with self-inhibition, large *A*_*ii*_ values produce low-biomass while small *A*_*ii*_ values generate high-biomass.

When self-inhibition values span the full interval (0, *D*), the biomass–likelihood relationship displays two distinct curvature regimes in coordinates where we do not take the logarithm of the likelihood (Fig. S36b). At low-biomass, the curve exhibits concave-upward curvature, while at high-biomass, it becomes concave-downward. Plotting likelihood on a logarithmic scale often masks this transition between curvature regimes.

We can isolate each curvature type by constraining the distribution of self-inhibition values. When all *A*_*ii*_ values cluster near *D*—producing uniformly low-biomass—we observe only concave-upward curvature (Fig. S36d). Conversely, when *A*_*ii*_ values concentrate near zero—generating uniformly high-biomass—the relationship shows concave-downward curvature (Fig. S36f).When self-inhibition values cover the complete range from zero to *D*, both curvature types emerge across different biomass ranges, creating the full biphasic relationship we observe in our simulations (Fig. S36b). In Fig. S36a, due to the log scale used for likelihood, the biphasic behavior is not clearly visible.

The two regimes of the biomass–likelihood relationship can be broken down as a roughly exponential part at low likelihoods (*p* ≲ 1*/S*) and an approach to a plateau where *p* necessarily approaches a constant less than 1 as the biomass gets very large. The hyperbolic form log (*p*) ∝ (*B* − *B*_*c*_)^*−*1^ is in some sense the simplest functional form that approaches a probability of 1 at large biomass and approaches a constant biomass at zero probability. To capture both of these limits, in the main text we thus used the log-hyperbolic relation as a simplified, rough description of the behavior:

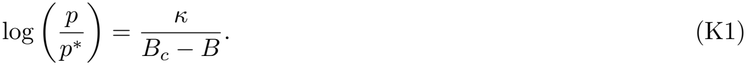

Here, *B*_*c*_ represents the lower biomass threshold. As *B* → *B*_*c*_, the denominator goes to zero and *p* → 0, which captures the sharp drop in likelihood at low-biomass. The critical biomass, *B*_*c*_ decreases with increasing mean interaction strength *µ* and decreasing standard deviation of interaction strength *σ* (Fig. S37).When *µ* decreases, the overall biomass increases, which leads to an increase in *B*_*c*_. Similarly, when *σ* increases, the variation in biomass becomes larger, producing more high-biomass states. This again increases the least likely biomass *B*_*c*_.

**FIG. S36:**
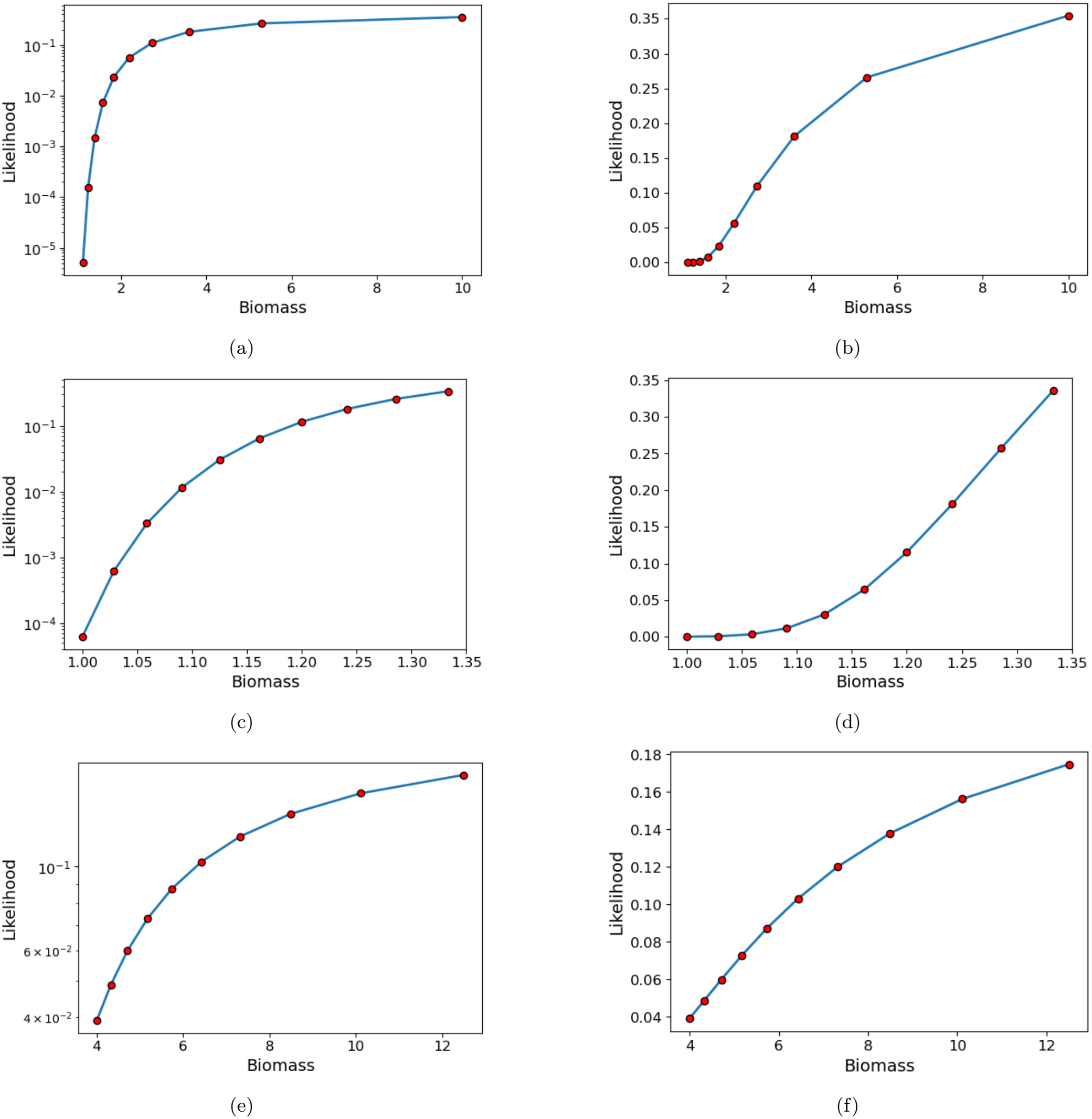
Likelihood versus biomass across different self-inhibition regimes. Note that the pairs of plots on the left and the right in each row represent the same data points, but the left has the *y*-axis on log scale while the right has it on a linear scale. Panels (a,b) show the case where self-inhibition terms span the full range (0, *D*), panels (c,d) show the case where *A*_*ii*_ ≈ 0, and panels (e,f) show the case where self-inhibition values *A*_*ii*_ ≈ *D*. Each pair compares linear y-scale (a,c,e) with logarithmic y-scale (b,d,f). High self-inhibition *A*_*ii*_ ≈ *D* produces a concave-downward biomass–likelihood relationship (f), while low self-inhibition *A*_*ii*_ ≈ 0 generates a concave-upward relationship (d). When self-inhibition values cover the complete range (0, *D*), both curvature types appear across different biomass ranges (f), though the logarithmic y-scale masks this transition (b).

In niche-structured models, it was shown in Ref. [43] that in a limit of large *S* with exponentially many stable states available, there is a precise sense in which probability scales exponentially with biomass. While these results share the common theme that states with greater biomass are associated with larger basins of attraction and higher probability, it is not completely clear exactly how these different functional relations are related. In the models of Ref. [43], however, the exponential/log-linear biomass-probability relationship emerges most clearly in a large *S* limit where the central limit theorem is relevant, corresponding to a limit in which the probabilities of almost all individual states become very low. This is not related in any obvious way to any clear regime of the random matrix systems, but further inquiries in this direction may lead to fruitful further insights.

**FIG. S37:**
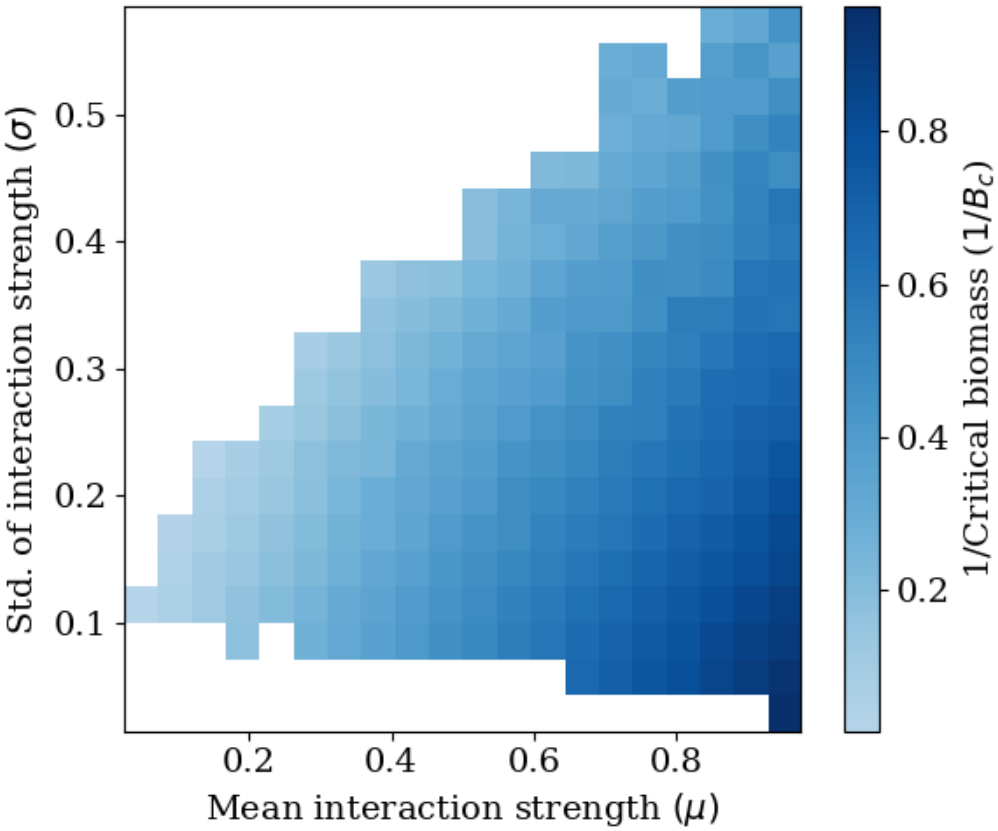
inverse of the critical biomass (1*/B*_*c*_) decreases as mean interaction strtength *µ* decreases and standard deviation of interaction strength *σ* increases across the multistability regime. For each (*µ, σ*) point, we generated a random interaction matrix and fitted log-hyperbolic relation log (*p*) ∝ (*B*_*c*_ − *B*)^*−*1^ to obtain *B*_*c*_. When *µ* decreases, the overall biomass increases, which raises *B*_*c*_ and hence lowers 1*/B*_*c*_. Similarly, increasing *σ* increases the variation in biomass, producing more high-biomass states, which again increases *B*_*c*_ and therefore decreases 1*/B*_*c*_.

**FIG. S38:**
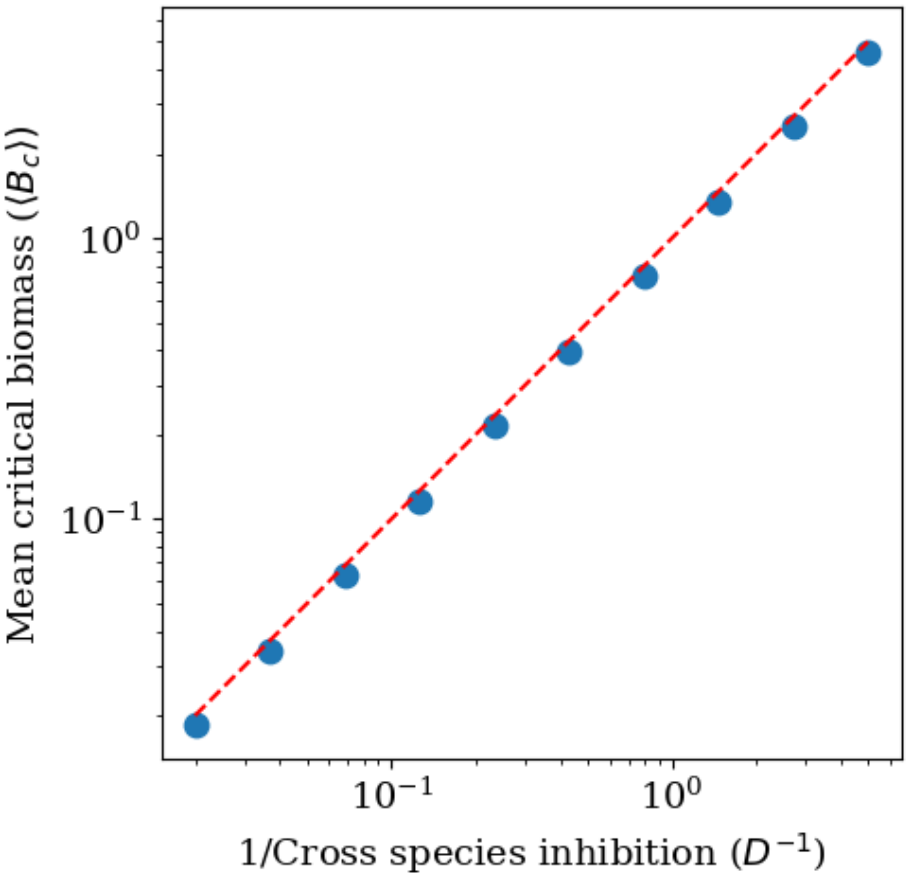
The critical biomass *B*_*c*_ scales as inverse of cross-species inhibition 1*/D* for monodominant model. We generated multiple monodominant interaction matrices with self-inhibition of species uniformly sampled between 0 and *D*, and the cross-species inhibition set to *D*. For each matrix, we computed the likelihoods of the states and averaged over 100 such matrices for each value of *D*. The blue dots show the mean *B*_*c*_ obtained by fitting log-hyperbolic relation log (*p*) ∝ (*B*_*c*_ − *B*)^*−*1^. The mean *B*_*c*_ shows a linear relation with 1*/D*.

## Appendix L: Details of simulation methods for main text figures, including parameters used

We used the simulation method described in Appendix A to estimate the likelihoods for the biomass–likelihood relationship for all results. The code used for the simulations and for generating the figures is available at: https://github.com/eltanin4/emergent-self-inhibition-biomass-likelihood.

**Fig. 1a:** We considered a 35 *×* 35 grid in the phase space of mean interaction strength *µ* (ranging from 0 to 1) and standard deviation of interaction strength *σ* (ranging from 0 to 0.6) . For each point, we generated five symmetric interaction matrices from a normal distribution. For every matrix, we ran simulations with 10^4^ random initial conditions and got the stable states (Appendix A).

If all five matrices led to only one stable state, we colored the point white (single stable state). If, in any of the five matrices, the abundance diverged, we colored the point gray, indicating no stable states. If all five matrices converged and at least one of them showed more than one stable state, we defined the point multistable. We colored these points based on the average number of stable states obtained across the five matrices. We obtained the multistability boundary using the method described in Appendix J.

**Fig. 1c:** The inset shows a specific interaction matrix drawn from the ensemble with mean interaction strength *µ* = 0.5 and standard deviation of interaction strength *σ* = 0.3 from a normal distribution. For this matrix, we computed the biomass and likelihood of stable states using 10^6^ random initial conditions (Appendix A), and plotted them as red points. We then repeated this process for 110 different matrices sampled with the same *µ* and *σ* from a normal distribution.

To obtain the average biomass-likelihood relationship from the resulting data points, we did the following. Starting from the lowest biomass state that we observed, we binned states with similar biomass. Because we observed many more low-biomass states than high-biomass states, we used geometric binning, starting with a bin size of 0.1 and subsequently increasing bin width by a factor of 1.3. Within each bin, we computed the mean log likelihood over all states in that bin log_10_(*p*). This resulted in the average biomass-likelihood relationship across all 110 matrices we sampled, shown as the black curve.

We also fitted biomass–likelihood data for each of the 110 sampled matrices to a log-hyperbolic relation of the form

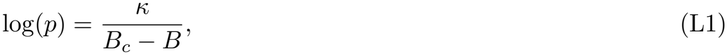

where *B*_*c*_ and *κ* were two fit parameters, which we found to be on average 2 and 16, respectively. We then averaged these fitted curves using the same geometric binning procedure described above, which resulted in the blue curve.

**Fig. 2a:** To check the difference between the interaction of the species which coexist with the ones which don’t, we used the interaction matrix shown in the inset of Fig. 1c and its corresponding stable states. For each species, we divided its *N* ^2^ − *N* cross-species interactions into two groups. In the first group (same state), we included interactions with species that co-occur with it in at least one stable state. In the second group (different state), we included interactions with species that do not co-occur with it in any stable state. We then plotted the distribution of interaction strengths for these two groups: same state (blue) and different state (red).

**Fig. 2b:** This panel shows the same interaction matrix as in the inset of Fig. 1c.

**Fig. 2c:** We constructed an effective coarse-grained model at the level of states for the matrix shown in Fig. 2b. For 55 states, we built a 55 *×* 55 interaction matrix. The diagonal elements are given by 1*/B*_*α*_, where *B*_*α*_ is the biomass of state *α*. We set all off-diagonal elements to a constant value *µ* = 0.5.

**Fig. 2d:** The inset shows an example of a monodominant interaction matrix. We constructed a 10 *×* 10 matrix where the diagonal elements are linearly spaced from 0.1 to 0.3, and all off-diagonal elements are equal to 1.01. In the main plot, we show the biomass and likelihood of all stable states obtained from simulations (red points) using 10^5^ random initial conditions (Appendix A). We also computed the likelihood using *A*_*ii*_, where *A*_*ii*_ is the self-inhibition of species *i*, in Eq. (D) and plotted it as the theory (blue curve).

**Fig. 2e:** To introduce the effect of species diversity on likelihood of the state, we constructed a block-diagonal interaction matrix with block size 4. We set the diagonal elements to 1.01. Within each block, species interacted weakly with a constant interaction strength. For species in different blocks, we set the interaction strength to 1.01. We chose the inter-block self-inhibition values to vary across blocks and linearly spaced them from 0.2 to 0.8.

**Fig. 2f:** To check the prediction accuracy of likelihood for Eq. (5), we constructed a similar matrix for block size 6 with the same in-block and off-block interaction structure (not shown). We computed the likelihood of stable states from simulations for both matrices using 10^5^ random initial conditions (Appendix A). We also estimated the likelihood using biomass and block size in Eq. (5). The plot compares the predicted and observed likelihoods for both the matrices of different block sizes.

**Fig. 3a:** To obtain the average biomass-likelihood relationship for the same 110 matrices used in Fig. 1c, we did the following. We divided the biomass into 15 uniformly spaced bins between 3 and 9. Within each bin, we computed the mean likelihood over all states in that bin. This resulted in the average biomass-likelihood relationship across all 110 matrices we sampled, shown as the red curve. For the same set of states, we also estimated the likelihood using Eq. (5) with the mean block size and biomass. We then averaged the estimated likelihoods using the same binning procedure described above, which resulted in the black curve.

We considered 7 values of the number of sampled matrices, uniformly spaced between 10 and 110. For each case, we sampled that number of matrices and computed the mean observed and predicted likelihoods by binning the biomass into the same bins as mentioned above, which gave 15 paired points 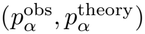 for each case. We then calculated RMSE_log_ (Eq B4) as a measure of prediction error by taking the root mean square of the difference between their logarithms. In the inset, we showed how this error changed with the number of sampled matrices. The error decreased as the number of matrices increased, following a power law. The black line represents a fitted power law with exponent −0.4.

**Fig. 3b:** For the states obtained from the inset matrix shown in Fig. 1c, we used the biomass of the states together with the mean block size in Eq. (5) to estimate the likelihood of the states. We then plotted the observed likelihoods against the predicted likelihood of the states to compare the two.

